# Expanding the catabolic capacity of *Pseudomonas putida* to acetovanillone, 5-carboxyvanillate, and vanillyl glyoxylate for muconate production from kraft lignin-derived aromatics

**DOI:** 10.64898/2026.08.18.745639

**Authors:** Kathryn M. Mains, Dillon T. Hofsommer, Michael A. Gapuz, Prajakta Dongre, Philip S. Zhou, Andrea Salazar, Morgan A. Ingraham, Alexander F. Benson, Kelsey J. Ramirez, Thatcher W. Root, Shannon S. Stahl, Gregg T. Beckham, Allison Z. Werner

## Abstract

The pulp and paper industry produces large volumes of condensed kraft lignin, which is challenging to convert to single chemical products. For this purpose, tandem chemical depolymerization and bioconversion to a single atom-efficient product is a potentially promising strategy. In this study, we conducted copper-catalyzed oxidative depolymerization using pine-derived kraft lignin to generate multiple bioavailable aromatic monomers at a yield of 4.5 weight% (wt%; g monomers per g lignin) from both C– O and C–C bond cleavage, followed by counter-current extraction with a 52 wt% monomer recovery. This resulted in an oxidized lignin product containing vanillin, vanillate, 4-hydroxybenzaldehyde, 4-hydroxybenzoate, 5-formylvanillin, 5-carboxyvanillin, 5-carboxyvanillate, acetovanillone, and vanillyl glyoxylate. Based on this stream composition, we engineered the industrially relevant soil bacterium *Pseudomonas putida* KT2440 to catabolize the latter five compounds via overexpression of ten heterologous genes (*acvABCDEF*_SYK-6_, *vceAB*_SYK-6_, *ligW2*_SYK-6_, and *mdlC*_PP_). We combined these engineered pathways with previously reported strategies for muconate production from G- and H-type monomers to generate *P. putida* KMM428, which utilized 93.6 ± 0.2 mol% of the quantified aromatic monomers in a depolymerized kraft lignin mixture, and produced muconate at a yield of 99 ± 3 mol%, on a quantified monomer basis. Together, this work increases the theoretical carbon conversion efficiency of this process by 37.6 ± 0.1 mol% through incorporation of three β-5 cleavage products, in addition to traditional G-type monomers.

## 1. Introduction

The pulp and paper industry generates abundant sources of lignin that could support the production of biofuels or biochemicals, but this prospect is challenged by the highly condensed kraft lignin resulting from pulping conditions (Argyropoulos et al., 2023; Crestini et al., 2017; Lancefield et al., 2018). The high amount of carbon-carbon (C–C) bonds in kraft lignin are difficult to cleave via conventional lignin depolymerization methods, which primarily focus solely on carbon-oxygen (C–O) bond cleavage, historically limiting the utility of the substrate (Palumbo et al., 2024b). Recent efforts that have focused on C–C bond cleavage in lignin primarily used stabilized lignin oils from reductive catalytic fractionation (RCF) (Gu et al., 2023; Omolabake et al., 2026; Palumbo et al., 2024a; Subbotina et al., 2021), while relatively few studies have targeted C–C linkages in kraft lignin (Dong et al., 2019; Kong et al., 2026; Shuai et al., 2018). Additionally, due to the chemical structure of lignin substrates, depolymerization usually results in heterogeneous mixtures of aromatic monomers and oligomers, making additional processing to valuable chemicals necessary.

Microbial conversion of multiple aromatic monomers through common intermediates into a single product, termed biological funneling, has been demonstrated for diverse heterogeneous feedstocks (Abdelaziz et al., 2016; Becker et al., 2018b; Gómez-Álvarez et al., 2026; Jiménez et al., 2002; Linger et al., 2014; Z. H. Liu et al., 2022; H. Liu et al., 2022; Sodré and Bugg, 2024; Weiland et al., 2022; Werner and Eltis, 2023; Werner et al., 2023; Wu et al., 2023). Converting as much of the lignin-derived carbon into bioavailable aromatics as possible, which requires C–C bond cleavage, is of particular importance for biological funneling (Mains et al., 2026). To this end, we previously demonstrated that copper (Cu)-catalyzed oxidative depolymerization of C–C-linked oligomers from pine- and poplar-derived RCF lignin oils produces bioavailable aromatic monomers that can be biologically converted to muconate or 2-pyrone-4,6-dicarboxylic acid, respectively, by engineered strains of *Pseudomonas putida* KT2440 (Omolabake et al., 2026). From pine lignin, muconate was produced at high molar yields from vanillate and vanillin, but the strain was incapable of utilizing other lignin-derived aromatics including acetovanillone and those generated by β-5 bond cleavage (*i.e.*, 5-carboxyvanillate, 5-carboxyvanillin, and 5-formylvanillin), limiting the utilization of aromatic monomers by 21 wt%.

Metabolism of acetovanillone (Dexter et al., 2022; Higuchi et al., 2022; Hall et al., 2025; Lalande et al., 2026b) and 5-carboxyvanillate (Michener, 2026; Peng et al., 2002, 2005; Vladimirova et al., 2016) has been reported in other aromatic-degrading bacteria, with the latter being a catabolic intermediate for the degradation of lignin-derived dimer 5,5′-dehydrodivanillate (DDVA). Lalande et al. recently engineered conversion of acetovanillone to muconate in *Rhodococcus aromaticivorans RHA1* (*Lalande et al., 2026a*) and Kamada et al. enabled conversion of acetovanillone, 5-carboxyvanillate, and 5-carboxyvanillin to vanillate in *Pseudomonas* sp. NGC7 via heterologous overexpression from plasmids (Kamada et al., 2024). However, these two pathways have yet be evaluated or combined in *P. putida* KT2440.

In this work, we engineered *P. putida* KT2440 to convert an expanded suite of aromatic monomers derived from the oxidative deconstruction of pine-derived kraft lignin to muconic acid (**Fig. 1a**), a well-established precursor for adipic acid (Draths and Frost, 1994; Vardon et al., 2016; Capelli et al., 2019; Kohlstedt et al., 2018) and emerging monomer in performance-advantaged biopolymers (Carraher et al., 2017; Quintens et al., 2019; Carraher et al., 2020; Cywar et al., 2022; Carter et al., 2024; Dardé et al., 2024; Hu et al., 2025; Wu et al., 2025). Expanding on our previous work using Cu-catalyzed oxidative depolymerization (Omolabake et al., 2026), we applied the same catalyst to kraft lignin in a slug-flow reactor followed by counter-current extraction. We quantified the generated aromatic monomers, which included vanillin, vanillate, acetovanillone, 5-carboxyvanillin, 5-formylvanillin, and vanillyl glyoxylate, with smaller amounts of 4-hydroxybenzaldehyde and 4-hydroxybenzoate. We then engineered *P. putida* KT2440 for catabolism of acetovanillone via heterologous chromosomal overexpression of the *acv* and *vce* pathways from *Sphingobium lignivorans* SYK-6. To enable utilization of 5-carboxyvanillate, 5-carboxyvanillin, and 5-formylvanillin, we engineered chromosomal overexpression of *ligW2* from *Sphingobium lignivorans* SYK-6. Vanillyl glyoxylate consumption was then enabled via overexpression of the glyoxylate decarboxylase MdlC. Lastly, building off our previous muconate-producing strain, CJ781 (Kuatsjah et al., 2022), we generated KMM428, a *P. putida* strain that generates muconate from all nine identified and quantified aromatic monomers from deconstructed pine kraft lignin.

**Figure 1.**
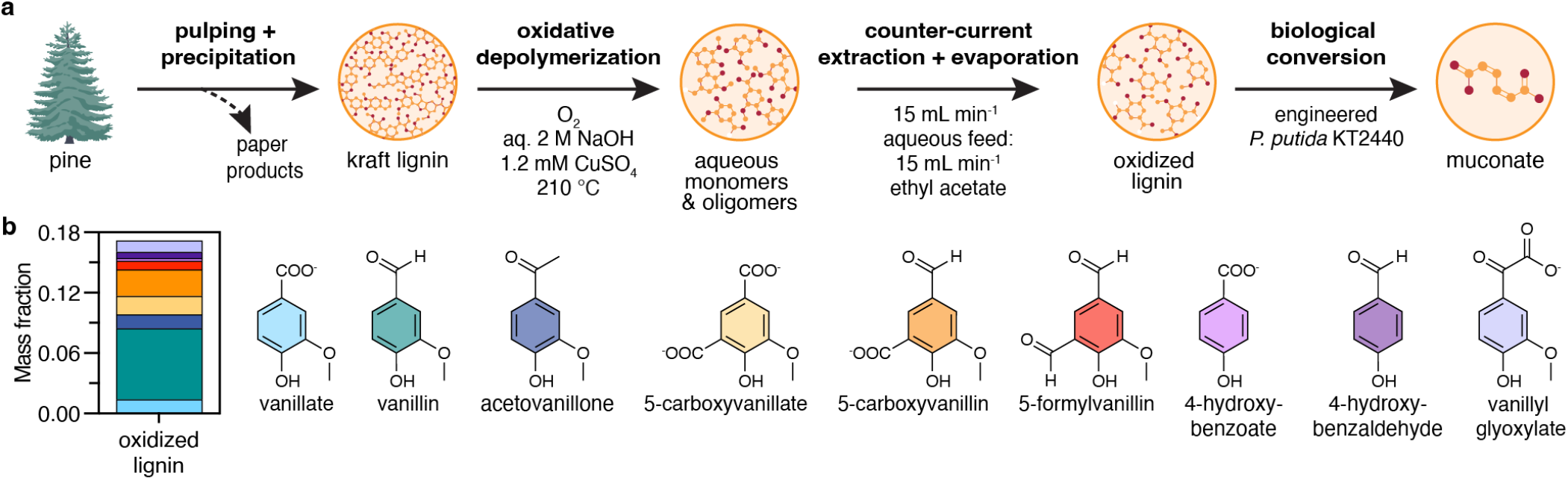
Conversion of aromatic monomers from pine-derived kraft lignin to muconate via engineered *Pseudomonas putida* KT2440. **(a)** Conversion of pine-derived kraft lignin to muconate via Cu-catalyzed oxidative depolymerization, counter-current extraction, rotary evaporation, and biological funneling with engineered *Pseudomonas putida* KT2440. **(b)** Aromatic monomer composition of the oxidized lignin produced from oxidative depolymerization of kraft lignin followed by counter-current extraction, rotary evaporation, and solubilization in water on a mass monomer per mass oxidized lignin basis. Quantified monomers include (from bottom to top) vanillate, vanillin, acetovanillone, 5-carboxyvanillate, 5-carboxyvanillin, 5-formylvanillin, 4-hydroxybenzoate, 4-hydroxybenzaldehyde, and vanillyl glyoxylate. Data in this figure panel are provided in **Data S1**.

## 2. Results

### 2.1. Generation of aromatic monomers from pine-derived kraft lignin

We recently reported the ability of Cu-catalyzed oxidation to cleave C–C bonds in lignin-derived oligomers isolated from RCF of poplar and pine, which produced the typical monomers from oxidative cleavage of pine lignin, namely vanillate, vanillin, and acetovanillone (Li et al., 2024; Luo et al., 2021; Schutyser et al., 2018; Weeda et al., 2024), as well as 5-carboxyvanillin and 5-formylvanillin, which are products of β-5 bond cleavage or the cleavage of other condensed linkages at the 5-position in kraft lignin (Omolabake et al., 2026). Motivated by the success of this oxidation technique to generate bio-available monomers from C–C bond cleavage of RCF oligomers, here we applied Cu-catalyzed oxidation to softwood kraft lignin using a slug-flow reactor to enable continuous oxidation (see section 4.2 and 4.3 for experimental details).

Oxidation of the polymeric lignin led to a dramatic decrease in molecular weight to monomeric products and low molecular weight oligomers as determined by gel permeation chromatography (GPC) and ultra-performance liquid chromatography (UPLC) (**Fig. S1, S2**). The conditions used in this study resulted in a monomer yield of 4.5 wt% (g monomers per g kraft lignin basis; **Table S1**). After acidification, counter-current extraction in ethyl acetate to remove salt, and rotary evaporation to remove the solvent, we obtained an oxidized lignin mixture with a composition of 17 wt% (g monomers per g oxidized lignin; **Fig. 1b**). The quantified lignin-derived aromatics include vanillin (45 mol% on a mol monomer per ∑mol monomer basis), vanillate (8 mol%), acetovanillone (8 mol%), 5-carboxyvanillin (13 mol%), 5-carboxyvanillate (8 mol%), 5-formylvanillin (5 mol%), 4-hydroxybenzaldehyde (5 mol%), and 4-hydroxybenzoate (2 mol%). We also identified vanillyl glyoxylate as a monomeric product, a compound we did not quantify in previous studies, at 5 mol% of the monomer fraction. Given the high proportion of aromatic monomers present in the oxidized lignin that previously engineered strains of *P. putida* KT2440 are incapable of converting to an atom-efficient product (Beckham et al., 2016), we set out to increase carbon conversion efficiency of *P. putida* KT2440 via metabolic engineering.

### 2.2. Heterologous expression of acv and vce pathways enables utilization of acetovanillone in the presence of a supplemental carbon and energy source

We first sought to engineer utilization of acetovanillone, a reported substrate for multiple bacteria (Dexter et al., 2022; Hall et al., 2025; Higuchi et al., 2022; Lalande et al., 2026b) but not *P. putida* KT2440 (**Fig. S3**). Enabling acetovanillone utilization by *P. putida* KT2440 is of particular importance to create a robust bioprocess as acetovanillone negatively impacts cellular growth at concentrations of 5 mM and above (**Fig. S4**). Lalande *et al*. recently compared three catabolic pathways for acetovanillone catabolism from *Sphingobium lignivorans* SYK-6, *Rhodococcus rhodochrous* GD02, and *Actinomadura macra* NBRC-14102 in engineered *Rhodococcus aromaticivorans* RHA1 (formerly known as *Rhodococcus jostii* RHA1) (Lalande et al., 2026b). The authors found that catabolic genes from all three organisms enabled the conversion of acetovanillone to expected pathway intermediates in RHA1, with the *hpe* genes from GD02 enabling the fastest conversion. The pathway from SYK-6, however, is the only pathway shown to have activity on the S-type analog, acetosyringone.

While not present in the pine-derived kraft lignin streams, acetosyringone is a common product in oxidized lignin streams derived from poplar (Omolabake et al., 2026). Thus, we selected the acetovanillone/acetosyringone pathway from SYK-6 (**Fig. 2a**), which includes the genes *acvABCDEF* and *vceAB*, for our engineering efforts to enable flexibility for both G- and S-type lignin streams. Together, these genes convert acetovanillone and acetosyringone to vanillate and syringate, respectively: *acvAB* encodes a 4-acetyl-2-methoxyphenylphosphate (AVP)/4-acetyl-2,6-dimethoxyphenylphosphate (ASP) synthetase, *acvCDE* encodes a biotin-dependent carboxylase, *acvF* encodes an AVP/ASP phosphatase, *vceA* encodes an acetyl coenzyme A (acetyl-CoA)-dependent vanilloyl acetic acid (VAA)/3-(4-hydroxy-3,5-dimethoxyphenyl)-3-oxopropanoic acid (SAA)-converting enzyme, and *vceB* encodes a vanilloyl-CoA/syringoyl-CoA thioesterase. We split the eight SYK-6 genes (Higuchi et al., 2022) across three synthetic operons (*fpva*::P*_tac_*:*vceA*_SYK-6_:*vceB*_SYK-6_:*acvF*_SYK-6_, *PP_5042*::P*_tac_*:*acvAB*_SYK-6_, and *PP_5322*::P*_tac_*:*acvCDE*_SYK-6_), each driven by the strong and constitutive *tac* promoter (P*_tac_*). Each gene was codon-optimized for *P. putida* (**Table S5**) and placed behind a strong synthetic ribosome binding site (RBS) (Cetnar and Salis, 2021; Reis and Salis, 2020). Synthetic operons were integrated into the wild-type *P. putida* KT2440 chromosome via markerless homologous recombination (Johnson and Beckham, 2015), generating strain KMM022 (**Table 1**, **Table S3**).

**Figure 2.**
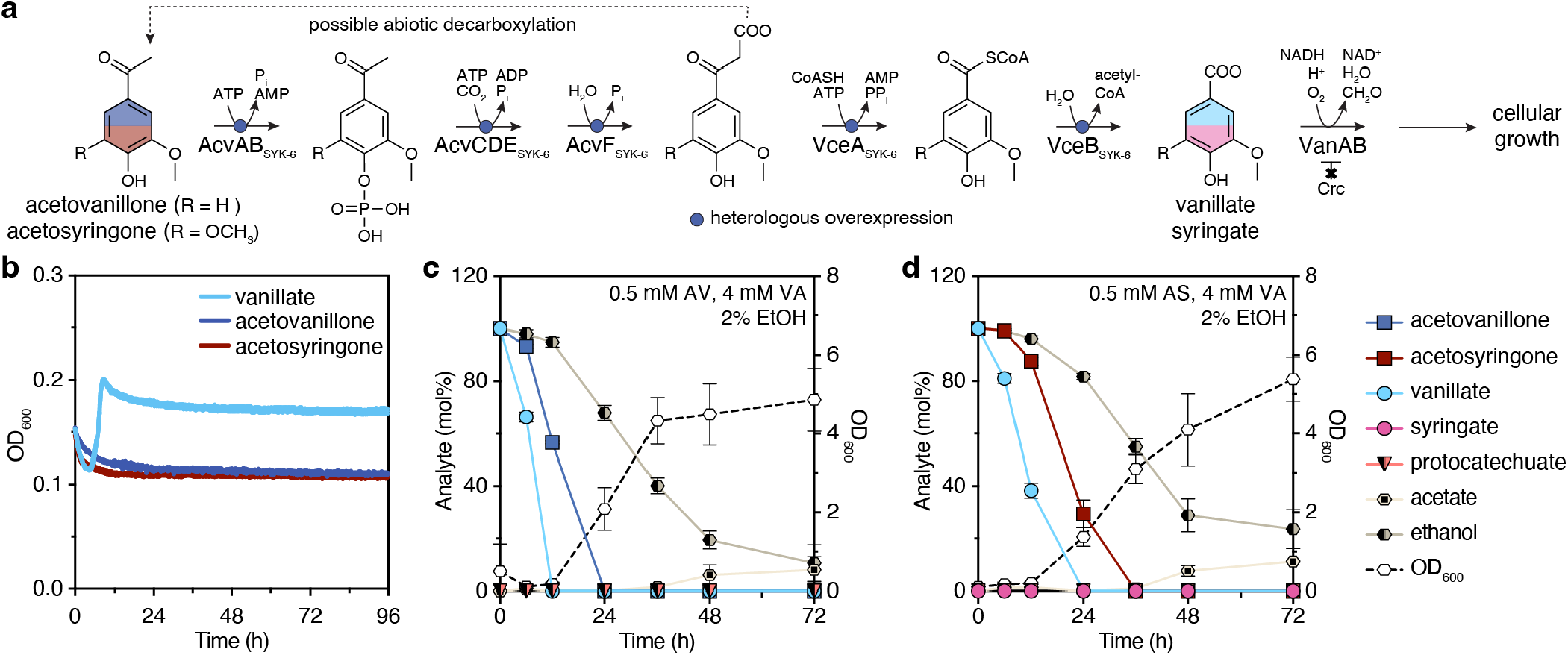
Overexpression of *acvAB*, *acvCDE*, *acvF, vceA,* and *vceB* from *Sphingobium lignivorans* SYK-6 in *P. putida* KMM022 enables utilization of acetovanillone and acetosyringone in the presence of a supporting carbon and energy source. **(a)** Heterologous metabolic pathway for the conversion of acetovanillone to vanillate in engineered *Pseudomonas putida* KMM022. The subscript SYK-6 indicates the protein originated from *Sphingobium lignivorans* SYK-6. Genotype and strain construction details are provided in **Table 1 and Tables S2-S5**. Enzyme abbreviations: AcvAB, AVP/ASP synthetase; AcvF, AVP/ASP phosphatase; AcvCDE, biotin-dependent carboxylase; VceA, acetyl coenzyme A (acetyl-CoA)-dependent VAA/SAA-converting enzyme; VceB, vanilloyl-CoA/syringoyl-CoA thioesterase. **(b)** Engineered strain KMM022 was cultivated in M9 media containing 5 mM vanillate, acetovanillone, or acetosyringone as the sole carbon and energy source in a microtiter plate. Cell growth was determined by optical density at 600 nm (OD_600_). KMM022 was cultivated in M9 media containing 4 mM VA, 2% (v/v) ethanol (EtOH), and 0.5 mM **(c)** acetovanillone or **(d)** acetosyringone in 125-mL shake flasks (25 mL working volume). Time profiles with molar substrate concentrations are provided in **Fig. S5**. For panels (b)-(d), the data points represent the average of three biological replicates, and the error bars represent the standard deviation. All data shown graphically in this figure are provided in **Data S1**.

**Table 1.**
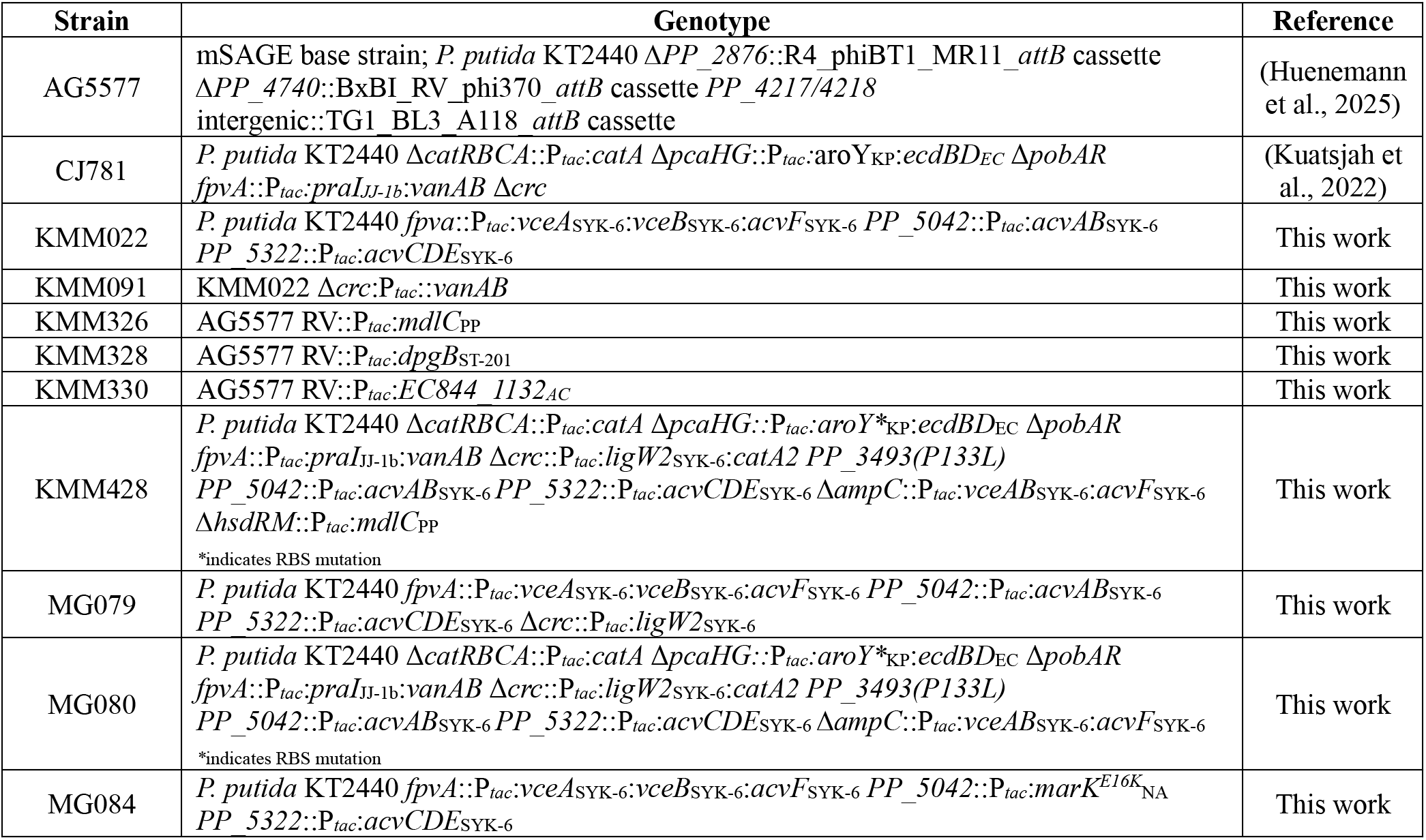
Bacterial strains utilized in this study. Key bacterial strains utilized in this study listed in alphabetical order. Subscripts indicate a protein of heterologous origin, where SYK-6 indicates *Sphingobium lignivorans* SYK-6, NA stands for *Novosphingobium aromaticivorans* DSM12444, PP represents *Pseudomonas putida*, ST-201 indicates *Pseudomonas stutzeri* ST-201, AC represents *Acinetobacter calcoaceticus*, JJ-1b is *Paenibacillus* sp. JJ-1b, EC indicates *Enterobacter cloacae*, and KP represents *Klebsiella pneumoniae*. *RV* indicates an *attB* site for the serine-recombinase from *Mycobacterium* prophage ɸRv1 (Elmore et al., 2023; Smith, 2015). Genotypes and strain construction details are provided in **Tables S2-S5**.

When cultivated in M9 minimal media with 5 mM acetovanillone or acetosyringone (solubilized in dimethyl sulfoxide [DMSO], which *P. putida* KT2440 cannot metabolize), KMM022 did not grow on either substrate as the sole carbon and energy source in microtiter plates despite being able to grow on vanillate as expected (**Fig. 2b**). However, co-feeding KMM022 with 0.5 mM acetovanillone or acetosyringone and 4 mM vanillate solubilized with ethanol (2% v/v final concentration) enabled full utilization within 48 h in shake flasks (**Fig. 2c-d**). We increased the acetovanillone and acetosyringone concentration to 5 mM and 2.5 mM, respectively, to characterize the strain at higher substrate loading. Here, after 72 h, the strain utilized 2.1 ± 0.1 mM acetovanillone and 1.2 ± 0.1 mM acetosyringone alongside 5 mM vanillate and 2% (v/v) ethanol (**Fig. S5**), suggesting further strain improvement was warranted.

It has previously been observed that strong, constitutive expression of *vanAB*, which encodes a Rieske non-heme iron monooxygenase (Erickson et al., 2022; Notonier et al., 2021), and deletion of the catabolite repressor control gene *crc* improved growth on vanillate and syringate (Morales et al., 2004; Johnson et al., 2017). We thus hypothesized that *vanAB* overexpression and *crc* deletion would improve acetovanillone and acetosyringone utilization, and to test this, we generated strain KMM091 (KMM022 *Δcrc:P_tac_:vanAB*). KMM091 did not grow on acetovanillone or acetosyringone as the sole carbon source (**Fig. S6a**). Hall *et al*. recently reported that a mutation to the multiple aromatic kinase, MarK, from *Novosphingobium aromaticivorans* DSM12444 enabled growth of the strain on acetovanillone as the sole growth substrate (Hall et al., 2025). To evaluate if this enzyme would similarly improve acetovanillone utilization in *P. putida* KT2440, we replaced *acvAB* with *marK^E16K^* (strain MG084), but this modification also did not enable growth on acetovanillone as the sole carbon and energy source (**Fig. S6b**).

The requirement for an additional carbon and energy source could suggest either that acetovanillone and acetosyringone do not sufficiently activate the pathway or that additional energy is required. Notably, the reported catabolic pathway for acetovanillone and acetosyringone requires 3 mol of ATP per mol of acetovanillone or acetosyringone, and the second ATP-requiring step could unproductively cycle due to abiotic decarboxylation (**Fig. 2a**). To evaluate these possibilities, we provided KMM022 with 5 mM acetovanillone and combinations of glucose, ethanol, or vanillate equating to the same moles of total carbon (**Fig. S7**). Interestingly, the identity of the additional carbon source did not play a major role, consistent with an energy or redox limitation to convert acetovanillone to vanillate. Since acetovanillone is only a small fraction of monomers present in pine-derived oxidatively depolymerized lignin streams (<10 mol%), and we intended to feed glucose as a growth substrate for muconate production, we continued our engineering efforts by adding to KMM022.

### 2.3. Overexpression of ligW2 enables utilization of 5-carboxyvanillate, 5-carboxyvanillin, and 5-formylvanillin

We next aimed to expand the metabolism of *P. putida* KT2440 to include 5-carboxyvanillate, 5-carboxyvanillin, and 5-formylvanillin (**Fig. S3**). While 5-carboxyvanillate and its aldehyde derivatives have not historically been the focus of biological lignin valorization studies, 5-carboxyvanillate is a known catabolic intermediate for the degradation of 5,5′-dehydrodivanillate. Because of this, at least two 5-carboxyvanillate decarboxylases have been identified, LigW and LigW2, from multiple organisms, including SYK-6, *Novosphingobium aromaticivorans* F199, and *Novosphingobium rhizosphaerae* LY (Peng et al., 2002, 2005; Vladimirova et al., 2016; Michener, 2026). In SYK-6, LigW2 appears to play a more substantial role than LigW in metabolism of DDVA to vanillate (Bleem et al., 2023), and has previously been shown to be active in a *P. putida* strain (Peng et al., 2005).

Thus, to engineer utilization of 5-carboxyvanillate, we added a synthetic operon to express *ligW2* under the *tac* promoter at the *crc* locus, generating strain MG079 (KMM022 Δ*crc*::P_tac_:*ligW2*) (**Table 1**, **Fig. 3a**). 5-Formylvanillin, which requires two oxidation reactions for conversion to 5-carboxyvanillate, was catabolized at low levels by MG079, namely only 21 ± 2 mol% of 5-formylvanillin provided was converted by MG079 in shake flasks over 72 h (**Fig. 3b**). MG079 grew on both 5-carboxyvanillin and 5-carboxyvanillate as the sole carbon and energy source (**Fig. 3c-d**). In vanillin metabolism, *P. putida* KT2440 natively harbors multiple, redundant aldehyde dehydrogenases, including Vdh, that mediate oxidation to vanillate, and we posit that the same is true for conversion of 5-formylvanillin and 5-carboxyvanillin to 5-carboxyvanillate (Graf and Altenbuchner, 2014; Simon et al., 2014; Ruhl et al., 2025).

**Figure 3.**
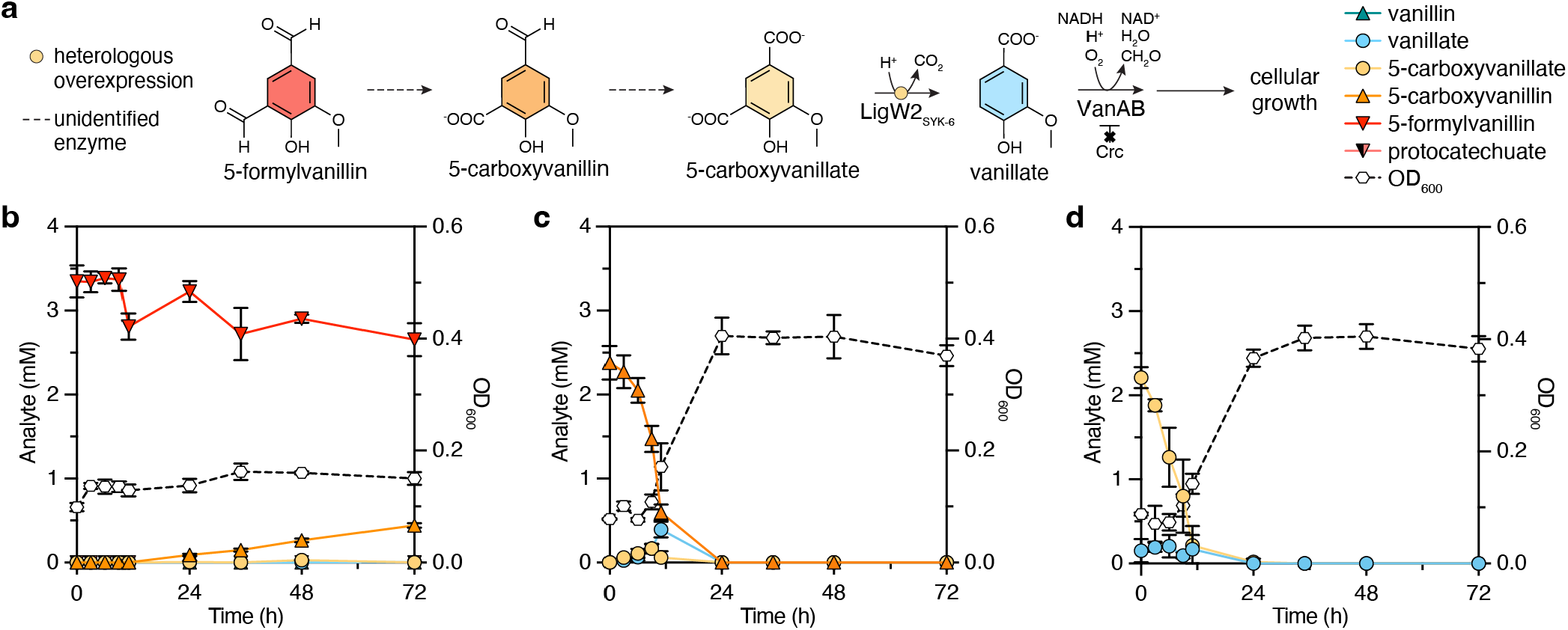
Overexpression of *ligW2* enables robust growth of *P. putida* MG079 on 5-carboxyvanillate and 5-carboxyvanillin, but only minor growth on 5-formylvanillin, as the sole carbon and energy source. **(a)** *ligW2* from SYK-6 was added to KMM022 to generate strain MG079, enabling the conversion of 5-carboxyvanillate to vanillate. Notably, no additional enzymes were added to convert 5-formylvanillin and 5-carboxyvanillin to 5-carboxyvanillate. **(b)-(d)** Growth (OD_600_) and metabolite concentrations for MG079 shake flask cultivations in M9 media supplemented with **(b)** 5-formylvanillin, **(c)** 5-carboxyvanillin, or **(d)** 5-carboxyvanillate as the sole carbon and energy source. For panels (b)-(d), the data points represent the average of three biological replicates, and the error bars represent the standard deviation. All data shown graphically in this figure are provided in **Data S1**.

To evaluate whether the relatively low conversion of 5-formylvanillin in MG079 was sufficient for full conversion in pine-derived oxidatively deconstructed lignin streams where it represents ∼5 mol% of the lignin-derived aromatics, we provided MG079 5 mM of an aromatic mixture containing approximately 20 mol% 5-formylvanillin, which exceeded the amount expected in the oxidized lignin. When the substrate was provided in a mixture, MG079 consumed all 5-formylvanillin within 36 h (**Fig. S8**), demonstrating that *ligW2* overexpression, combined with native aldehyde dehydrogenase activity, was sufficient for the conversion of 5-formylvanillin, 5-carboxyvanillin, and 5-carboxyvanillate to vanillate in *P. putida* KT2440.

### 2.4. Vanillyl glyoxylate consumed by strains expressing phenyl glyoxylate decarboxylases in the presence of vanillate

Next, we were interested in enabling the conversion of vanillyl glyoxylate, and thus we investigated the ability of three thiamine pyrophosphate (TPP)-dependent phenyl glyoxylate decarboxylases to convert vanillyl glyoxylate to vanillin when expressed in *P. putida* KT2440. Phenyl glyoxylate decarboxylases with potential activity on vanillyl glyoxylate have been identified due to the role of vanillyl glyoxylate and phenyl glyoxylate as intermediates in the degradation pathways of vanillyl mandelate and mandelate, respectively (Hasson et al., 1995; Turner et al., 1996; Saehuan et al., 2007). For rapid testing of the three homologs, we placed each homolog under control of the *tac* promoter with the RBS site JER01 (Elmore et al., 2017) and integrated each operon into the RV site of a serine-recombinase assisted genetic engineering (SAGE)-compatible strain, AG5577 (Huenemann et al., 2025). Strains KMM326, KMM328, and KMM330 were designed to express *mdlC* from *P. putida* (Hasson et al., 1995), *dpgB* from *Pseudomonas stutzeri* ST-201 (Saehuan et al., 2007), and *EC844_1132* from *Acinetobacter calcoaceticus* (Turner et al., 1996), respectively.

Following strain construction, each strain (including AG5577 as a control) was cultivated in a microtiter plate with 5 mM vanillyl glyoxylate alone or 5 mM vanillyl glyoxylate alongside 5 mM vanillate. While no strain exhibited growth on vanillyl glyoxylate alone, KMM326, KMM328, and KMM330 grew to higher optical densities than the control strain AG5577 on 5 mM vanillyl glyoxylate and 5 mM VA (**Fig. 4b**). Initial and final measurements of vanillyl glyoxylate concentration confirmed KMM326, KMM328, and KMM330 utilized 1.6 ± 0.1, 1.6 ± 0.3, and 1.1 ± 0.2 mM vanillyl glyoxylate, respectively, compared to 0.11 ± 0.02 mM utilized by AG5577 (**Fig. 4c**, p-values < 0.05).

**Figure 4.**
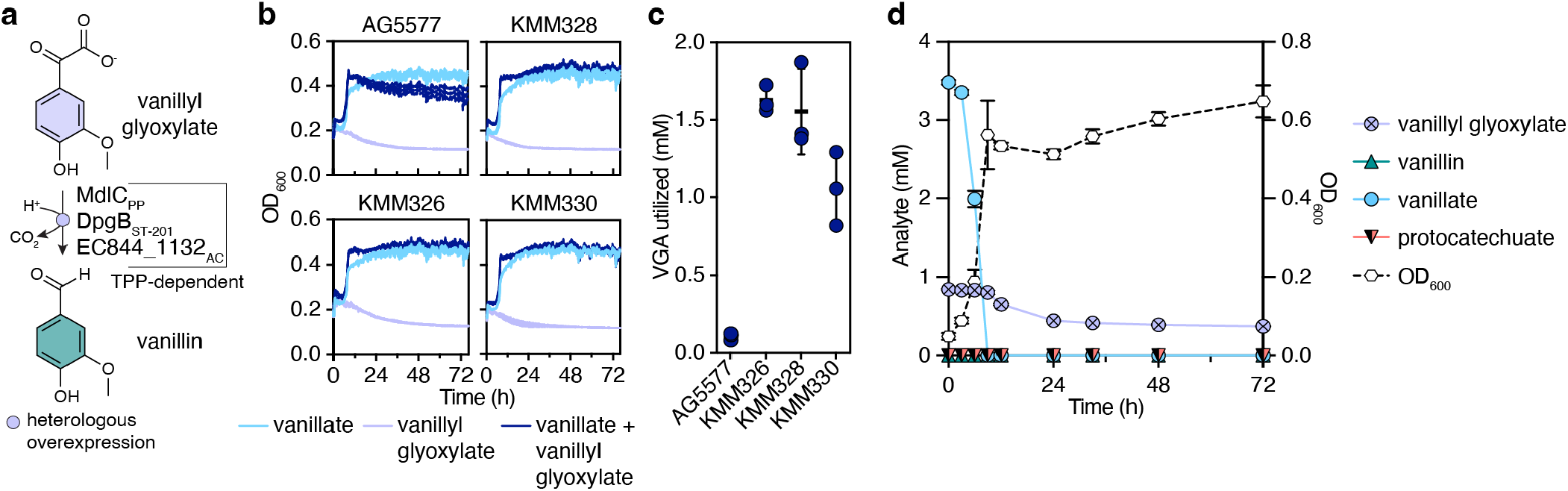
Phenyl glyoxylate decarboxylase homologs confer utilization of vanillyl glyoxylate in *P. putida* KT2440. **(a)** Three thiamine pyrophosphate (TPP)-dependent phenyl glyoxylate decarboxylase homologs, MdlC from *Pseudomonas putida* (previously *Arthrobacter siderocapsulatus*; PP), DpgB from *Pseudomonas stutzeri* ST-201 (ST-201), and EC844_1132 from *Acinetobacter calcoaceticus* (AC) were added to strain AG5577 to generate KMM326, KMM328, and KMM330. These enable the conversion of vanillyl glyoxylate to vanillin, which can natively be utilized by *P. putida* KT2440 for cellular growth. **(b)** Growth (OD_600_) curves for AG5577, KMM326, KMM328, and KMM330 microtiter plate cultivations in M9 media supplemented with 5 mM vanillate, 5 mM vanillyl glyoxylate, or 5 mM vanillate and 5 mM vanillyl glyoxylate. **(c)** Concentration of vanillyl glyoxylate consumed by strains cultivated in (b) with 5 mM vanillate and 5 mM vanillyl glyoxylate. **(d)** Shake flask cultivations of KMM326 in M9 media containing 3.5 mM vanillate and 0.5 mM vanillyl glyoxylate. For panels (b)-(d), the data points represent the average of three biological replicates, and the error bars represent the standard deviation. All data shown graphically in this figure are provided in **Data S1**.

We chose to further characterize vanillyl glyoxylate utilization by KMM326 in shake flasks with 3.5 mM VA and 1 mM vanillyl glyoxylate and measured analyte concentrations over time. KMM326 consumed all vanillate within 12 h but only utilized 56 ± 1 mol% of the vanillyl glyoxylate over the course of 72 h, with vanillyl glyoxylate utilization ceasing after 24 h (**Fig. 4d**). We posited that providing an additional carbon and energy source via a fed-batch approach would enable further utilization of vanillyl glyoxylate. Therefore, we evaluated vanillyl glyoxylate utilization in a muconate production strain with glucose supplementation, as described in the following section.

### 2.5. Expression of the SYK-6 acetovanillone pathway, LigW2, and MdlC confers muconate production from acetovanillone, 5-carboxyvanillate, 5-carboxyvanillin, 5-formylvanillin, and vanillyl glyoxylate

Having built strains for utilization of acetovanillone, 5-carboxyvanillate, 5-carboxyvanillin, and 5-formylvanillin through the common protocatechuate intermediate, we next combined the SYK-6 acetovanillone pathway and LigW2 for muconate production in a derivative of CJ781 (Kuatsjah et al., 2022) (**Table 1**, **Fig. 5a**). It has been observed previously that overexpression of catechol dioxygenases (*catA* or *catA2*) improve catechol conversion to muconate (Bleem et al., 2026; Kim et al., 2026; Wilkes et al., 2026; Kim et al., 1998), a strong RBS driving *aroY* translation in addition to helper proteins *EcdDB* improved protocatechuate conversion to catechol (Johnson et al., 2016; Sonoki et al., 2014; Wilkes et al., 2026), and a mutation in the *PP_3493* gene, which encodes a LysR family transcriptional regulator, increased growth rate of *P. putida* KT2440 on vanillate (Bleem et al., 2024). The resultant strain, MG080, contained each of these modifications in addition to overexpression of *acvABCDEF*_SYK-6_, *vceAB*_SYK-6_, and *ligW2*_SYK-6_ (see **Table 1** for genotype details).

**Figure 5.**
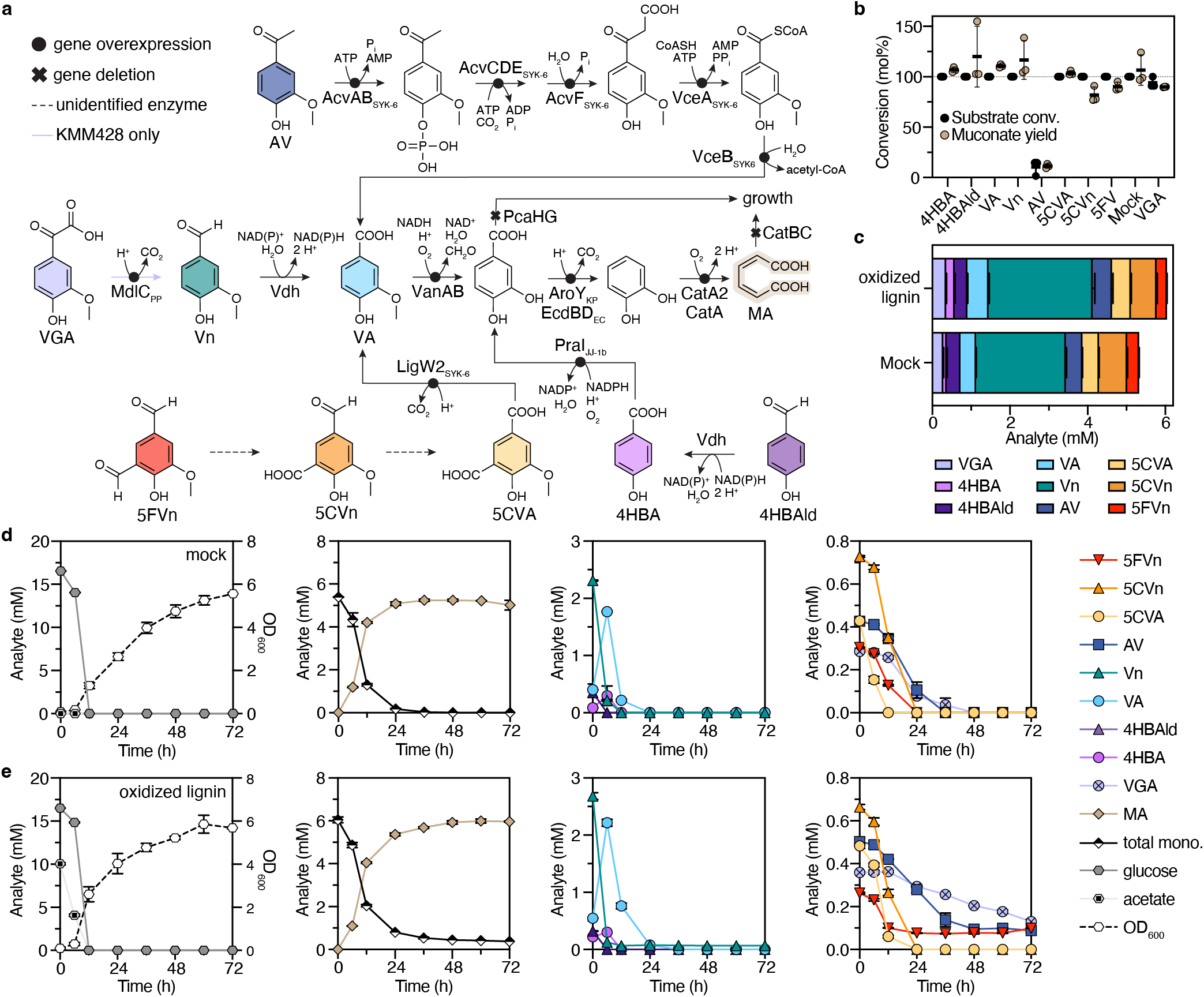
KMM428 produces muconate from pine-derived depolymerized kraft lignin. **(a)** Heterologous and native metabolic pathways for the conversion of acetovanillone (AV), vanillyl glyoxylate (VGA), vanillin (Vn), vanillate (VA), 5-formylvanillin (5FVn), 5-carboxyvanillin (5CVn), 5-carboxyvanillate (5CVA), 4-hydroxybenzaldehyde (4HBAld), and 4-hydroxybenzoate (4HB) to muconate in engineered *Pseudomonas putida* KT2440 strains MG080 and KMM428. Subscripts indicate a protein of heterologous origin, where SYK-6 indicates *Sphingobium lignivorans* SYK-6, PP represents *Pseudomonas putida*, JJ-1b signifies *Paenibacillus* sp. JJ-1b, EC indicates *Enterobacter cloacae*, and KP represents *Klebsiella pneumoniae*. Genotypes and strain construction details are provided in **Table 1 and Tables S2-S5**. Enzyme abbreviations: AcvAB, AVP/ASP synthetase; AcvF, AVP/ASP phosphatase; AcvCDE, biotin-dependent carboxylase; VceA, acetyl coenzyme A (acetyl-CoA)-dependent VAA/SAA-converting enzyme; VceB, vanilloyl-CoA/syringoyl-CoA thioesterase; LigW2, 5-carboxyvanillate decarboxylase; MdlC, phenyl glyoxylate decarboxylase; Vdh, vanillin dehydrogenase; VanAB, vanillate dioxygenase; PraI, para-hydroxybenzoate-3-hydroxylase; AroY, protocatechuate (PCA) decarboxylase; EcdBD, *Enterobacter cloacae* decarboxylase; CatA, catechol 1,2-dioxygenase; CatA2, catechol 1,2-dioxygenase; CatB, muconate cycloisomerase 1; CatC, muconolactone delta-isomerase; and PcaHG, PCA 3,4-dioxygenase. **(b)** Utilization of aromatic monomers (substrate conversion; conv.) and muconate yield by MG080 (all monomers except vanillyl glyoxylate) and KMM428 (vanillyl glyoxylate) when fed individual aromatic monomers or a mock mixture representative of oxidized lignin (excluding vanillyl glyoxylate) alongside glucose. **(c)** Molar composition of aromatic monomers present in the mock mixture and the oxidatively depolymerized and extracted kraft lignin sample (oxidized lignin) provided to KMM428 in cultivations displayed in d) and e). In addition to quantified aromatic analytes, the oxidized lignin contains unquantified unknowns. **(d)**, **(e)** KMM428 shake flask cultivations (30 °C and 225 rpm) with 15 mM glucose and 5 mM total aromatic monomers (total mono.) from either (d) a mock mixture or (e) oxidized lignin. For panels (b)-(e), the data points represent the average of three biological replicates, and the error bars represent the standard deviation. All data shown graphically in this figure are provided in **Data S1**.

Notably, MG080 converted eight of the monomers detected in the lignin oil – 4-hydroxybenzaldehyde, 4-hydroxybenzoate, vanillin, vanillate, acetovanillone, 5-carboxyvanillate, 5-carboxyvanillin, and 5-formylvanillin – to muconate in shake flasks with a co-feed of glucose for cellular growth (**Fig. 5b, Fig. S9**). While MG080 only consumed a small amount of acetovanillone, resulting in a muconate yield of 11 ± 2 mol%, MG080 fully utilized all other substrates provided in the media giving 82 ± 8 mol%, 90 ± 4 mol%, and quantitative yields for muconate from 5-carboxyvanillin, 5-formylvanillin, and all other substrates, respectively. Excitingly, when provided 4 mM of a mixture of aromatic monomers representative of pine-derived oxidatively depolymerized lignin streams (excluding vanillyl glyoxylate, which we detected and identified after this experiment was conducted), MG080 utilized 100 mol% of all monomers, including acetovanillone, achieving a quantitative muconate yield (**Fig. 5b, Fig. S9**). As MG080 successfully produced muconate at high yields from a relevant mixture of aromatic monomers, we used MG080 as our base strain for further engineering muconate production from vanillyl glyoxylate.

We next added the synthetic operon for *mdlC*_PP_ expression to MG080 at the *hsdRM* locus to generate KMM428 (**Table 1**) and tested the ability of KMM428 to convert 1 mM vanillyl glyoxylate to muconate alongside a glucose feed for cellular growth. KMM428 consumed 94.0 ± 5.2 mol% of the vanillyl glyoxylate fed, resulting in a muconate yield of 89.9 ± 0.4 mol% (**Fig. 5b**, **Fig. S10**). The sustained presence of an additional carbon and energy source (via a fed-batch glucose feed) led to higher utilization of vanillyl glyoxylate than observed with the batch approach described above (**Fig. 4d**). Overall, this engineering workflow resulted in a *P. putida* KT2440 strain capable of muconate production from all nine aromatic monomers quantified in the pine-derived oxidatively depolymerized lignin stream.

### 2.6. KMM428 simultaneously converts nine monomers from kraft lignin-derived aromatic monomers to muconate at near-quantitative yields

To determine the capacity of KMM428 to convert monomers from oxidatively depolymerized and extracted kraft lignin (oxidized lignin) compared to a mixture prepared from pure monomers (denoted as “mock”), we cultivated KMM428 with ∼5 mM total aromatic monomers (batch) in shake flasks alongside glucose for growth (fed-batch). The mock mixture was prepared to have similar concentrations of each monomer present in the oxidized lignin (**Fig. 5c**). The oxidized lignin only contains 17 wt% quantified monomers, meaning the remaining 83 wt% of the sample is unquantified and likely contains oligomeric lignin according to GPC (*i.e.*, components with molecular weights >200 Da; **Fig. S1**).

While KMM428 utilized all aromatic monomers from the mock mixture and converted them to muconate at a yield of 97 ± 1 mol% within 48 h, after 72 h, KMM428 utilized 93.6 ± 0.2 mol% of the quantified aromatic monomers from oxidized lignin (**Fig. 5d-e**). The 3 aromatic monomers that remained after the 72-h cultivation period were acetovanillone (18 ± 2 mol%; 0.09 ± 0.01 mM remaining), 5-formylvanillin (38 ± 1 mol%; 0.099 ± 0.001 mM remaining), and vanillyl glyoxylate (36 ± 4 mol%; 0.13 ± 0.01 mM remaining). Despite the small amount of quantified aromatic monomers remaining, muconate yields for the oxidized lignin case were 99 ± 3 mol%, suggesting conversion of unquantified compounds in the sample to muconate. Overall, these metabolic engineering efforts show that *P. putida* KT2440 can be engineered to convert all the quantified aromatic monomers present in depolymerized kraft lignin stream to muconate at 100% theoretical yield.

## 3. Discussion

In this work, we demonstrate that Cu-catalyzed oxidative depolymerization can generate bioavailable aromatic monomers from pine-derived kraft lignin and that these monomers were efficiently converted to muconate by an engineered *P. putida* KT2440 strain. The *P. putida* KT2440 strain developed in this work, KMM428, represents a single engineered strain capable of producing muconate from all nine identified aromatic monomers: 4-hydroxybenzoate, 4-hydroxybenzaldehyde, vanillin, vanillate, acetovanillone, 5-carboxyvanillate, 5-carboxyvanillin, 5-formylvanillin, and vanillyl glyoxylate. Overall, KMM428 expanded bioconversion for this process by 37.6 ± 0.1 mol% and achieved muconate yields of 99 ± 3 mol% from quantified monomers in an oxidized lignin stream. More broadly, this work expands the carbon bioconversion beyond previous efforts that targeted conversion of subsets of these compounds (Kamada et al., 2024; Omolabake et al., 2026; Lalande et al., 2026a).

In parallel work, *Rhodococcus aromaticivorans* RHA1 was engineered to convert vanillin, vanillate, and acetovanillone from a similar pine kraft lignin stream to muconate (Lalande et al., 2026a). Lalande *et al*. utilized a fed-batch approach to overcome initial muconate yield limitations caused by protocatechuate build-up and achieved a muconate yield of 99 mol% with resting cells. Protocatechuate build-up continued to limit this process, albeit to a lesser extent. In comparison, the *P. putida* KT2440 strain developed in this work, KMM428, was not limited by protocatechuate build-up in batch cultivations, but rather by slow conversion of acetovanillone, 5-formylvanillin, and vanillyl glyoxylate. Like RHA1, this limitation may be overcome with a fed-batch approach. Although engineered RHA1 and KMM428 each achieved near-theoretical yields in shake flasks, bioreactor studies are needed to evaluate scaled-up performance including with authentic lignin-derived substrates.

Improving utilization of acetovanillone, 5-formylvanillin, and vanillyl glyoxylate represents a clear target for future strain engineering. Potential approaches include bioprospecting for more active enzymes and transporters (Lämmle et al., 2007; Roy et al., 2025; Xiang et al., 2025), adaptive laboratory evolution (Mueller et al., 2022; Bleem et al., 2024; Feist and Woo, 2026), and dynamic regulation strategies (Dahl et al., 2013; Elmore et al., 2021; Xu et al., 2026). For acetovanillone utilization, optimizing expression of the *hpe* pathway from GD02 or different combinations of genes from the identified pathways from SYK-6, GD02, and *A. macra* could enable growth of *P. putida* KT2440 on acetovanillone as the sole carbon source and full conversion of acetovanillone to muconate, similar to observations in RHA1 (Lalande et al., 2026b, 2026a). In the case of 5-formylvanillin, aldehyde dehydrogenases from *P. putida* KT2440 active on 5-formylvanillin and 5-carboxyvanillin could be identified via random barcoded transposon insertion sequencing (Borchert et al., 2023; Thompson et al., 2020; Wetmore et al., 2015) or CRISPR interference (Bales et al., 2024; Batianis et al., 2020; Fenster et al., 2022) screening and subsequently overexpressed or engineered for improved activity. For vanillyl glyoxylate, additional phenyl glyoxylate decarboxylase homologs and the impact of thiamine pyrophosphate availability warrant further investigation. Intracellular thiamine pyrophosphate levels, which are highly regulated in bacterial cells by the thiamine pyrophosphate binding riboswitch (Jurgenson et al., 2009; Miranda-Ríos, 2007), could be limiting to TPP-dependent MdlC-mediated conversion of vanillyl glyoxylate. Addressing these remaining metabolic bottlenecks in engineered *P. putida* KT2440 will be critical for achieving efficient, complete conversion of oxidized lignin streams.

Beyond expanding the metabolic capabilities of *P. putida* KT2440, overcoming process-level limitations associated with deconstructed lignin streams will also be essential for efficient biological conversion. Low aromatic monomer yields from lignin and dilute concentrations of lignin-derived aromatic compounds in deconstructed lignin streams have historically limited bioconversion of lignin to products (Salvachúa et al., 2018; Sonoki et al., 2018; Becker et al., 2018a; Werner et al., 2023). While low aromatic purity does not always impact small-scale strain performance (Omolabake et al., 2026; Vilbert et al., 2023; Weiland et al., 2025), in which strains are tested in batch or with low feed rates, these effects become more apparent during the scaling process. Almqvist et al. showed explicitly that concentrating and purifying bio-available aromatics from non-bioavailable components via rotary evaporation and liquid-liquid extraction improves *P. putida* KT2440 growth rate and muconate titers from deconstructed pine kraft lignin (Almqvist et al., 2021)8/18/26 9:40:00 PM. Thus, integrating lignin-derived aromatic purification methods, such as centrifugal partition chromatography (Alherech et al., 2021; Lalande et al., 2026a) and membrane filtration (Aher et al., 2020; Li et al., 2019; Saboe et al., 2024), with bioconversion represents a promising opportunity to advance the field.

## 4. Materials and Methods

### 4.1. Materials and reagents

All commercial reagents were purchased and used as received; reagents and the corresponding vendors are listed in the supplementary information. Vanillyl glyoxylate was synthesized as described in the supplementary methods and displayed in **Fig. S11**.

### 4.2. Slug flow reactor design

A custom-built continuous slug-flow reactor was constructed following the block flow diagram shown in **Fig. S12**. The liquid feed solution was pumped using dual Teledyne ISCO (260D) syringe pumps operating in continuous mode. Oxygen (Airgas, industrial grade) was introduced using a Teledyne mass flow controller calibrated at 25 °C to 1,000 sccm O_2_. The liquid and gas phases were combined using a Swagelok T-joint before flowing through the reaction loop. The reaction loop consisted of a coiled 316 stainless steel tube (2 mm i.d., 60 mL reactor volume) submerged in an oil bath (Chemglass high-temperature silicone bath oil, CG-1100-31; operating temperature up to 315 °C) held at 210 °C. The reaction loop was connected to a heat exchanger constructed by swaging the reaction tubing inside a 3/8” o.d. 316 stainless steel tube equipped with water inlet/outlet connected to a Vivosun submersible pump in a 5-gallon bucket of water. The reactor pressure was maintained using a 316 stainless steel Equilibar back pressure regulator equipped with a PTFE Glass laminate diaphragm.

### 4.3. Cu-catalyzed oxidation of pine kraft lignin

A 30.72 kg lignin feed was prepared by adding 2.68 kg wet lignin (33% moisture content; 1.79 kg dry lignin) to 28.04 kg of 2 M NaOH solution, corresponding to approximately 6 wt% lignin on a dry-lignin basis. CuSO_4_·5H_2_O (8.04 g, 32.2 mmol) was dissolved in 10 mL of deionized water and added to the lignin solution under continuous agitation with an overhead mixer to ensure homogeneous mixing. The solution and oxygen were then co-fed through the slug flow reactor at a liquid flow rate of 5 mL/min with an oxygen flowrate of 360 mL/min under a back pressure of 311 psi. The reactor operated under a slug-flow regime with a residence time of 20.3 s. The O_2_/lignin molar ratio was estimated to be 9.0 by calculating relative molar flow rates using a lignin repeat unit molecular weight of 196.2 g/mol for an idealized poly-coniferyl alcohol. After thermal equilibration of the reactor using deionized water, the feed stream was switched to the lignin stock solution and continuously processed for 89.5 h at 210 °C. The bulk solution was acidified to pH 2 under constant mechanical stirring using concentrated sulfuric acid added in small portions to allow effervescence to subside between additions. The resulting dark precipitate was then removed using a coarse sieve. The resulting solution was then stored for analysis and further extraction.

### 4.4. Analytics of deconstructed lignin

Sample composition of the acidified deconstructed lignin was determined using a Waters Acquity Class H QSM Plus UPLC system equipped with a BEH C18 column (1.7 μm, 2.1 x 50 mm) heated to 40 °C. Data acquisition was performed using Empower software, with calibration curves and sample traces obtained via a photodiode array monitoring elution at 280 nm. 10 mM 1,4-dimethoxybenzene was used as an internal standard. Calibration curves for all compounds were obtained by injection of solutions after serial dilution of a stock analyte solution (**Fig. S13**, **Table S6**). Elution was performed using a UPLC gradient profile (**Table S7**) using 0.1% formic acid in water (solvent A) and 0.1% formic acid in acetonitrile (solvent B).

### 4.5. Counter-current extractor design

A counter-current extractor was constructed as depicted in **Fig. S14**. Glass pieces were blown at the University of Wisconsin-Madison Chemistry Glass Laboratory. Aqueous and organic liquids were pumped using dual Iwaki metering pumps (model EZBD1). The pumps were connected to the glass mixing/settling chambers using PharMed^®^ BPT tubing. The column was stirred using a Caframo mechanical stirrer connected to a 32-inch stainless steel rod fitted with 5 impellers and 6 disks positioned alternatingly down the rotating shaft.

### 4.6. Isolation and gel permeation chromatography of deconstructed lignin

The acidified deconstructed lignin solution described above was extracted using the counter-current extractor using flow rates of 15 mL/min for both the ethyl acetate and the aqueous feed and a stirring rate of 800 rpm. The ethyl acetate extract was collected and concentrated using an Across International 5 L rotary evaporator. We obtained 8.11 wt% of the initial lignin substrate as isolated oxidized lignin. Aromatic monomers present in the resultant deconstructed lignin were quantified prior to biological conversion as described below. Analytical GPC was performed on a Shimadzu Prominence LC20 with a photodiode array detector (SPD-M20A). Separation was performed using a PSS PolarSil Linear S column (7.8 mm ID × 30 cm L × 30 μm particle size) in a 50 °C oven, with 0.1 M lithium bromide (LiBr) in *N*,*N*-dimethylformamide (DMF) flowing at 0.5 mL/min for 40 min. The samples were prepared as 0.33 mg/mL solution of lignin in 0.1 M LiBr in DMF. The molecular weight distribution was calibrated at λ = 270 nm using PSS Polystyrene ReadyCal Standard Set *M*(p) 474–2,520,000 Da (P/N PSS-pskitr4; PSS-Polymer Standards Service, Amherst, MA, USA) and acetovanillone (166 Da).

### 4.7. Preparation of aromatics for bacterial cultivations

Aromatics were prepared for bacterial cultivations by dissolving in ethanol, DMSO, or water, depending on the study. When ethanol was present as a co-substrate, acetovanillone, vanillate, and acetosyringone were solubilized in ethanol at 0.5 M, 0.5 M, and 0.25 M, respectively (data represented in **Fig. 2c, d**). For studies in which aromatics were fed individually as the sole carbon source (data in **Figs. 2b, 3-4**) or with glucose (data represented in **Figs. 5b, S9, S10**), aromatics were dissolved in DMSO between 0.25 and 2 M. DMSO concentrations in the final media never exceeded 1% v/v. We note acetosyringone showed limited solubility in ethanol, while 5-carboxyvanillate and 5-carboxyvanillin showed limited solubility in DMSO, and were therefore fed at lower concentrations than the other aromatics. For the final study in which KMM428 converted aromatics from a mock mixture or oxidized lignin to muconate (data in **Fig. 5c-e**), the oxidized lignin was dissolved in water at 50 mg/mL by slowly pH adjusting the solution to neutral pH with 4 M NaOH. The composition of the resulting solution was determined via ultra-high-performance liquid chromatography with diode array detection (UHPLC-DAD) as described in section 4.10 (**Fig. 1b**). Similarly, the aromatics in the mock mixture were dissolved in water as a mixture at the same concentration as in the aqueous oxidized lignin solution by slowly pH adjusting to neutral pH with 4 M NaOH.

### 4.8. Plasmid and strain construction

All plasmids, oligonucleotides, and strains in this study are listed in **Table 1**, **Tables S2-S4**. Sequences of synthesized DNA used in this study are listed in **Table S5**. Coding sequences from heterologous sources were codon-optimized for *P. putida* using the Integrated DNA Technologies (USA) codon optimization tool. Synthetic ribosome binding sites were generated using the Salis RBS calculator (Cetnar and Salis, 2021; Reis and Salis, 2020). Plasmids were either ordered from Twist Bioscience (USA) or cloned using standard protocols. In brief, polymerase chain reaction (PCR) was performed using Q5 (NEB) and plasmid construction was performed using NEBuilder HiFi DNA Assembly (NEB). Plasmid sequences were confirmed upon construction via Plasmidsaurus. Engineered strains of *P. putida* KT2440 were generated either via homologous recombination (Johnson and Beckham, 2015; Schäfer et al., 1994) or serine-recombinase assisted genome engineering (SAGE) (Elmore et al., 2017; Huenemann et al., 2025) as previously described. Genetic deletions and insertions were confirmed via colony polymerase chain reaction with MyTaq™ HS Red Mix (Bioline) and sequencing via Plasmidsaurus.

### 4.9. Cultivation of Pseudomonas putida KT2440 strains

For microtiter plate and shake-flask cultivations, *P. putida* KT2440 strains were first grown overnight in LB broth (Miller) inoculated from glycerol stocks at 30 °C and 225 rpm. Cells were subsequently washed with 1x M9 salts and transferred to M9 minimal medium (6.78 g/L Na_2_HPO_4_, 3 g/L KH_2_PO_4_, 0.5 g/L NaCl, 1 g/L NH_4_Cl, 2 mM MgSO_4_, 100 μM CaCl_2_, and 18 μM FeSO_4_; pH 7.0) supplemented with the indicated aromatic substrate(s), ethanol, or glucose. Cultures were inoculated in triplicate to an initial optical density at 600 nm (OD₆₀₀) of 0.1. For microtiter plate cultivations, cells were cultivated in 200 μL volumes in Honeycomb 100-well plates at 30 °C with maximum orbital shaking. Cell growth was monitored using a Bioscreen C® (Growth Curves) by measuring absorbance at 600 nm every 15 min. For shake flask cultivations, cells were cultivated in 125 mL baffled, glass Erlenmeyer flasks (metal caps) with a 25-30 mL working volume at 30°C and 225 rpm in a benchtop incubator (0.75” orbital) for 72 h. In the case of muconate production studies, glucose was added to the initial media at 15 mM and fed to 10 mM every 12 h of cultivation. 800 μL samples were taken at specified times to determine cell growth via OD_600_ and metabolite concentration via HPLC diode array. For metabolite analysis, samples were centrifuged for 2 min at >18,000*g*, and filtered (0.2 μm syringe filter) into amber glass vials. Samples were diluted 10x in water for aromatic and muconate analysis; they were stored at −20 °C prior to analysis.

### 4.10. Quantification of metabolic analytes

Vanillin, vanillate, 4-hydroxybenzaldehyde, 4-hydroxybenzoate, protocatechuate, catechol, and muconate were quantified by UHPLC-DAD as described on protocols.io (Woodworth et al., 2024). This method was expanded to quantify 5-carboxyvanillate, 5-carboxyvanillin, 5-formylvanillin, and vanillyl glyoxylate by generating standard curves for each new compound under the same conditions (**Fig. S15, S16**). Muconate yields were calculated as *Y_muconate_ = mol_muconate produced_ / ∑mol_bioavailable aromatics fed_*, where the sum of bioavailable aromatics fed is the sum of quantified 4-hydroxybenzoate, 4-hydroxybenzaldehyde, vanillate, vanillin, acetovanillone, 5-carboxyvanillate, 5-carboxyvanillin, 5-formylvanillin, and vanillyl glyoxylate added to the cultivation media and muconate is the measured sum of both *cis, cis*-muconate and *cis, trans*-muconate. Ethanol, acetate, and glucose were quantified using high-performance liquid chromatography with refractive index detection as described on protocols.io (Alt et al., 2024).

## Data availability

All data from this study are included in the main text, supplementary information, and Data S1.

## Acknowledgements

This work was authored in part by the National Laboratory of the Rockies for the U.S. Department of Energy (DOE) under Contract No. DE-AC36-08GO28308. Funding to KMM, DTH, PD, AS, TWR, SSS, GTB, and AZW, was provided by the U.S. DOE Office of Critical Minerals and Energy Innovation Alternative Fuels and Feedstocks Office under Award DE-EE0011114. For KMM, GTB, and AZW this material is based upon work at the Center for Bioenergy Innovation supported by the U.S. Department of Energy, Office of Science, Biological and Environmental Research under Contract Number ERKP886. Funding to PSZ and use of NMR spectrometers and gel permeation chromatography were supported by Great Lakes Bioenergy Research Center, U.S. Department of Energy, Office of Science, Biological and Environmental Research Program under Award Number DE-SC0018409. The authors thank Lindsay D. Eltis and Anne T. Lalande for thoughtful scientific discussion. The authors thank Adam M. Guss for providing strain AG5577.

The views expressed in the article do not necessarily represent the views of the DOE or the U.S. Government. The U.S. Government retains and the publisher, by accepting the article for publication, acknowledges that the U.S. Government retains a nonexclusive, paid-up, irrevocable, worldwide license to publish or reproduce the published form of this work, or allow others to do so, for U.S. Government purposes.

## Author contributions

Conceptualization: S.S.S., A.Z.W., and G.T.B. Methodology: K.M.M., D.T.H, M.A.G., P.D., P.S.Z., A.S., M.A.I., A.F.B., K.J.R, T.W.R., S.S.S., A.Z.W., G.T.B. Investigation: K.M.M., D.T.H, M.A.G., P.D., P.S.Z., A.S., M.A.I., A.F.B., K.J.R. Visualization: K.M.M. and D.T.H. Funding acquisition: T.W.R., S.S.S., A.Z.W., and G.T.B. Supervision: T.W.R., S.S.S., A.Z.W., and G.T.B. Writing—original draft: K.M.M., D.T.H., K.J.R., S.S.S, A.Z.W., and G.T.B. Writing—review and editing: all authors reviewed and approved the manuscript.

