## Supplemental Information for "Expanding the catabolic capacity of *Pseudomonas putida* to acetovanillone, 5-carboxyvanillate, and vanillyl glyoxylate for muconate production from kraft lignin-derived aromatics"

**The PDF file includes:**

Materials
Supplementary methods
Figs. S1 to S22
Tables S1 to S7
References

**Other Supplementary Materials for this manuscript include the following:**

Data S1

### Materials

Unless otherwise stated, all reagents were purchased from commercial sources and used without further purification. The following chemicals were used:

• 4-hydroxybenzaldehyde (Sigma-Aldrich, Cat. #144088-250g)
• 4-hydroxybenzoic acid (Sigma-Aldrich, Cat. #240141-50g)
• vanillin (Sigma-Aldrich, Cat. #V1104-500g)
• vanillic acid (Thermo-Scientific, Cat. #A12074.22)
• 5-carboxyvanillic acid (Aurum Pharmatech LLC, Cat. #MR16997)
• 5-carboxyvanillin (Biosynth, Cat. #FC70545)
• 5-formylvanillin (Ambeed, Cat. #A1323729)
• acetovanillone (Sigma-Aldrich, Cat. #W508454)
• vanillyl glyoxylic acid (prepared as described in supplementary methods below)
• acetosyringone (Thermo-Scientific, Cat. #115540050)
• syringic acid (AK-Scientific, Cat. #C591-500g)
• *cis, cis*-muconic acid (Fluka Chemical Corporation, Cat. #15992-5g)
• ethanol (KOPTEC, Cat. #V1016G)
• D-glucose (Fisher Chemical, Cat. #D16-10)
• 1,4-dimethoxybenzene (Fisher Scientific)
• sodium hydroxide (Fisher Scientific)
• copper (II) sulfate (Fisher Scientific)
• HPLC grade methanol (Fisher Scientific)
• HPLC grade ethyl acetate (Fisher Scientific)

**Supplementary methods**

Synthesis of vanillyl glyoxylic acid (**VGA**) over three steps (**Fig. S11, S17-S22**).

Benzoyl acetovanillone (**1**): Acetovanillone (20.81 g, 125.2 mmol, 1.00 equiv) and chloroform (250 mL) were added to a 500 mL round bottom flask, followed by the slow addition of a mixture of pyridine (11.58 mL, 143.8 mmol, 1.15 equiv) and benzoyl chloride (16.70 mL, 143.8 mmol, 1 equiv). The mixture was stirred at room temperature overnight, then concentrated *in vacuo*. 300 mL of water was added to the residue and the mixture was extracted with ethyl acetate. The combined extracts were dried over magnesium sulfate and evaporated to dryness *in vacuo*. The crude product was recrystallized using an ethyl acetate-pentane mixture to afford the product as fine white needles (22.11 g, 65%). $^{1}$H NMR (500 MHz, $CDCl_3$) δ 8.25 – 8.19 (dd, *J* = 7 Hz, 1Hz, 2H), 7.69 – 7.63 (m, 2H), 7.61 (dd, *J* = 8.1, 1.9 Hz, 1H), 7.53 (t, *J* = 8 Hz, 2H), 7.26 (d, *J* = 8 Hz, 1 H), 3.88 (s, 3H), 2.63 (s, 3H). $^{13}$C NMR (126 MHz, $CDCl_3$) δ 197.02, 164.29, 151.60, 144.07, 135.95, 133.78, 130.37, 128.90, 128.61, 122.94, 122.02, 111.44, 56.07, 26.62.

Benzoyl vanillyl glyoxylic acid (**2**): Using a previously described method with a few modifications (Scheffer and Wang, 2001). Benzoyl acetovanillone (**1**, 6.76 g, 25.0 mmol, 1.00 equiv), $SeO_2$ (4.44 g, 40.0 mmol, 1.60 equiv), and pyridine (100 mL) were added to a 250 mL round bottom flask and charged with $N_2$. The mixture was heated to 100 °C under $N_2$ for 3.5 h, then cooled to room temperature and filtered, the filtrate was dried *in vacuo*. The residue was dissolved in 100 mL 2 M HCl and extracted with ethyl acetate. The combined organic extracts were washed with water, then extracted with saturated $NaHCO_3$ solution. The combined aqueous extracts were acidified with concentrated HCl and again extracted with ethyl acetate. The combined organic phases were washed with brine, dried over magnesium sulfate, and evaporated to dryness *in vacuo*. The crude product was recrystallized using a dichloromethane-heptane mixture to afford the product as a light yellow powder (5.07 g, 68%). $^{1}$H NMR (500 MHz, $CDCl_3$) δ 8.21 (td, *J* = 8.0, 1.6 Hz, 3H), 8.02 (d, *J* = 2.0 Hz, 1H), 7.71 – 7.64 (m, 1H), 7.54 (t, *J* = 7.3 Hz 1H), 7.34 (d, *J* = 8.3 Hz, 1H), 3.91 (s, 3H). $^{13}$C NMR (126 MHz, $CDCl_3$) δ 183.07, 164.14, 161.50, 151.85, 146.31, 134.02, 130.45, 130.36, 128.69, 128.53, 125.69, 123.58, 113.89, 77.28, 77.02, 76.77, 56.19.

Vanillyl glyoxylic acid (**VGA**): Benzoyl vanillyl glyoxylic acid (**2**, 93.2 mg, 0.310 mmol, 1.00 equiv), $K_2CO_3$ (42.8 mg, 0.309 mmol, 1.00 equiv), and 3 mL methanol were added to an 8 mL glass vial. The mixture was heated to reflux for 4 h, cooled to room temperature and quenched with 2 M HCl until pH ~1, then dried *in vacuo*. The residue was purified by flash chromatography over silica gel using a methanol-dichloromethane gradient elution to yield the product as a yellow solid (40 mg, 48%). $^{1}$H NMR (500 MHz, $D_2O$) δ 7.42 – 7.27 (m, 2H), 6.82 (d, *J* = 8.8 Hz, 1H), 3.74 (s, 3H). $^{13}$C NMR (126 MHz, $D_2O$) δ 215.24, 193.54, 171.84, 152.06, 147.59, 126.30, 124.43, 115.17, 111.77, 55.62.

### Supplementary Figures

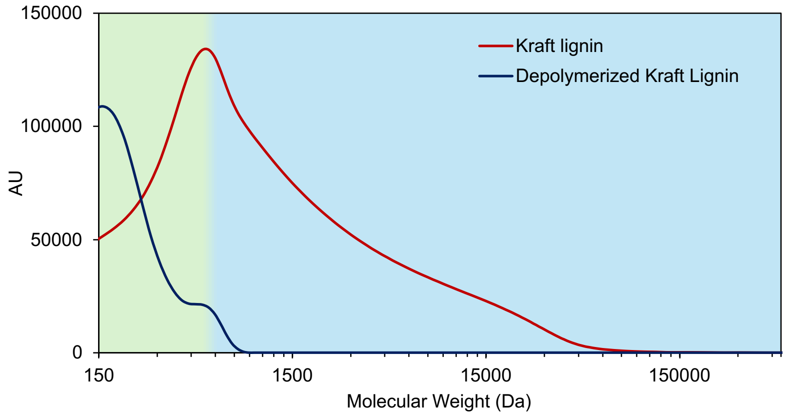

**Figure S1. GPC of Kraft lignin before and after oxidative depolymerization.** The green region represents low molecular weight components outside of the molecular weight calibration. Monomers typically have molecular weights <200 Da, while dimers, trimers, and oligomers are >200 Da.

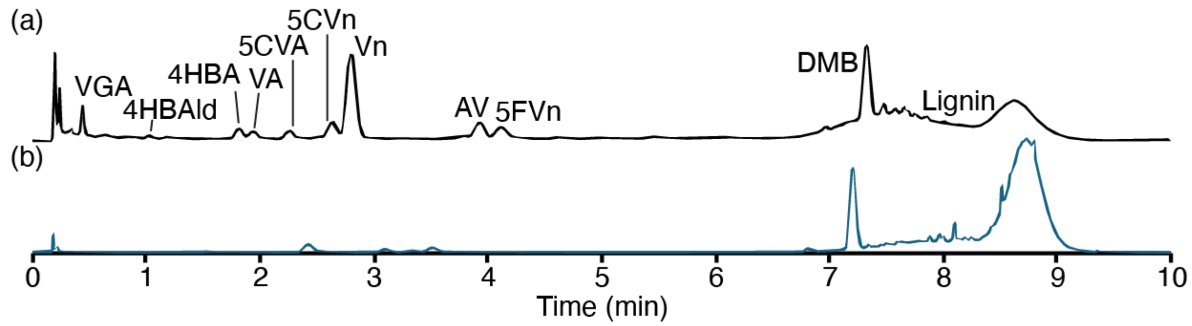

**Figure S2. UPLC trace of (a) reactor feed after oxidation acidified to pH < 2 and (b) the reactor feed containing 60 g/L kraft lignin.**

Abbreviations: VGA, vanillyl glyoxylate; 4HBAld, 4-hydroxybenzaldehyde; 4HB, 4-hydroxybenzoate; VA, vanillate; 5CVA, 5-carboxyvanillate; 5CVn, 5-carboxyvanillin; Vn, vanillin; AV, acetovanillone; 5FVn, 5-formylvanillin; DMB, 1,4-dimethoxybenzene.

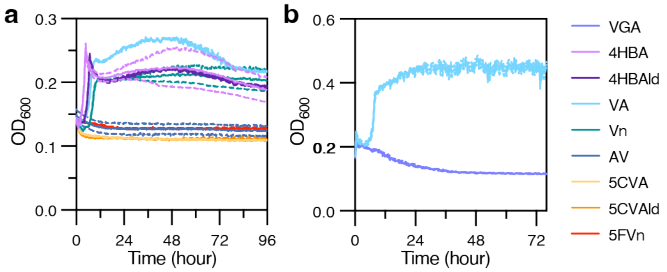

**Fig. S3. Wildtype *Pseudomonas putida* KT2440 grows on 4-hydroxybenzoate, 4-hydroxybenzaldehye, vanillin, and vanillate but not acetovanillone, 5-carboxyvanillate, 5-carboxyvanillin, 5-formylvanillin, or vanillyl glyoxylate.**

**(a)** Wildtype *Pseudomonas putida* (*P. putida*) KT2440 was cultivated in a microtiter plate (Bioscreen C format) in M9 minimal media supplemented with 5 mM 4-hydroxybenzoate (4HBA), 5 mM 4-hydroxybenzaldehyde (4HBAld), 5 mM vanillate (VA), 5 mM vanillin (Vn), 5 mM acetovanillone (AV), 2.5 mM 5-carboxyvanillate (5CVA), 2.5 mM 5-carboxyvanillin (5CVn), or 5 mM 5-formylvanillin (5FVn). $OD_{600}$ readings were taken every 15 minutes to evaluate bacterial cell growth. Solid and dashed lines represent the average and standard deviation, respectively, of three biological replicates. **(b)** Serine-recombinase assisted genetic engineering (SAGE)-compatible *P. putida* KT2440 strain, AG5577, was cultivated in a microtiter plate (Bioscreen C format) in M9 minimal media supplemented with 5 mM vanillate (VA) or 5 mM vanillyl glyoxylate (VGA). We note, strain AG5577 does not express any heterologous genes. $OD_{600}$ readings were taken every 15 minutes to evaluate bacterial cell growth. Solid and dashed lines represent the average and standard deviation, respectively, of three biological replicates.

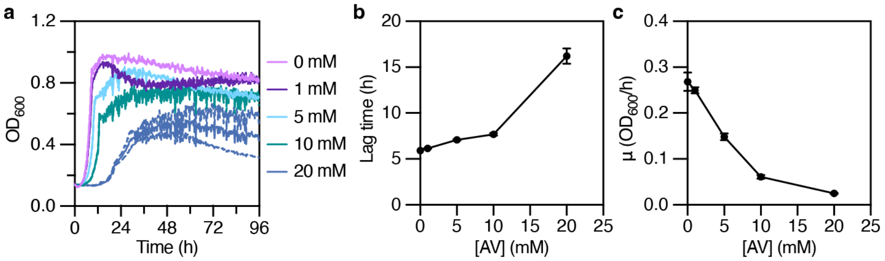

**Fig. S4. Growth of wildtype *Pseudomonas putida* KT2440 on glucose is negatively impacted by increasing acetovanillone levels.**

**(a)** Wildtype *Pseudomonas putida* (*P. putida*) KT2440 was cultivated in a microtiter plate (Bioscreen C format) in M9 minimal media supplemented with 20 mM glucose and increasing concentrations of acetovanillone. $OD_{600}$ readings were taken every 15 minutes to evaluate bacterial cell growth. Solid and dashed lines represent the average and standard deviation, respectively, of three biological replicates. **(b)** Growth rates and **(c)** lag times were determined by fitting growth curves to a Single-Gompertz model and extracting the parameters µ (absolute growth rate; $OD_{600}$/h) and lag time (h). In b and c, plots depict the mean and standard deviation for three biological replicates. Abbreviations: $OD_{600}$: optical density, measured as absorbance at 600 nm; h: hours.

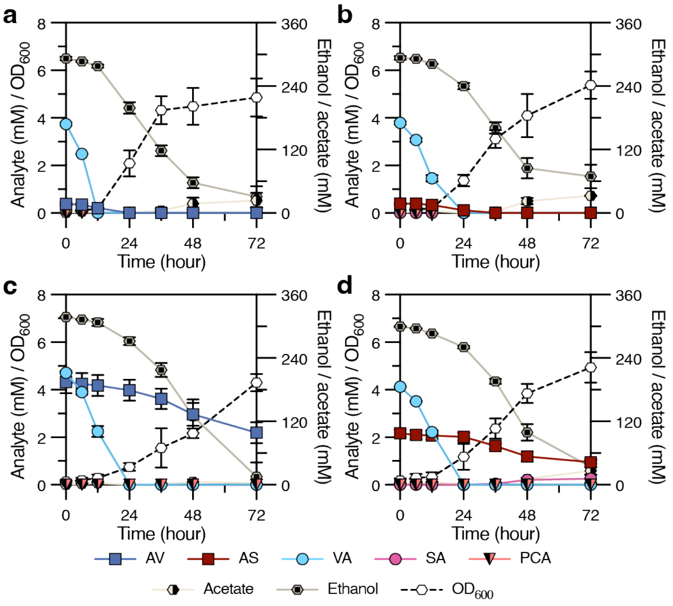

**Fig. S5. Utilization of acetovanillone and acetosyringone by KMM022.**
KMM022 cultivations in shake flasks with M9 minimal media supplemented with **(a)** 4.5 mM vanillate (VA) and 0.5 mM acetovanillone (AV), **(b)** 4.5 mM vanillate and 0.5 mM acetosyringone (AS), **(c)** 5 mM vanillate and 5 mM acetovanillone, or **(d)** 5 mM vanillate and 5 mM acetosyringone. Aromatics were dissolved in ethanol resulting in a final ethanol concentration of 2% volume ethanol/volume media. Cultivations were sampled at the time points indicated to evaluate growth by $OD_{600}$ (using a cell-free blank) and metabolite concentration. Symbols represent the average and error bars represent the standard deviation across three biological replicates. Abbreviations: AV, acetovanillone; AS, acetosyringone; VA, vanillate; SA, syringate; PCA, protocatechuate; $OD_{600}$, optical density, measured as absorbance at 600 nm.

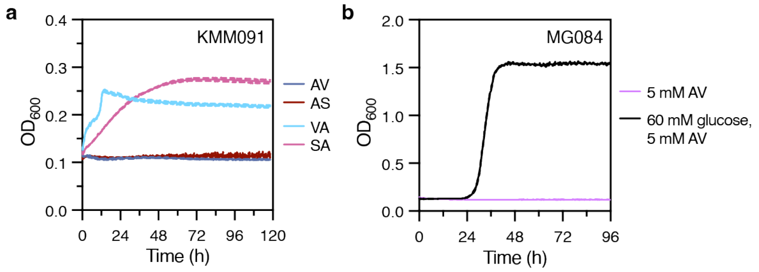

**Fig. S6. Overexpression of VanAB or MarK E16K does not enable growth on acetovanillone or acetosyringone.**

Engineered strains **(a)** KMM091 (*P. putida* KT2440 *fpva*::$P_{tac}$:*VceAB*$_{\text{SYK-6}}$:*AcvF*$_{\text{SYK-6}}$ *PP_5042*::$P_{tac}$:*AcvAB*$_{\text{SYK-6}}$ *PP_5322*:$P_{tac}$:*AcvCDE*$_{\text{SYK-6}}$ Δ*crc::*$P_{tac}$*:vanAB*) and **(b)** MG084 (*P. putida* KT2440 *fpva*::$P_{tac}$:*VceAB*$_{\text{SYK-6}}$:*AcvF*$_{\text{SYK-6}}$ *PP_5042*::$P_{tac}$:*MarK*$_{\text{NA}}$ *PP_5322*:$P_{tac}$:*AcvCDE*$_{\text{SYK-6}}$ *Δcrc*::$P_{tac}$:*LigW2*$_{\text{SYK-6}}$ were cultivated in a microtiter plate (Bioscreen C format). **(a)** KMM091 was cultivated in M9 minimal media supplemented with 5 mM acetovanillone (AV), 5 mM acetosyringone (AS), 5 mM vanillate (VA), or 5 mM syringate (SA). **(b)** MG084 was cultivated in M9 minimal media supplemented with 5 mM acetovanillone (AV) or 60 mM glucose and 5 mM acetovanillone. Acetovanillone was dissolved in dimethyl sulfoxide (DMSO); media contained final DMSO concentration of 1% volume/volume. $OD_{600}$ readings were taken every 15 minutes to evaluate bacterial cell growth. Solid and dashed lines represent the average and standard deviation, respectively, of three biological replicates. SYK-6 signifies *Sphingobium lignivorans* SYK-6 and NA represents *Novosphingobium aromaticivorans* DSM12444.

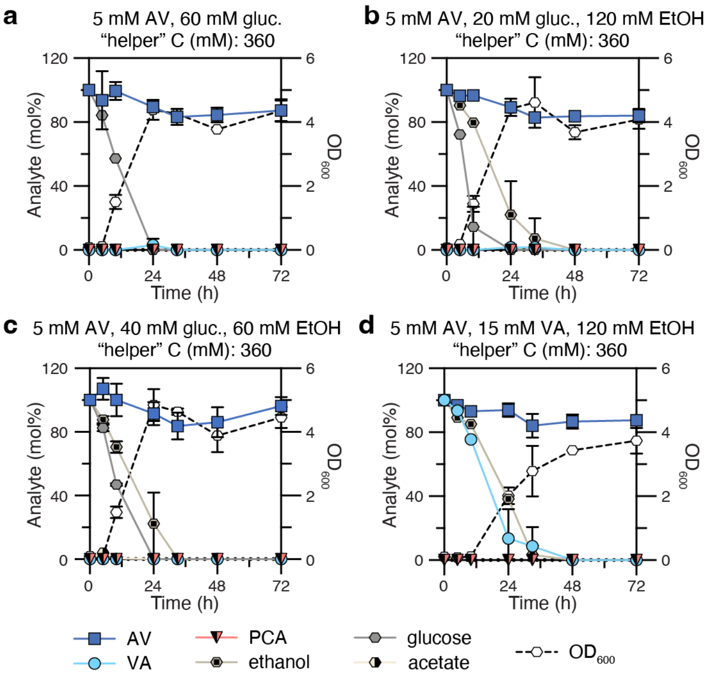

**Fig. S7. Identity of co-carbon does not impact acetovanillone utilization in KMM022.**
KMM022 cultivations in shake flasks with M9 minimal media supplemented with 5 mM acetovanillone (AV) and **(a)** 60 mM glucose (gluc.), **(b)** 20 mM glucose and 120 mM ethanol (EtOH) **(c)** 40 mM glucose and 60 mM ethanol, or **(d)** 15 mM vanillate (VA) and 120 mM ethanol. Aromatics were dissolved in water by pH adjusting to neutral pH slowly with 1 M sodium hydroxide. Cultivations were sampled at the time points indicated to evaluate growth by $OD_{600}$ (using a cell-free blank) and metabolite concentration. Symbols represent the average and error bars represent the standard deviation across three biological replicates. Abbreviations: AV, acetovanillone; VA, vanillate; PCA, protocatechuate; $OD_{600}$, optical density, measured as absorbance at 600 nm; h, hours; C, carbon; EtOH, ethanol; gluc., glucose.

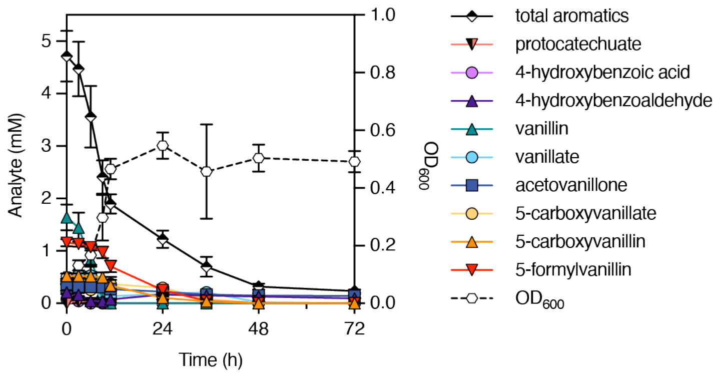

**Fig. S8. MG079 fully utilizes 1 mM 5-formylvanillin from aromatic mixture.**
MG079 was cultivated in shake flasks with M9 minimal medium supplemented with 5 mM of a mock mixture containing 1 mM 5-formylvanillin and other aromatic monomer components present in oxidized lignin. The mock mixture was prepared to have a similar composition of aromatics as the depolymerized pine-derived kraft lignin, except for higher 5-formylvanillin levels and the absence of vanillyl glyoxylate as MG079 cannot utilize this compound. The symbol 'total aromatics' represents the sum of 4-hydroxybenzaldehyde, 4-hydroxybenzoate, vanillin, vanillate, acetovanillone, 5-carboxyvanillate, 5-carboxyvanillin, and 5-formylvanillin. Symbols represent the average and error bars represent the standard deviation across three biological replicates. Abbreviations: $OD_{600}$: optical density, measured as absorbance at 600 nm; h, hours.

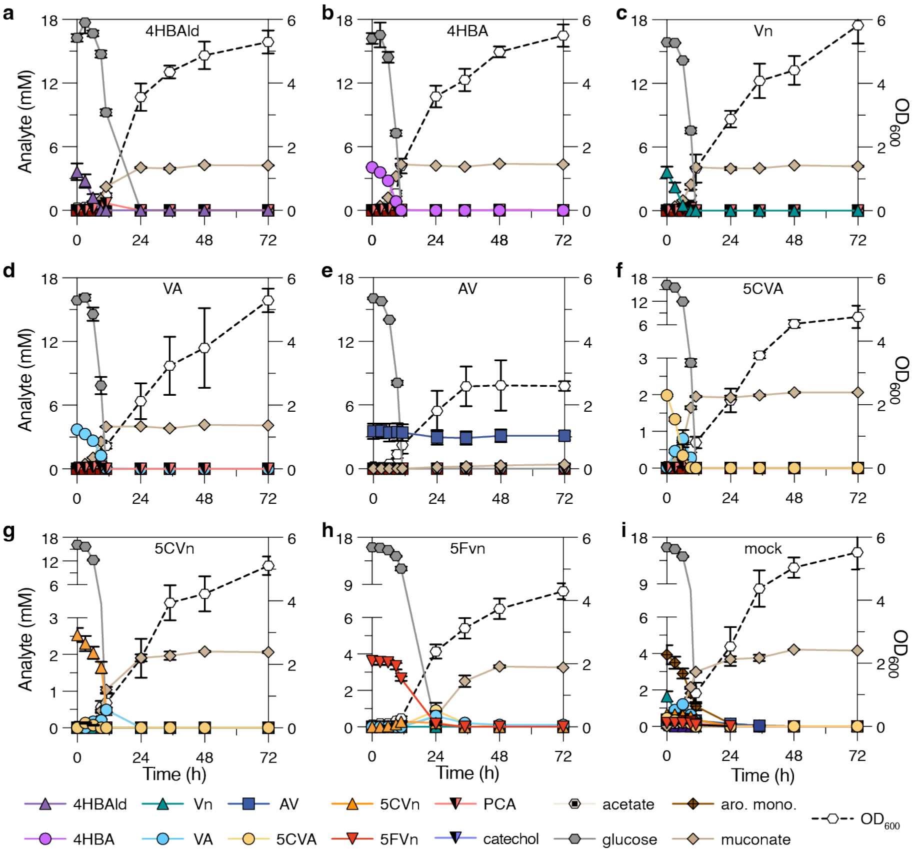

**Fig. S9. MG080 converts 4-hydroxybenzaldehyde, 4-hydroxybenzoate, vanillin, vanillate, acetovanillone, 5-carboxyvanillate, 5-carboxyvanillin, and 5-formylvanillin to muconate.**
MG080 was cultivated in shake flasks with M9 minimal medium supplemented with 15 mM glucose plus **(a)** 5 mM 4-hydroxybenzaldehyde, **(b)** 5 mM 4-hydroxybenzoate, **(c)** 5 mM vanillin, **(d)** 5 mM vanillate, **(e)** 5 mM acetovanillone, **(f)** 2.5 mM 5-carboxyvanillate, **(g)** 2.5 mM 5-carboxyvanillin, **(h)** 5 mM 5-formylvanillin, or **(i)** 5 mM of a mock mixture. The mock mixture was prepared to have the same composition of aromatics as the depolymerized pine-derived kraft lignin. Glucose was fed to 10 mM post sampling every 12 hours. The symbol 'aro. mono' (aromatic monomers) represents the sum of 4-hydroxybenzaldehyde, 4-hydroxybenzoate, vanillin, vanillate, acetovanillone, 5-carboxyvanillate, 5-carboxyvanillin, and 5-formylvanillin. Symbols represent the average and error bars represent the standard deviation across three biological replicates. Abbreviations: 4HBAld, 4-hydroxybenzaldehyde; 4HBA, 4-hydroxybenzoate; Vn, vanillin; VA, vanillate; AV, acetovanillone; 5CVA, 5-carboxyvanillate; 5CVn, 5-carboxyvanillin; 5FVn, 5-formylvanillin; PCA, protocatechuate; aro. mono., aromatic monomers; $OD_{600}$: optical density, measured as absorbance at 600 nm; h, hours.

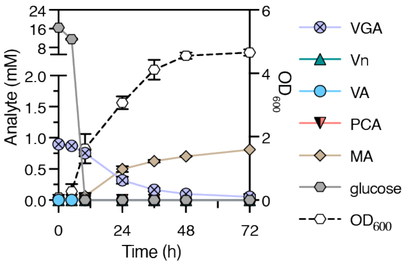

**Fig. S10. KMM428 produces muconate from vanillyl glyoxylate when cofed glucose.**
KMM428 was cultivated in shake flasks with M9 minimal medium supplemented with 15 mM glucose plus 1 mM vanillyl glyoxylate (VGA)**.** Glucose was fed to 10 mM post sampling every 12 hours. Symbols represent the average and error bars represent the standard deviation across three biological replicates. Abbreviations: VGA, vanillyl glyoxylate; Vn, vanillin; VA, vanillate; PCA, protocatechuate; MA, muconate; $OD_{600}$: optical density, measured as absorbance at 600 nm; h, hours.

**a**

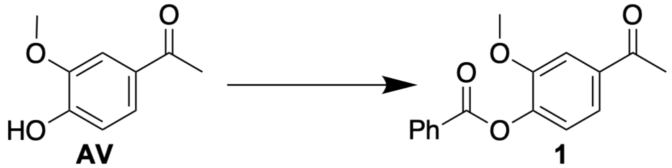

**AV** → **1**

[1]H NMR (500 MHz, $CDCl_3$) δ 8.25 – 8.19 (dd, *J* = 7 Hz, 1Hz, 2H), 7.69 – 7.63 (m, 2H), 7.61 (dd, *J* = 8.1, 1.9 Hz, 1H), 7.53 (t, *J* = 8 Hz, 2H), 7.26 (d, *J* = 8 Hz, 1 H), 3.88 (s, 3H), 2.63 (s, 3H). [13]C NMR (126 MHz, $CDCl_3$) δ 197.02, 164.29, 151.60, 144.07, 135.95, 133.78, 130.37, 128.90, 128.61, 122.94, 122.02, 111.44, 56.07, 26.62.

**b**

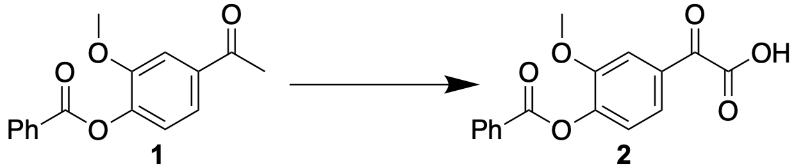

**1** → **2**

[1]H NMR (500 MHz, $CDCl_3$) δ 8.21 (td, *J* = 8.0, 1.6 Hz, 3H), 8.02 (d, *J* = 2.0 Hz, 1H), 7.71 – 7.64 (m, 1H), 7.54 (t, *J* = 7.3 Hz 1H), 7.34 (d, *J* = 8.3 Hz, 1H), 3.91 (s, 3H). [13]C NMR (126 MHz, $CDCl_3$) δ 183.07, 164.14, 161.50, 151.85, 146.31, 134.02, 130.45, 130.36, 128.69, 128.53, 125.69, 123.58, 113.89, 77.28, 77.02, 76.77, 56.19.

**c**

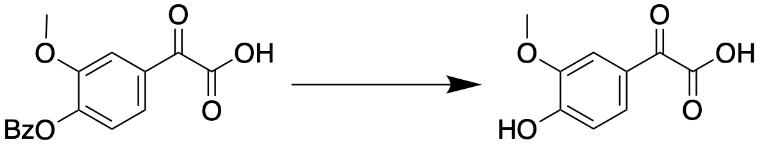

**2** → **VGA**

[1]H NMR (500 MHz, $D_2O$) δ 7.42 – 7.27 (m, 2H), 6.82 (d, *J* = 8.8 Hz, 1H), 3.74 (s, 3H). [13]C NMR (126 MHz, $D_2O$) δ 215.24, 193.54, 171.84, 152.06, 147.59, 126.30, 124.43, 115.17, 111.77, 55.62.

**Figure S11. Synthesis of (a) benzoyl acetovanillone (1) from acetovanillone (AV), (b) synthesis of benzoyl vanillyl glyoxylic acid (2) from (1), and synthesis of vanillyl glyoxylic acid (VGA) from (2).**

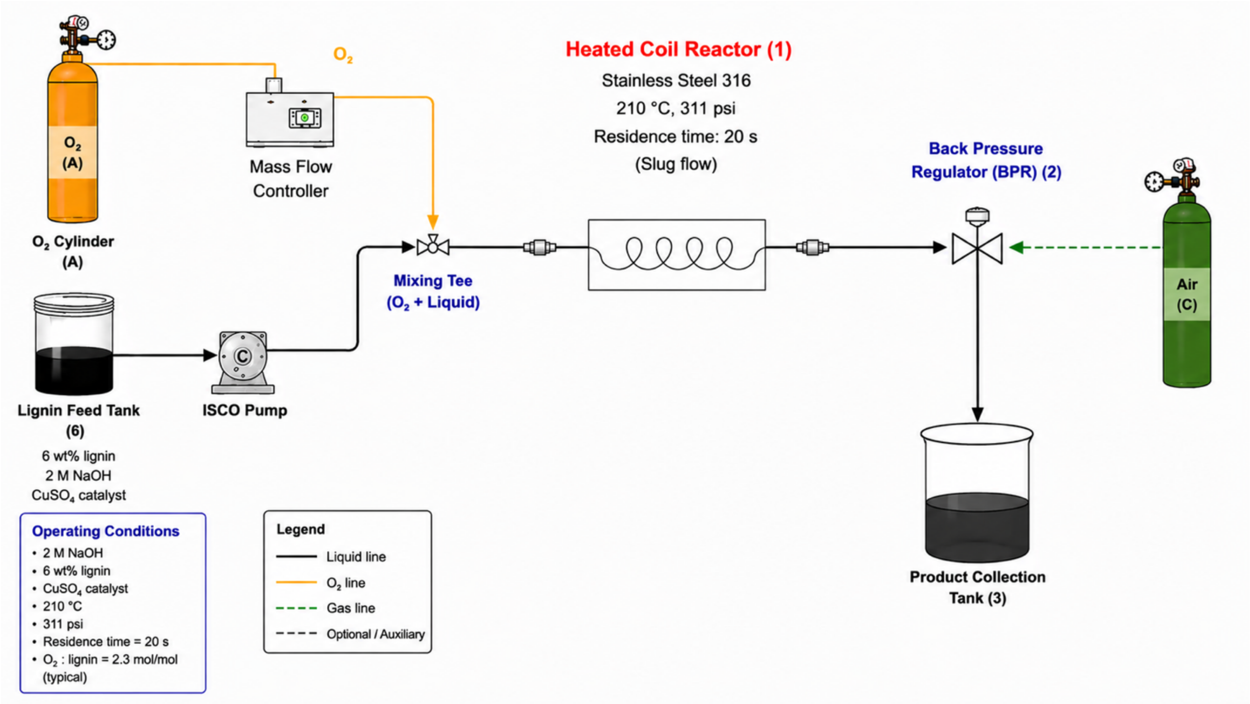

**Figure S12. Block flow diagram of continuous slug flow reactor.**

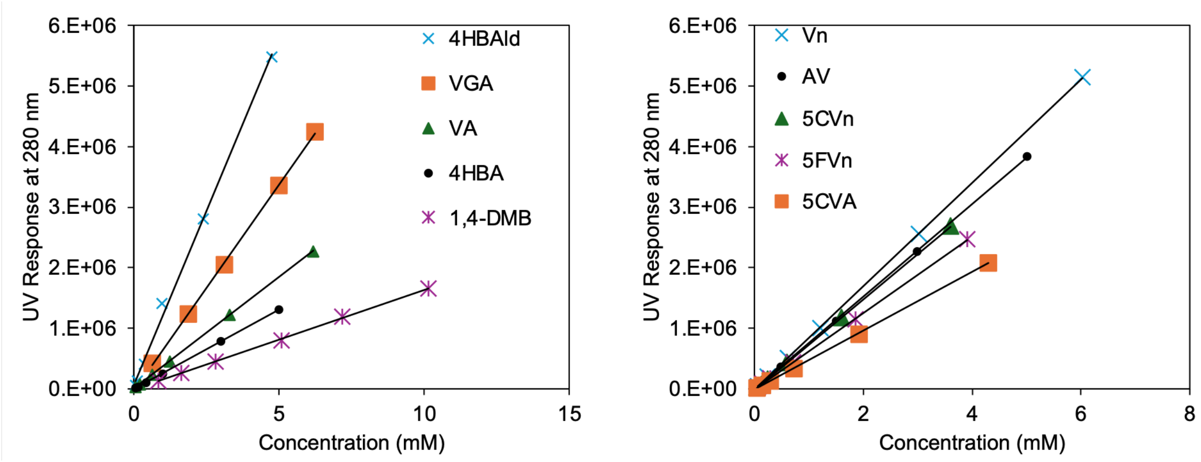

**Figure S13. Calibration curves for kraft-derived compounds studied in this manuscript as described in section 4.4.**

1,4-dimethoxybenzene (1,4-DMB) is the internal standard added to all samples.

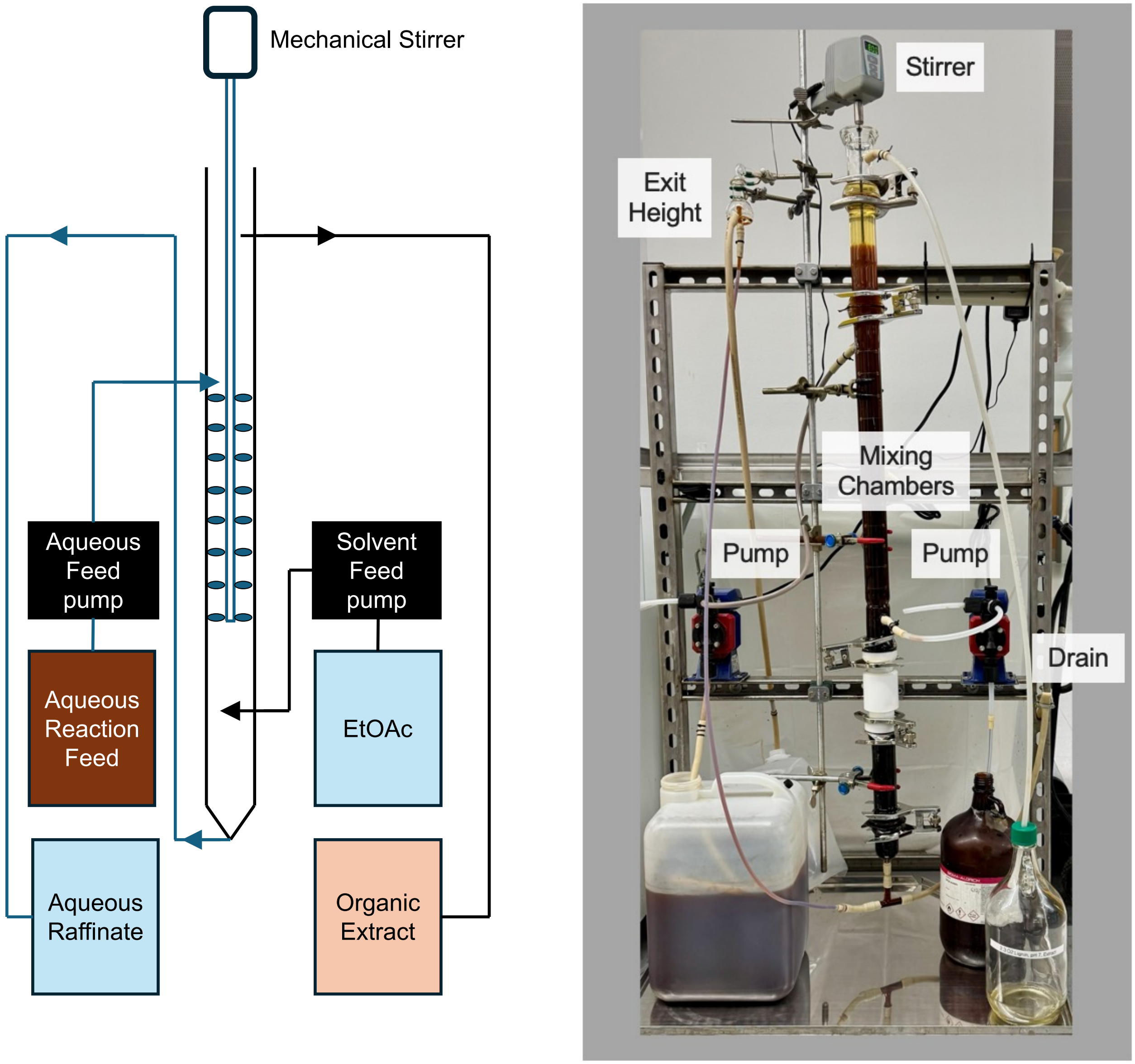

**Figure S14. Design of the counter current extractor.**

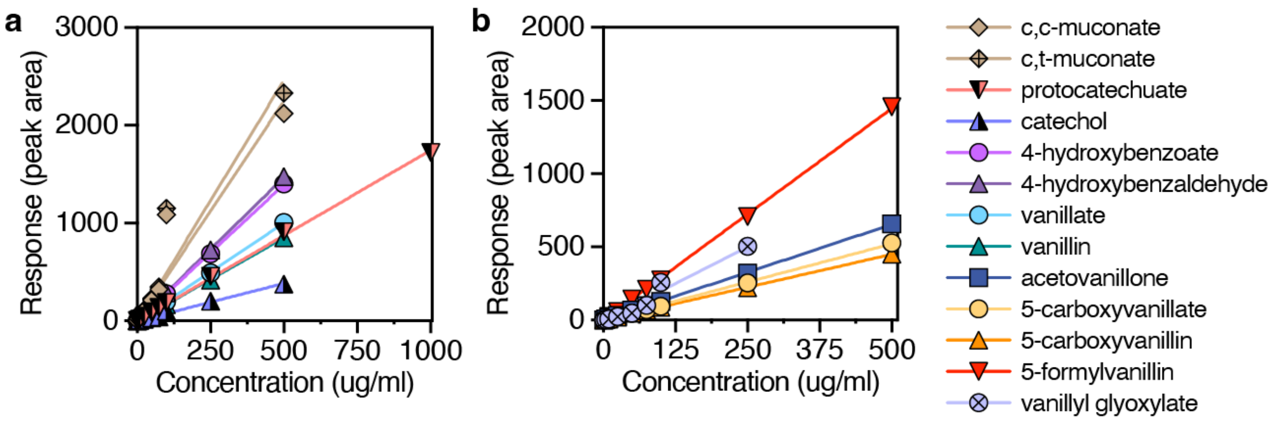

**Figure S15. Calibration curves for aromatic monomers, biological intermediates, and muconate via ultra-high-performance liquid chromatography with diode array detection (UHPLC-DAD) as described in section 4.10.**

Calibration curves for compounds that have been (a) described previously on protocols.io (Woodworth et al., 2024) and (b) that are new to this study.

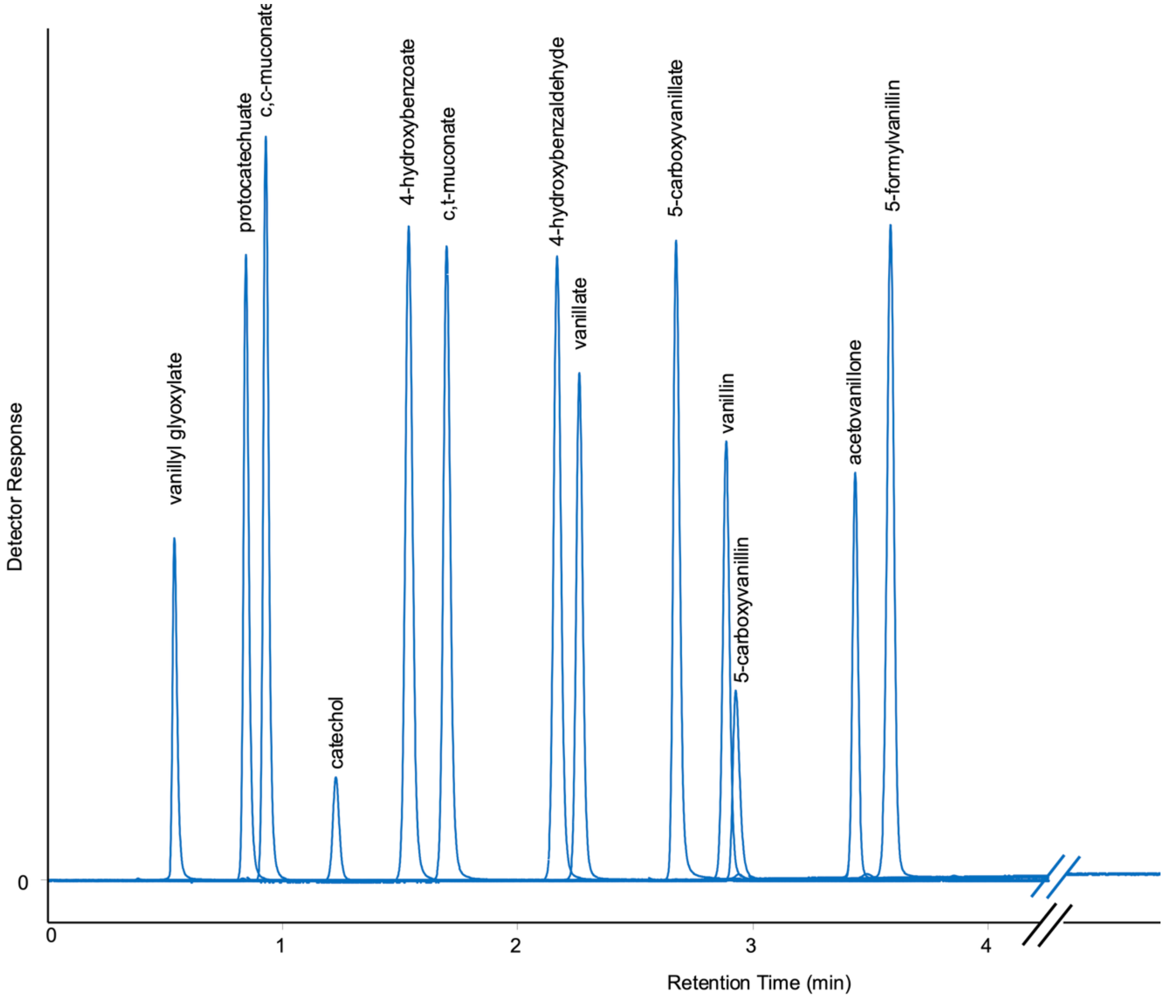

**Figure S16. UHPLC-DAD trace for standards of aromatic monomers, biological intermediates, and muconate as described in section 4.10.**

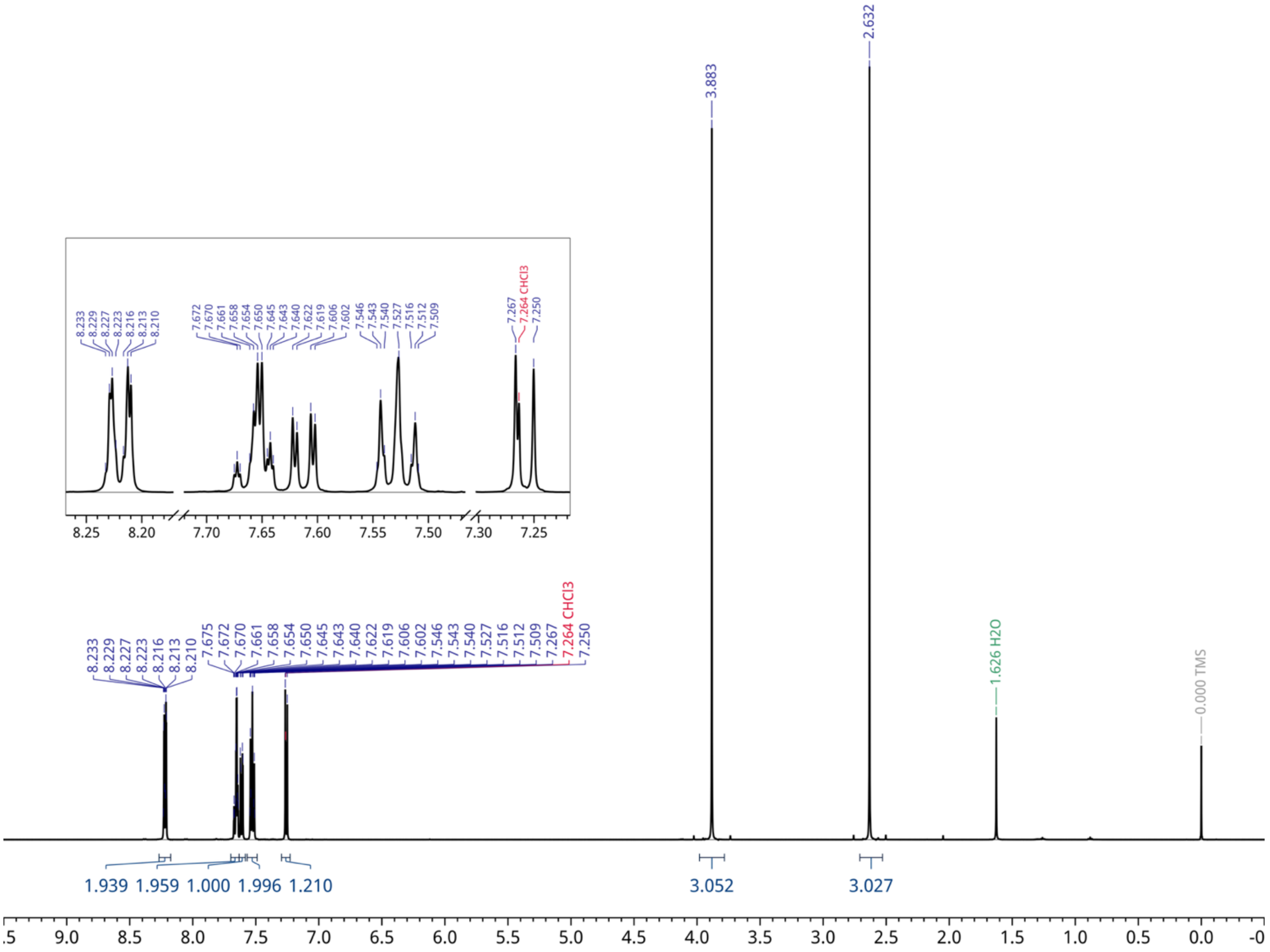

**Figure S17. $^1$H NMR (500 MHz) spectra of benzoyl acetovanillone in $CDCl_3$.**

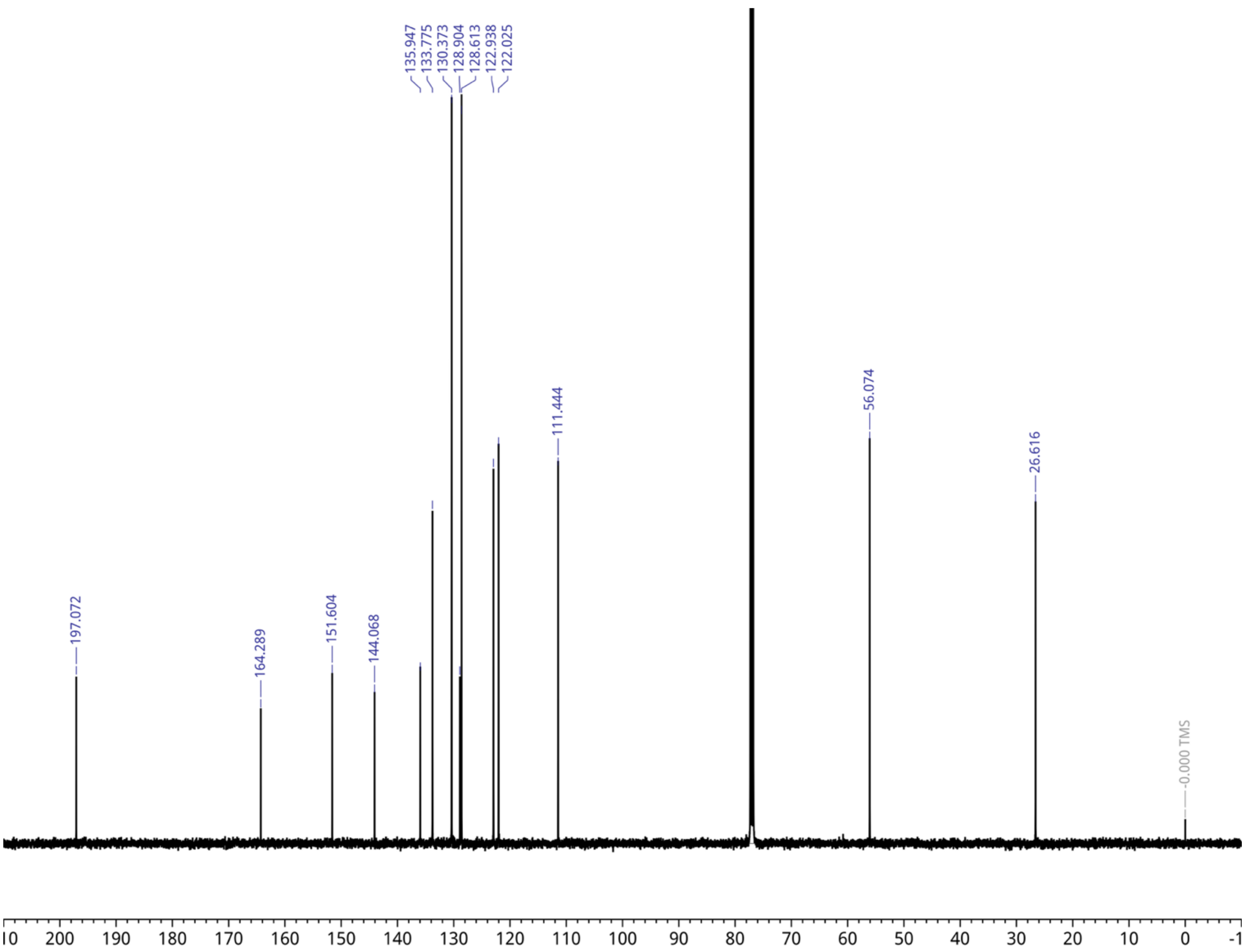

**Figure S18.** $^{13}$**C NMR (126 MHz) spectra of benzoyl acetovanillone (1) in** $CDCl_3$**.**

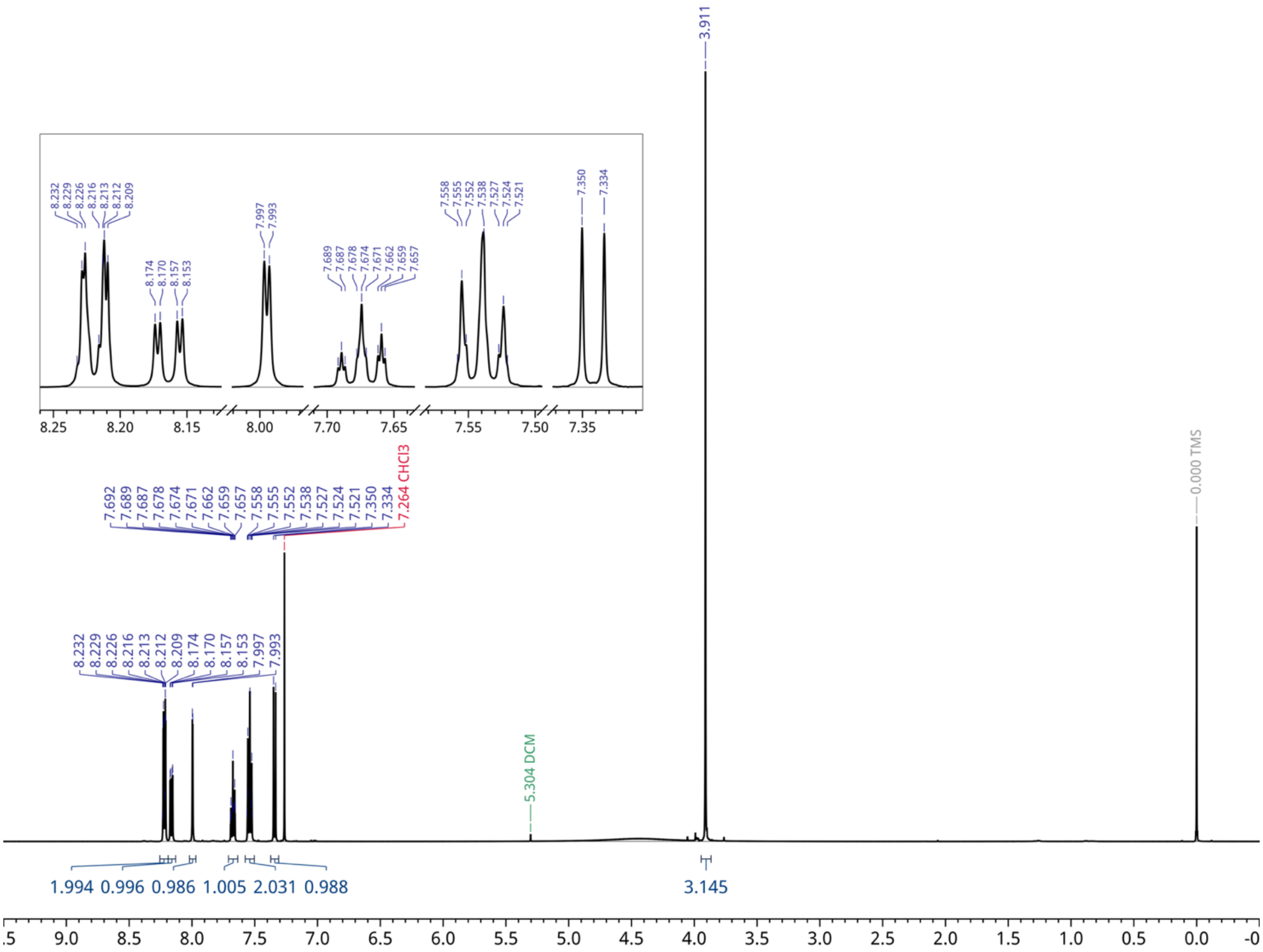

**Figure S19. $^1$H NMR (500 MHz) spectra of benzoyl vanillyl glyoxylic acid in $CDCl_3$.**

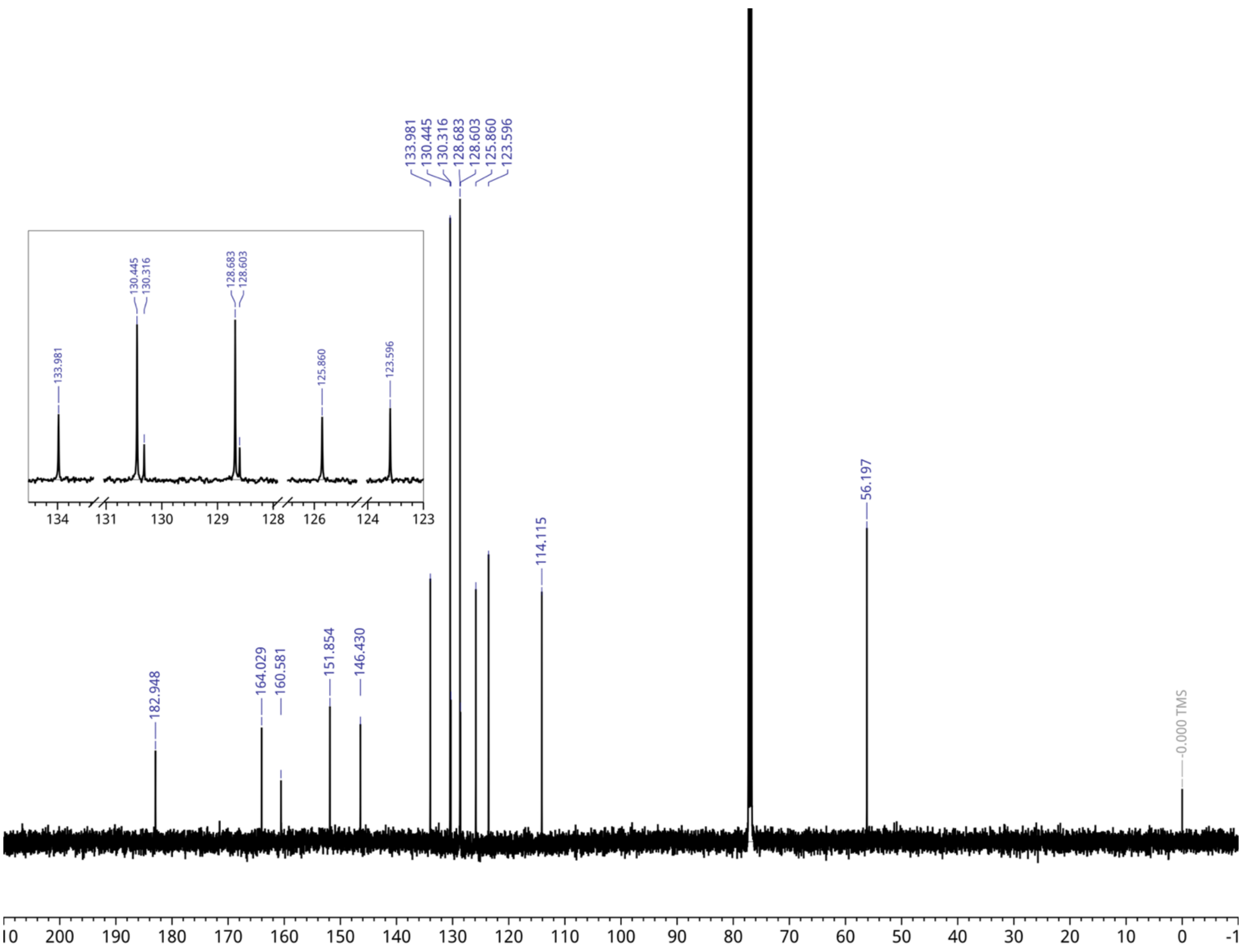

**Figure S20.** $^{13}$**C NMR (126 MHz) spectra of benzoyl vanillyl glyoxylic acid (2) in CDCl**$_3$**.**

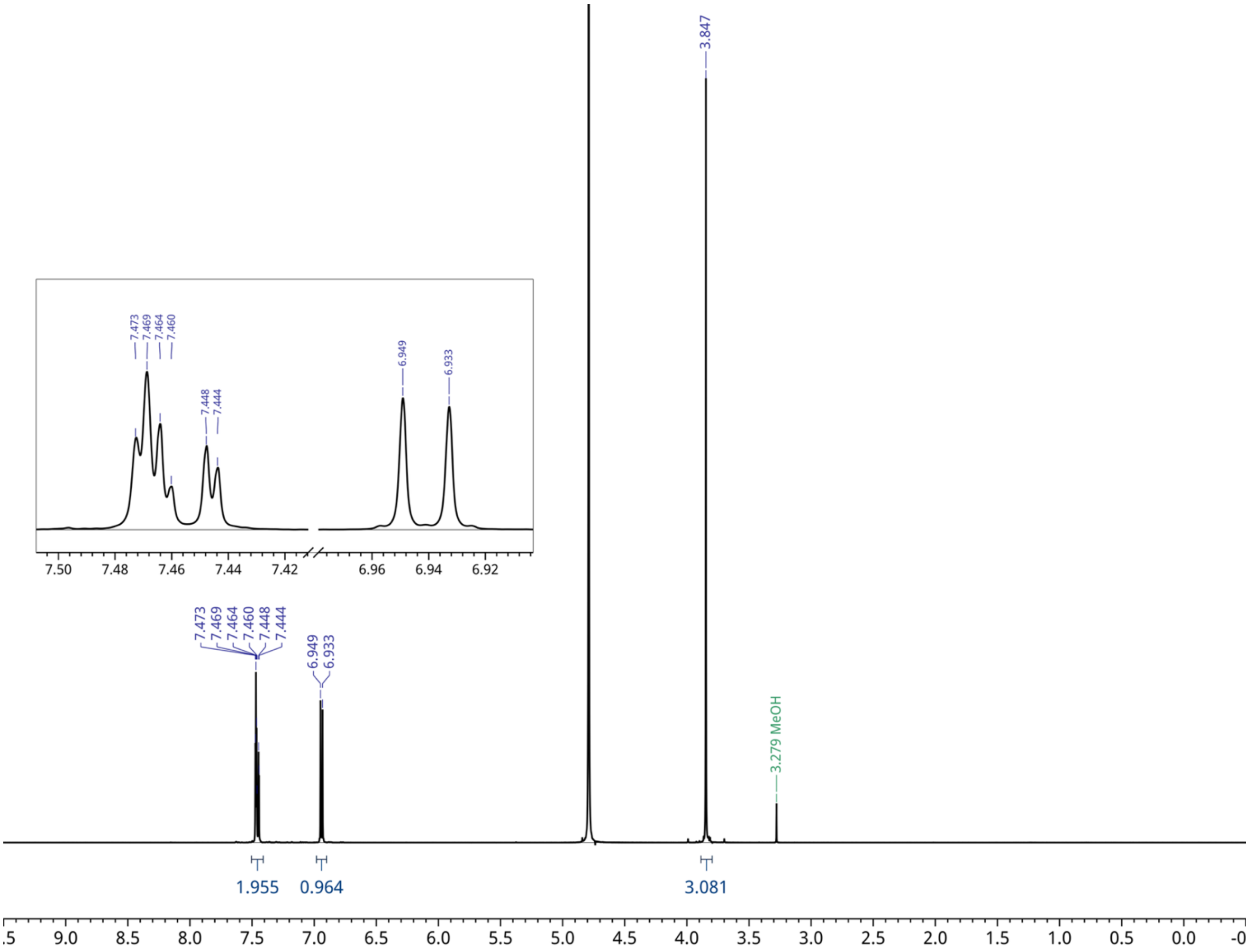

**Figure S21.** $^{1}$**H NMR (500 MHz) spectra of vanillyl glyoxylic acid (VGA) in** $CDCl_3$**.**

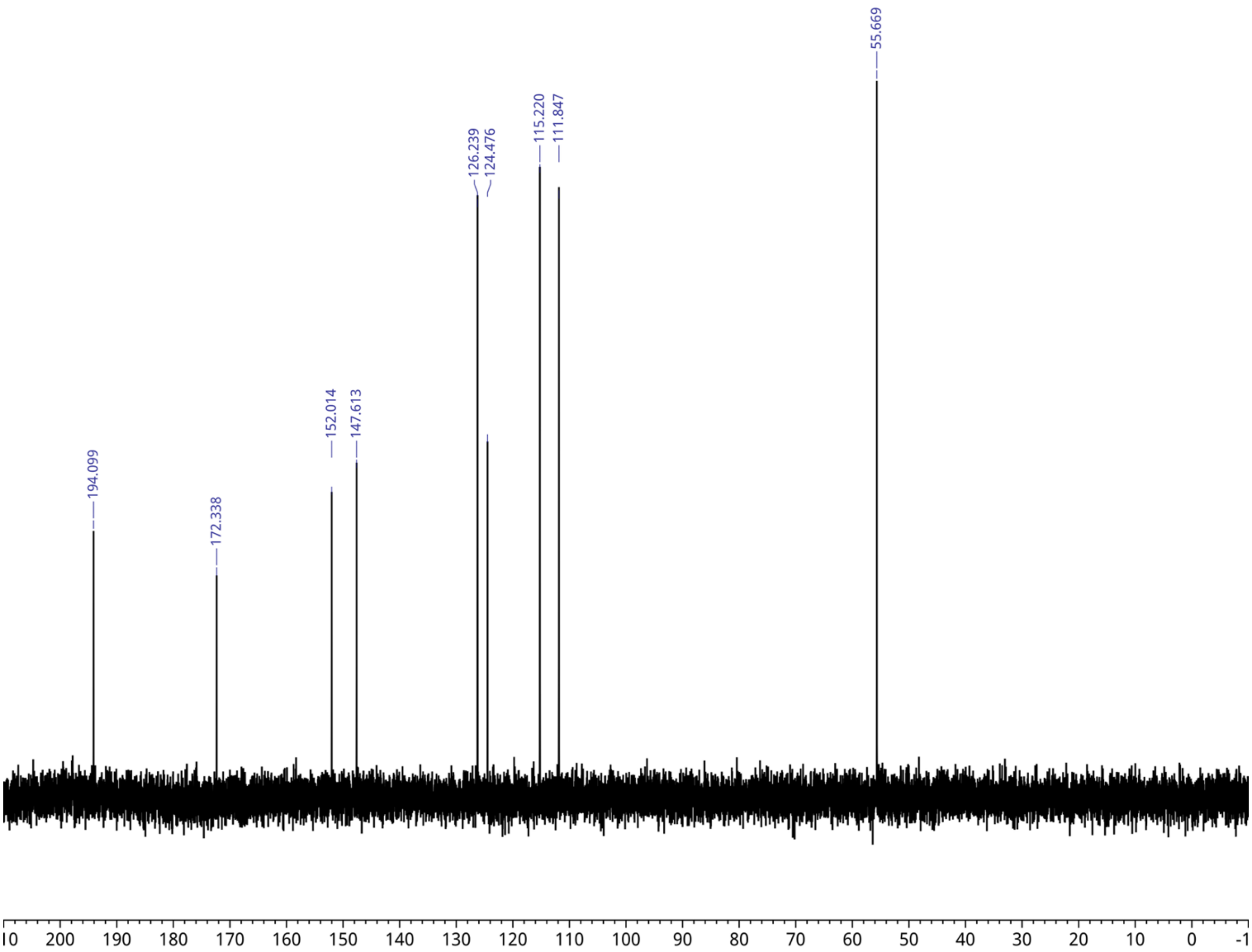

**Figure S22.** $^{13}C$ **NMR (126 MHz) spectra of benzoyl of vanillyl glyoxylic acid (VGA) in $CDCl_3$.**

### Supplemental Tables

**Table S1. Aromatic monomer yields in the reactor feed, depolymerized lignin (post-catalysis, prior to extraction), and oxidized lignin post extraction and rotary evaporation.** Weight% was calculated as g monomer / 60 g lignin. Yield for oxidized lignin extract was calculated based on an 8.11 wt% recovery of isolated oxidized lignin (post-catalysis, extraction, and rotary evaporation) from the initial lignin loaded into the reactor. Abbreviations: Abbreviations: VGA, vanillyl glyoxylate; 4HBA, 4-hydroxybenzoate; 4HBAld, 4-hydroxybenzaldehyde; VA, vanillate; 5CVA, 5-carboxyvanillate; 5CVn, 5-carboxyvanillin; Vn, vanillin; AV, acetovanillone; 5FVn, 5-formylvanillin; n.d., not detected.

| | VGA | 4HBA | 4HBAld | VA | Vn | 5CVn | 5CVA | AV | 5FVn |
|---|---|---|---|---|---|---|---|---|---|
| **Reactor Feed (g monomers/g kraft lignin)** | n.d. | n.d. | n.d. | 0.5% | 0.3% | n.d. | n.d. | 0.1% | n.d. |
| **Depolymerized kraft lignin yield (g monomer/g kraft lignin)** | 0.3% | 0.1% | 0.1% | 0.4% | 1.9% | 0.4% | 0.4% | 0.5% | 0.4% |
| **Oxidized lignin extract yield (g monomer/g kraft lignin)** | 0.09% | 0.02% | 0.05% | 0.12% | 0.57% | 0.21% | 0.15% | 0.11% | 0.07% |

**Table S2. Plasmids utilized in this study.**

| Plasmid | Description | Source |
|---|---|---|
| pK18msB | Suicide vector for kanamycin/sucrose selection and counterselection-mediated gene replacements in *P. putida* KT2440; confers kanamycin resistance | ATCC® 87097™; [12] |
| pK18sB | pK18mobsacB derivative that lacks oriT required for conjugation. | (Jayakody et al., 2018) |
| pJH402 | RV attP; kanR cargo vector | (Huenemann et al., 2025) |
| pGW32 | suicide vector expressing RV integrase | (Elmore et al., 2023) |
| pK18msBI | Suicide vector for kanamycin/sucrose selection and counterselection-mediated gene replacements in *P. putida* KT2440; confers kanamycin resistance. Derived from pK18msB but includes the lacI repressor for stronger repression of exogenous genes under control of promoters containing the lac operator (i.e., tac, lac, trc). | This work |
| pK51msBI | Suicide vector for kanamycin/sucrose selection and counterselection-mediated gene replacements in *P. putida* KT2440; confers kanamycin resistance. Derived from pK18msB but includes the strong endogenous promoter of *P. putida*, P51, to control expression of the sacB gene and the lacI repressor for stronger repression of exogenous genes under control of promoters containing the lac operator (i.e., tac, lac, trc). | (Mains et al., 2026) |
| pACB116 | pK18sB derivative for mutation of *PP_3493(P133L)* in *P. putida* KT2440 and derived strains. | (Bleem et al., 2024) |
| pKMM001 | pK18sB derivative for the integration of $P_{tac}$:$vceAB_{SYK-6}$:$acvF_{SYK-6}$ behind the *fpva* locus in *P. putida* KT2440 and derived strains. | This work |
| pKMM002 | pK18sB derivative for the integration of $P_{tac}$:$acvAB_{SYK-6}$ at the *PP_5042* safe integration site in *P. putida* KT2440 and derived strains. | This work |
| pKMM017 | pK18msBI derivative for the integration of $P_{tac}$:$acvCDE_{SYK-6}$ at the *PP_5322* safe integration site in *P. putida* KT2440 and derived strains. | This work |
| pKMM037 | pK51msBI derivative for the integration of $P_{tac}$:*vanAB* and simultaneous deletion of *crc* in *P. putida* KT2440 and derived strains. | This work |
| pKMM044 | pK51msBI derivative for the integration of $P_{tac}$:$vceAB_{SYK-6}$:$acvF_{SYK-6}$ and simultaneous deletion of *ampC* in *P. putida* KT2440 and derived strains. | This work |
| pKMM072 | pK51msBI derivative for the integration of $P_{tac}$:*catA2* and simultaneous deletion of *crc* in *P. putida* KT2440 and derived strains. | This work |
| pKMM073 | pK51msBI derivative for the integration of $P_{tac}$:$ligW2_{SYK-6}$ and simultaneous deletion of *crc* in *P. putida* KT2440 and derived strains. | This work |
| pKMM074 | pK51msBI derivative for the mutation of the *aroY* ribosome binding site in CJ781 and derived strains. | This work |
| pKMM075 | pK51msBI derivative for the integration of $P_{tac}$:$ligW2_{SYK-6}$:*catA2* and simultaneous deletion of *crc* in *P. putida* KT2440 and derived strains. | This work |
| pKMM111 | pK51msBI derivative for the integration of $P_{tac}$:$marK_{E16S\,Novosphingobium\ aromaticivorans\ DSM12444}$ at the *PP_5042* safe integration site in *P. putida* KT2440 and derived strains. | This work |
| pKMM118 | pJH402 derivative for the integration of $P_{tac}$:$mdlC_{PP}$ into the RV site in AG5577 and derived strains. | This work |
| pKMM119 | pJH402 derivative for the integration of $P_{tac}$:$dpgB_{ST-201}$ into the RV site in AG5577 and derived strains. | This work |
| pKMM120 | pJH402 derivative for the integration of $P_{tac}$:$EC844\_1132_{AC}$ into the RV site in AG5577 and derived strains. | This work |
| pKMM123 | pK18msB derivative for the integration of $P_{tac}$:$mdlC_{PP}$ and simultaneous deletion of *hsdRM* in *P. putida* KT2440 and derived strains. | This work |

**Table S3. Oligonucleotides utilized in this study.**

| Oligo Name | Sequence (5’ → 3’) |
|---|---|
| oKMM005 | GTTCATCCCGGTCGACAG |
| oKMM006 | CTGCTTCATTGAACTTTCATGAGCTTTC |
| oKMM008 | CAATGACCCCATTCGATCCGATAG |
| oKMM009 | GAATGCGTTCGAGGTCCACATTC |
| oKMM010 | GAAACGCGGTTGGAGTGC |
| oKMM013 | GTGCCGCTTTACCTGTTCTACTG |
| oKMM014 | CGACAGATCATAACTGTCAAAAAGCCAC |
| oKMM081 | CAACGGAGGATTGAATGCTCATG |
| oKMM082 | CAAACAGGATGGTGATGCGATC |
| oKMM172 | CCGCCATTTCCATGTGATAG |
| oKMM173 | AATGCTCGGCAGATTCAAAG |
| oKMM219 | CCGATAACTATGGAGGTCAGGTATGATTAC |
| oKMM237 | GAAATCACCGCACTGCTGAC |
| oKMM299 | CTCCTCGCTATCTACGATCCATTCTCTG |
| oKMM300 | GTTTTTAACGCCTGCCGGGTATTG |
| oACB423 | GAATGTCACCGCGGCAAAC |
| oACB424 | CCACAATAAATGGGCTAAACCGTG |
| oRW164 | GCAGATGGTCTCACTTGGTC |
| oRW165 | GTACACGTTCTGCAGGAAGC |

**Table S4. Strains utilized in this study.**

| Strain Name | Genotype | Construction details | Ref |
|---|---|---|---|
| NEB® 5-alpha F'$I^q$ *E. coli* | F′ *proA*$^+$*B*$^+$ *lacI*$^q$ *Δ(lacZ)M15 zzf::Tn10* (Tet$^R$) / *fhuA2Δ(argF-lacZ)U169 phoA glnV44 Φ80Δ(lacZ)M15 gyrA96 recA1 relA1 endA1 thi-1 hsdR17* | N/A | NEB Cat. C2992 |
| *P. putida* | Wild-type *Pseudomonas putida* KT2440 (KT2440) | N/A | ATCC® 47054 |
| *E. coli* WM6026 | lacI$^q$ , rrnB3, ΔlacZ4787, hsdR514, ΔaraBAD567, ΔrhaBAD568, rph-1, attλ::pAE12(ΔoriR6K cat::Frt5), ΔendA::Frt, uidA(ΔMluI)::pir, attHK::pJK1006 Δ(oriR6K- cat::Frt5;trfA::Frt) | N/A | Gift from Adam Guss |
| CJ781 | *P. putida* KT2440 Δ*catRBCA*::P$_{tac}$:*catA* Δ*pcaHG*::P$_{tac}$:aroY$_{KP}$:*ecdBD*$_{EC}$ Δ*pobAR* *fpvA*::P$_{tac}$:*praI*$_{JJ-1b}$:*vanAB* Δ*crc* | | (Kuatsjah et al., 2022) |
| AG5577 | mSAGE base strain; *P. putida* KT2440 Δ*PP_2876*::*R4_phiBT1_MR11_attB* cassette Δ*PP_4740*::*BxBI_RV_phi370_attB* cassette *PP_4217/4218* intergenic::*TG1_BL3_A118_attB* cassette | N/A | (Huenemann et al., 2025) |
| KMM002 | *P. putida* KT2440 *fpva*::P$_{tac}$:*vceA*$_{SYK-6}$:*vceB*$_{SYK-6}$:*acvF*$_{SYK-6}$ | pKMM001 was transformed into wildtype *P. putida* KT2440. Integration of *P*$_{tac}$*:VceAB*$_{SYK-6}$*:AcvF*$_{SYK-6}$ was confirmed by cPCR with oKMM005/oKMM008 and oKMM005/oKMM006. Oxford Nanopore sequencing (Plasmidsaurus, Inc.) on the linear amplicon verified the sequence. | This work |
| KMM011 | *P. putida* KT2440 *fpva*::P$_{tac}$:*vceA*$_{SYK-6}$:*vceB*$_{SYK-6}$:*acvF*$_{SYK-6}$ *PP_5042*::P$_{tac}$:*acvAB*$_{SYK-6}$ | pKMM002 was transformed into KMM002. Correct integration of *P*$_{tac}$*:AcvAB*$_{SYK-6}$ was confirmed by cPCR with oKMM009/oKMM010. Oxford Nanopore sequencing (Plasmidsaurus, Inc.) on the linear amplicon verified the sequence. | This work |
| KMM022 | *P. putida* KT2440 *fpva*::P$_{tac}$:*vceA*$_{SYK-6}$:*vceB*$_{SYK-6}$:*acvF*$_{SYK-6}$ *PP_5042*::P$_{tac}$:*acvAB*$_{SYK-6}$ *PP_5322*::P$_{tac}$:*acvCDE*$_{SYK-6}$ | pKMM017 was transformed into *E. coli* WM6026 and was conjugated into KMM011. Correct integration of *AcvCDE*$_{SYK-6}$ was confirmed by cPCR with oKMM013/oKMM014. Oxford Nanopore sequencing (Plasmidsaurus, Inc.) on the linear amplicon verified the sequence. | This work |
| KMM091 | *P. putida* KT2440 *fpva*::P$_{tac}$:*vceA*$_{SYK-6}$:*vceB*$_{SYK-6}$:*acvF*$_{SYK-6}$ *PP_5042*::P$_{tac}$:*acvAB*$_{SYK-6}$ *PP_5322*::P$_{tac}$:*acvCDE*$_{SYK-6}$ Δ*crc*:P$_{tac}$::*vanAB* | pKMM037 was transformed into *E. coli* WM6026 and was conjugated into KMM022. Deletion of *crc* and correct integration of P$_{tac}$::*vanAB* was confirmed by cPCR with oKMM172/oKMM173. Oxford Nanopore sequencing (Plasmidsaurus, Inc.) on the linear amplicon verified the sequence. | This work |
| MG079 | *P. putida* KT2440 *fpva*::P$_{tac}$:*vceA*$_{SYK-6}$:*vceB*$_{SYK-6}$:*acvF*$_{SYK-6}$ *PP_5042*::P$_{tac}$:*acvAB*$_{SYK-6}$ *PP_5322*::P$_{tac}$:*acvCDE*$_{SYK-6}$ Δ*crc*::P$_{tac}$:*ligW2*$_{SYK-6}$ | pKMM073 was transformed into *E. coli* WM6026 and was conjugated into KMM022. Correct integration of *P*$_{tac}$*:LigW2*$_{SYK-6}$ and deletion of *crc* was confirmed by cPCR with oKMM172/oKMM173. Oxford | This work |

| | | | |
|---|---|---|---|
| | | Nanopore sequencing (Plasmidsaurus, Inc.) on the linear amplicon verified the sequence. | |
| MG084 | *P. putida* KT2440 *fpva*::$P_{tac}$:*vceA*$_{SYK\text{-}6}$:*vceB*$_{SYK\text{-}6}$:*acvF*$_{SYK\text{-}6}$ *PP_5042*::$P_{tac}$:*markK*$^{E16K}$$_{Novosphingobium\ aromaticivorans}$ DSM12444 *PP_5322*::$P_{tac}$:*acvCDE*$_{SYK\text{-}6}$ Δ*crc*::$P_{tac}$:*ligW2*$_{SYK\text{-}6}$ | pKMM111 was transformed into *E. coli* WM6026 and was conjugated into MG079. Correct integration of *marK* E16K was confirmed by cPCR with oKMM009/oKMM010. Oxford Nanopore sequencing (Plasmidsaurus, Inc.) on the linear amplicon verified the sequence. | This work |
| MG005 | *P. putida* KT2440 Δ*catRBCA*::$P_{tac}$:*catA* Δ*pcaHG::*$P_{tac}$:*aroY*$_{KP}$:*ecdBD*$_{EC}$ Δ*pobAR* *fpvA*::$P_{tac}$:*praI*$_{JJ\text{-}1b}$:*vanAB* Δ*crc* *PP_3493(P133L)* | pACB116 was transformed into CJ781. The point mutation was confirmed by cPCR with oACB423/oACB424. Oxford Nanopore sequencing (Plasmidsaurus, Inc.) on the linear amplicon verified the sequence. | This work |
| MG027 | *P. putida* KT2440 Δ*catRBCA*::$P_{tac}$:*catA* Δ*pcaHG::*$P_{tac}$:*aroY*$_{KP}$:*ecdBD*$_{EC}$ Δ*pobAR* *fpvA*::$P_{tac}$:*praI*$_{JJ\text{-}1b}$:*vanAB* Δ*crc* *PP_3493(P133L)* *PP_5042*::$P_{tac}$:*acvAB*$_{SYK\text{-}6}$ | pKMM002 was transformed into MG005. The correct insertion was confirmed by cPCR with oKMM009/oKMM010. Oxford Nanopore sequencing (Plasmidsaurus, Inc.) on the linear amplicon verified the sequence. | This work |
| MG028 | *P. putida* KT2440 Δ*catRBCA*::$P_{tac}$:*catA* Δ*pcaHG::*$P_{tac}$:*aroY*$_{KP}$:*ecdBD*$_{EC}$ Δ*pobAR* *fpvA*::$P_{tac}$:*praI*$_{JJ\text{-}1b}$:*vanAB* Δ*crc* *PP_3493(P133L)* *PP_5042*::$P_{tac}$:*acvAB*$_{SYK\text{-}6}$ *PP_5322*::$P_{tac}$:*acvCDE*$_{SYK\text{-}6}$ | pKMM017 was transformed into *E. coli* WM6026 and was conjugated into MG027. Correct integration of *$P_{tac}$:acvCDE$_{SYK\text{-}6}$* was confirmed by cPCR with oKMM013/oKMM014. Oxford Nanopore sequencing (Plasmidsaurus, Inc.) on the linear amplicon verified the sequence. | This work |
| KMM140 | *P. putida* KT2440 Δ*catRBCA*::$P_{tac}$:*catA* Δ*pcaHG::*$P_{tac}$:*aroY*$_{KP}$:*ecdBD*$_{EC}$ Δ*pobAR* *fpvA*::$P_{tac}$:*praI*$_{JJ\text{-}1b}$:*vanAB* Δ*crc* *PP_3493(P133L)* *PP_5042*::$P_{tac}$:*acvAB*$_{SYK\text{-}6}$ *PP_5322*::$P_{tac}$:*acvCDE*$_{SYK\text{-}6}$ Δ*ampC*::$P_{tac}$:*vceAB*$_{SYK\text{-}6}$:*acvF*$_{SYK\text{-}6}$ | pKMM044 was transformed into *E. coli* WM6026 and was conjugated into MG028. Correct integration of *$P_{tac}$:VceAB$_{SYK\text{-}6}$:AcvF$_{SYK\text{-}6}$* and deletion of *ampC* was confirmed by cPCR with oKMM081/oKMM082. Oxford Nanopore sequencing (Plasmidsaurus, Inc.) on the linear amplicon verified the sequence. | This work |
| MG067 | *P. putida* KT2440 Δ*catRBCA*::$P_{tac}$:*catA* Δ*pcaHG::*$P_{tac}$:*aroY*$_{KP}$:*ecdBD*$_{EC}$ Δ*pobAR* *fpvA*::$P_{tac}$:*praI*$_{JJ\text{-}1b}$:*vanAB* Δ*crc*::$P_{tac}$:*catA2* *PP_3493(P133L)* *PP_5042*::$P_{tac}$:*acvAB*$_{SYK\text{-}6}$ *PP_5322*::$P_{tac}$:*acvCDE*$_{SYK\text{-}6}$ Δ*ampC*::$P_{tac}$:*vceAB*$_{SYK\text{-}6}$:*acvF*$_{SYK\text{-}6}$ | pKMM072 was transformed into *E. coli* WM6026 and was conjugated into KMM140. Correct integration of *$P_{tac}$:catA2* into the *crc* locus was confirmed by cPCR with oKMM172/oKMM173. Oxford Nanopore sequencing (Plasmidsaurus, Inc.) on the linear amplicon verified the sequence. | This work |
| MG074 | *P. putida* KT2440 Δ*catRBCA*::$P_{tac}$:*catA* Δ*pcaHG::*$P_{tac}$:*aroY**$_{KP}$:*ecdBD*$_{EC}$ Δ*pobAR* *fpvA*::$P_{tac}$:*praI*$_{JJ\text{-}1b}$:*vanAB* Δ*crc*::$P_{tac}$:*catA2* *PP_3493(P133L)* *PP_5042*::$P_{tac}$:*acvAB*$_{SYK\text{-}6}$ *PP_5322*::$P_{tac}$:*acvCDE*$_{SYK\text{-}6}$ Δ*ampC*::$P_{tac}$:*vceAB*$_{SYK\text{-}6}$:*acvF*$_{SYK\text{-}6}$ <br>*indicates RBS mutation | pKMM074 was transformed into *E. coli* WM6026 and was conjugated into MG067. The mutation of the *aroY* ribosome binding site (RBS) was confirmed by cPCR with oRW164/ oRW165. Oxford Nanopore | This work |

| | | | |
|---|---|---|---|
| | | sequencing (Plasmidsaurus, Inc.) on the linear amplicon verified the sequence. | |
| MG080 | *P. putida* KT2440 Δ*catRBCA*::$P_{tac}$:*catA* Δ*pcaHG*::$P_{tac}$:*aroY**$_{KP}$:*ecdBD*$_{EC}$ Δ*pobAR fpvA*::$P_{tac}$:*praI*$_{JJ-1b}$:*vanAB* Δ*crc*::$P_{tac}$:*ligW2*$_{SYK6}$:*catA2* *PP_3493(P133L) PP_5042*::$P_{tac}$:*acvAB*$_{SYK-6}$ *PP_5322*::$P_{tac}$:*acvCDE*$_{SYK-6}$ Δ*ampC*::$P_{tac}$:*vceAB*$_{SYK-6}$:*acvF*$_{SYK-6}$ <br>*indicates RBS mutation | pKMM075 was transformed into *E. coli* WM6026 and was conjugated into MG074. Correct integration of $P_{tac}$:*ligW2*$_{SYK-6}$:*catA2* into the *crc* locus was confirmed by cPCR with oKMM172/oKMM173. Oxford Nanopore sequencing (Plasmidsaurus, Inc.) on the linear amplicon verified the sequence. | This work |
| KMM326 | AG5577 *RV::*$P_{tac}$*: mdlC*$_{PP}$ | AG5577 was co-transformed with pGW032 and pKMM118 to integrate pKMM118 into the RV site of AG5577. Successful integration was confirmed via cPCR with oKMM219/oKMM237. Oxford Nanopore sequencing (Plasmidsaurus, Inc.) on the linear amplicon verified the sequence. | This work |
| KMM328 | AG5577 *RV::*$P_{tac}$*:dpgB*$_{ST-201}$ | AG5577 was co-transformed with pGW032 and pKMM119 to integrate pKMM119 into the RV site of AG5577. Successful integration was confirmed via cPCR with oKMM219/oKMM237. Oxford Nanopore sequencing (Plasmidsaurus, Inc.) on the linear amplicon verified the sequence. | This work |
| KMM330 | AG5577 *RV::*$P_{tac}$*:EC844_1132*$_{AC}$ | AG5577 was co-transformed with pGW032 and pKMM120 to integrate pKMM120 into the RV site of AG5577. Successful integration was confirmed via cPCR with oKMM219/oKMM237. Oxford Nanopore sequencing (Plasmidsaurus, Inc.) on the linear amplicon verified the sequence. | This work |
| KMM428 | *P. putida* KT2440 Δ*catRBCA*::$P_{tac}$:*catA* Δ*pcaHG*::$P_{tac}$:*aroY**$_{KP}$:*ecdBD*$_{EC}$ Δ*pobAR fpvA*::$P_{tac}$:*praI*$_{JJ-1b}$:*vanAB* Δ*crc*::$P_{tac}$:*ligW2*$_{SYK6}$:*catA2* *PP_3493(P133L) PP_5042*::$P_{tac}$:*acvAB*$_{SYK-6}$ *PP_5322*::$P_{tac}$:*acvCDE*$_{SYK-6}$ Δ*ampC*::$P_{tac}$:*vceAB*$_{SYK-6}$:*acvF*$_{SYK-6}$ Δ*hsdRM*::$P_{tac}$:*mdlC*$_{PP}$ <br>*indicates RBS mutation | MG080 was transformed with pKMM123 to simultaneously delete *hsdRM* and integrate $P_{tac}$:*mdlC*$_{PP}$. Correct integration was confirmed by cPCR with oKMM299/oKMM300. Oxford Nanopore sequencing (Plasmidsaurus, Inc.) on the linear amplicon verified the sequence. | This work |

**Table S5. Synthesized DNA utilized in this study.**

| Name | Relevant plasmid | Sequence (5'→3') | Description |
|---|---|---|---|
| *fpva*::$P_{tac}$:*vceAB*$_{SYK-6}$:*acvF*$_{SYK-6}$ | pKMM001 | aagccgaatgtcgatgatatctacaacctgagcgagcacgtcgagcaggaaggggcgtacgtcaccacccgcctga<br>acatcgccgacccgctgaaggtcatcctgggtgcacgcctggactggtacgacaacaagtcggtgtacagcgaaatc<br>aacgacggctactacaccaacagcgattacaaggtcacccgcaacgtcacccgctacgccggagtgatctacgacct<br>ggacgaccaccactcggtctacgccagctacaccgatatcttcatgccgcaatcggaactggcgcgtgaccgctccat<br>catccgcccaatcgaaggcaagaactacgagatcggcatcaagggcgagtacttcgacggcgcactcaacgccagc<br>gcggcgatcttccagatcgaccaggaaaaccgcgccgcagaagcttctaaccaggaaggttgcgtcgacatcacctg<br>ctacgaagcctcgggcaaggtacgcacccacggtatcgacctggagttgatgggcgcactgacccccaactggcaa<br>gtcggcgcaggctacacctactcgcaaaccaagtaccgcaaggatgccgacaagaacaaggaaggcaccaagttc<br>gacaccgacctgccagaacacctgttcaagctgagcaccacctacaccttgccgggcgagctgaaccagtggcgcgt<br>gggcggtaacgtgtatggccagagcagcatcttcaacaaaggcagcaacagcttcggcaactaccacatcgatcaag<br>gtgcatacgcggtagtgggcctgatggtcggctacaaggtcaacaagaacctcgacactcgcctgaacctcaacaac<br>gtattcgacaagaagtactaccagggcattgccagcaacaactcctggagcccgtacgacgtgtatggtgacccacgc<br>aacttcaccatcaccgccaagtacagcttctgatcgcctgacgttgaacgcaaaaaaccgcacccaggtgcggtttttt<br>GAGCTGTTGACAATTAATCATCGGCTCGTATAATGTGTGGAATTGTGAG<br>CGGATAACAATTTCACAC*GCGATTGTAAAAAGTTAAGTAACACACTAAGGA*<br>*GGTATTTTT*ATGGCCAAGACCTTCATCACCTGCGCGATCACCGGCGCCTC<br>CCCGATGCCCAAGCATCCCAATTTCCCCTTCCGTCCGGAACATGTCGCG<br>CAGGAAGCGCTCGACGCGGCGGCGGCCGGCGCCTCGATCATCCATGTC<br>CATGTGCGCAACGGCGCGGACGGCACGCCGAGCCAGGAACTTGAGGAT<br>TATCGCAAGGTCGTGGGCCTCATCCGCGAGAAGAACACCGACGTAATC<br>CTGAACGTCACCACCGGGCCCGGCTGCATGTGGTTCCCCAAGAGCGCC<br>GAGGAGCCGGCCGTGCCGGACATGGAAAAGACGCTGATGTTCACGGCC<br>GAGCGGCGCATCGAGCATATCCTCGAGCTCAAGCCGGACATGTGCACG<br>CTCGACATCTGCACGATGAACCTGTGGGGCGGCATCGCGATGAACCTG<br>GAGATGATCGTCGGCAAGATGGGCACCATGTTGCAGGATGCGGGCGTG<br>CTCACCGAGATCGAATGCTTCGAGGCCGGGGATTTCGTGTTCGCGGACG<br>ATCTGATGGCCAAGGGCCTCATCCCGAAGAACTCGCCCTTCACCTTCGT<br>GCTCGGCACGAAATACGGCCTGCCCGCCACGCCCGAGGCCATGATGTA<br>TTCCAGGAACCAGATCCCGCGCGGCGCGCACTTCACCGGTTTCGGCGTC<br>TCGCGGCACAGCTTCCCCATGGCCGCGCAGTCCGTGCTGCTCGGCGGGC<br>ACATGCGCGTCGGCTTCGAGGACACGATCTACCTGCGCAAGGGTGTGCT<br>CGCGAACAGCAATGCCGAGCTGGTCGAATGGGGCGCGGACATCGTGAA<br>CAAGCTGGGCGGTGAAGTGGCCACCCCGGGCGAGACGCGCGAGATGCT<br>CGGCCTCAAGGGCTGA*TCGGTAAGCCGTCTCGCGCCTGGGGTAAAACGAA*<br>*CCCAATAAGGAGGTTTTTTTT*ATGACCGACGAGCCGCGCGCGTTCACGCCT<br>TTCCTCTCCCGCCCCTATCGGATCGAATGGGGTCATTGCGACCCGGCGG<br>GCATCGTCTACGCGCCACGCTTCCTGGAGATGTTCGGCGAAAGCACCAT<br>CATGTTGTTCGAGAAGGCGCTGGGCGTGCGCAAGAGGGACATGGTGAA<br>GACGCGGGGCGTCGTCGGCTTCCCGATGGTCGATGTGTCGGCCCGTTTC<br>ATGCGCCCGGCCGCTTATGGCGACGATGTGGTGCTGGAAGTGGAAGCG<br>CCGGATTTCGGCAACTCCTCCTTCACCATCCGCCACCGCCTGCTGAAGG<br>ATGGGCACCCCTGCGTCGAGGGGACCGAGAAGCGTGTGTGGACCGTGC<br>GCGATGCCGAGCGTCCGGGCGGGATGCGCGCCGAGCGCGTGCCCGATG<br>ACATCCGGGCCATGTTCGGCTCCGCTGCCTGA*AAAAATTATCAAAAAGTAA*<br>*CAGGAGGGTCAGG*ATGACCGCACCCTTTACGGAGACGTTGCGGCCGGCA<br>CAGGGTATTATGTTCGATCTCGACGGCACCCTCATTTTGAGCGACCGGA<br>ACCTCGGTGGCTACAAGTTGCTCCCAGGGGCAGTCGAGTTGTTGTCCGA<br>GCTGGAGGCTTCGGGGTTCCCCTTCCTGGCCTTGACCAATGGCTCCGCA<br>TACCCCGCAGCGCAACAGGGGCCTCGCTTGCGCGCGCTGGGCTTGCCTA<br>TCCCAGACTCCCACTTGTTTACGCCCAACAGCGTGGCTGGCCATGTCTT<br>TCGGGAACGCGGCTTCGGTCGCGTCCTCGTGTTGGGGACCCAGGGCGTC<br>GTCGACGCCCTCACGGGTGAGGGGATTGCCACGGTGCGGCCGGGCGAA<br>GATGGTGCCACCAGCGCTGATGCCGTGTACGTGGCCTGGCACCCCGATT<br>GTGCAATGCCGGATATTCACGCGGCATGTGAGGCGGTCCTCGCTGGGG<br>CCGCTCTCTTTAGCGCTAGCGATGTGCCTTTTTTCGCGTCCCAATCCGGG<br>CGGGCCTTCGGGTATTCGTGCGCGATTGGGGGGGCAATCGCCCGCGTCA<br>CCGGCAAGGAGCCTGAACTCACCGGCAAACCTTCCCTCCACGCTATGA<br>ACTTTGTGGCTGCACGGCTGGGCGTCCCGATGGAGTCGATCGCCGTCAT<br>TGGGGATGATCCCAAAGTCGAAACCGAAATGGCGCGGGCCGGTGGGGC<br>TATGGGTATTGGGGTGACGACGGGGACCACCTCGGCACAAGAATGGGC<br>AGCCCAGCCGCCGGAGCGTCGTCCACACCGGATTATTGATGGTTTGGGT<br>GAATTGCGGGCTCTGGGGTTGCTCGGCTGA*agtcaaaagcctccgaccggaggcttttg*<br>*act*catggatgcctgaaaggctcccttacagatgctgctgcagtgcctcgaccaccatcgccaggccttcatcggcctc<br>ctcgatgatcgaacccatacgcatgaagtcatgggtcactcccgggatcaccgcaagctgcaccgccaccccggctt<br>gctccaagtgccgtgcataagcgacaccctggtcatgcagcgggtcgcactcggcaatcagcatcagtgcaggtgca<br>ctgttgctggcgacgctgcccagcaacggtgaaaaacgcggatcgtgacggtctgccggcaccgtggcgtattgctg | Homology arms are shown in red. $P_{tac}$ promoter shown in BLACK UNDERLINE. ORFs for *vceA*, *vceB*, and *acvF* codon optimized for *Pseudomonas putida* shown in BLUE, PURPLE, and ORANGE in that order. Synthetic ribosome binding sites for each gene shown in *ITALICIZED GRAY*. TonB terminator shown in *italicized black*. |

| | | | |
|---|---|---|---|
| | | gtagaaccactccagggtctgcgcttcaagcaggtaaccgctgccatagcgctgcaccgaaggccgccggcaactg<br>gcatcggtcaccgggtagatcattacctgcaggcgcggtgccggcagctcgcgctgcgcagccaactggttggcca<br>gaatggtagccaggctaccaccgacactgtcgcccaccaccgccagtcgctgcgcatcgatgcccagcgcctcggc<br>ctgctcgaccaaccagcgccaggcatccagggcatcgtcactggctgtcgggaaacgccactgcggcgccagccg<br>gtagcccacggcaatcaccggcaccccggcatcctgcgccaggttccagcacagcgtgtcatgcgaatcgaggctg<br>cccaccacgtagccgccgccatgcaggtacagcagcgccgcgccggccagtgcagggtcggcctgcggcgggcg<br>gtacaagcgcaccggcaaggtatggccatcgcgggtggtcaacgaaaggtcgctgatgcagtccggctcgtcggcc<br>ttgcccgcaatcagcgccgaggactcttcgaactggcggcgcgcctcgtccgccgccagggcatgcatgggcagca<br>ccttgccggcgctgcgtccggcctccaccagttgcaggtaggccgccaggtcagggttcagggacatcgttcgattct<br>ccagaggggc | |
| *PP_5042::P$_{tac}$:acvAB$_{SYK-6}$* | pKMM002 | ggcttcgaaccagtcatggtcgaaggcgtgctcggggtccttgccgactgcataggtcagcttgttcagtacagcagc<br>gcgaaaatccgcgacttcggcgtcacgtgctttaggttcctgggacatgcggcatcctcgggcaagttgacgaaagcg<br>gagggagagtgttgagactagacccttcgacagagagcgacagttacggttcgcagttttcatgcccactctgcgattc<br>cggcaaaaggttgttcataaattgaacaattccggtatcatcgcgcgccccaagaccgtgatccaccctataaagatga<br>aaacgaccctgatcgccgcggccgaagtcgaccgcctggagacctggcagcgctacaccagcaacatgtgccatgg<br>ctgccattcgacctgctgcaccctgccggtggaggtgaagatcaaggatctgatccgtatcggcgtggtcgacgagttc<br>gaaaaagacgaaccacccaagaacgtggccaagcgcctgcaaaaagaaggcatcatcgagcgcttcaaccagaag<br>tccgggatcttcaccctgacccggatgagcaacgacgactgcatgtacctggatcgtaaaagccggctgtgcaccattt<br>atgacaagcgcccggatacctgccgcaaccaccccaaggtcgggccacggccggggtattgcgcctacaagccca<br>aggtggttgggcgttgagtcttgcgccggggcataaatcgagcgccgcgcgggcggcgctcgattctgtggcgatgc<br>GAGCTGTTGACAATTAATCATCGGCTCGTATAATGTGTGGAATTGTGAG<br>CGGATAACAATTTCACAC*AACATACATCCAAGGAGAATATTATAT*ATGTCGGA<br>GCCTACGAAGGGCTTTCCGTTGCCCTCCTCGCTGGAAATTGTCCCTGGC<br>ACCGAGGCCGCACAGGCGGCCTACCCATATTACTGCCAATTCCATGATG<br>GTGATGACGAACGGTTCTGGTTTTACAACTCCATGCATTTCCCTGAGCC<br>CATGCATCACTTTGACATGATTACCGCGGAGGCAGCATACTGTGCTCTC<br>GGTAGCTTCAACACGCGCGTGCATGTCCTGCCTACGACGAAGGGTATCG<br>ACCATCGGATCATTAACGGCCGTGTCTTTATTGGGGGGTCGCTGTCAC<br>GGATCCGGCTGAAATTGAAGCTCGTGCCAAGGAATTCCAACAACGGGC<br>TTTCTACTACTATGAGCACTGGGAAGATTTGTACGCGCAGTGGAAAGAG<br>AAGATGAAGGCCTTGATCGCAGACGCCCAAGCGCTCTCGAAGCCTTCC<br>CTCCCTGACCTCGAGCCTATCGAGACGACCCATACCGGCAAGGGCGTG<br>GCAGCAAATCATGACCTCATCGATACGTACCGGAAAACGTTGGAGGGT<br>TACTTCCGGATGTGGCATCATCATTTCGAGTTCTTGTTGCTGGGCTATGG<br>TGCGTACCTCACCTTCTTTGATTTCTGCAAGAAGGCTTTTCCCGAAATTT<br>CGGATCAAGCAATTTCGCGTATGGTGGCAGGCATGGAAGCAGAGATTT<br>TCCGCCCCGACGAAGAGGTGAAACGGCTGGCCCGGCGTGCAGTCGAAT<br>TGGGGGTGGACGACCGCTTTCATGATGGCATGGTGATCGAGGATGTCCT<br>CACCGCTTTGGATCAACTGGGGGACAACGGGAAGGCGTGGTTGCGGGA<br>ATTGGAAACCTCGCGGAATCCTTGGTTTAACGTGAATGTCGGTGACGGG<br>TTCTACCACTATCACCGGAGCTGGAACGACGACTTGTCGATGCCGTTTG<br>CAGCACTCCCCGGTTACATTGCAAAGGTGAAGGCGGGTGCCGCGATTG<br>AGCGGCCGATGGAGCGTTTGATTGAAGAGCGTCAAGCGTTGATTGCCG<br>ACTATCGCGACCTCCTGGACACGGACGAGGATCGGGCGACGTACGATC<br>AACTCATCACGTTGGCGCACCGCGTGTTTCCATACGTGGAGGGCCATAA<br>ATTCTACTGTGAACACTGGTACACCAACTTGTTTTTTAATAAAATCCGG<br>GAGTTTGGTGCCCTCTTGCGGGACCATGGCTTCTGGGAAAAAGAAGAG<br>GATGTCTTCAATTTGACGCATTACGAGTTGGAGAGCGCGATTATCGACT<br>TGATGCTCGCCTGGTCCAGCGGTAGCGACCCGATCGGTCCACGTGTCTG<br>GCCTGCTAAGGTGGCCGAGCGTAAGGCGGCTATCAAGGCATGGGCGGG<br>GCATGTCGTGCCGCCTGCGCTGGGGCAAGTGCCGGATGTCATTGATGAT<br>CCTGCAATTGTCATGCTCTGGGGCATCACGCGCGAATCCCTGGATGTGT<br>GGCTGTCCGATGGCGAGGGTGATTCCAACGAACTCAAGGGTTTTGCTGC<br>GTCGTCGGGCGTGGTCGAAGGGACGGCCCGGGTCGTGCGCTCCGTGGA<br>AGAAATTAACCGGCTCCAACAGGGGGACATTCTCGTGTGTCAGGTCAC<br>GAACCCAACCTGGGCGCCGATCTTCCAAAAGATCGCTGCAGCGGTGTC<br>GGACATTGGCGGTTCGATGTCGCACATGGCGATCGTCGCTCGTGAGTAT<br>GGGCTGCCTGCTGTCGTCGGTACGGGGTCCGCTACCCAGAAGATCAAA<br>GATGGTCAACGTATTCGCGTGGACGGGGGCCGTGGCACGGTCACGATC<br>CTGGAGGAAGAACGTGTCTTGGAAGACGCATGA*AAGACGATATACACGA*<br>*TCCGATGAAGAGAAGTATTTTT*ATGACCGCTGCAGTGAGCTGGTTTCCTGA<br>CCTGTGGATCACCGACCGTCCAACCGTGGGCGGGAAAGGCGCGTCGCT<br>GGGTGAACTCACGCATGCTGGTATTGATGTCCCCCCTGGTTTCGTCGTG<br>ACGACGCAGGCCTTTGAGCTCTTTCTGGATGCATTGGAACGGAGCTCGC<br>CGGTGCGTGCGCTCGTCGAGACCTTGGATCCCAACGACCTCGATACCGT | Homology arms are shown in red. P$_{tac}$ promoter shown in BLACK UNDERLINE. ORFs for *acvA* and *acvB* codon optimized for *Pseudomonas putida* shown in BLUE and ORANGE in that order. Synthetic ribosome binding sites for each gene shown in *ITALICIZED GRAY*. TonB terminator shown in *italicized black*. |

| | | | |
|---|---|---|---|
| | | GTCCCGCATCTCCCGTTCGTTGCGGGTGCGGGTGGAGAATGAACCTTTG<br>CCAGCTATGGTGGAGACCGCAATTCTGGAAGCTCATGCGACGTTGGCC<br>AAAGACAACCCAGCCTTGACGGTCGCCGTGCGTAGCAGCGCTACGACG<br>GAAGATGCAGAAGATGCAAGCTTCGCAGGGCTCCAAGATACGTTTTTG<br>TGGGTGCCAGATGCTCAGGATGCCATCCACCGCGTCCGGGAATGCTGG<br>GGCAGCCTGTATAGCGTGGAGTCCATCTGTTATCGCCGGAAACACGGTC<br>TCCCGGAAGACGGCGTCGCCATGGCGGTGGTCGTGCAGACCATGGTGG<br>ACGCACGGACGGCAGGGGTGATGTTTACGCGTAGCCCCACCACCGGCG<br>ACAAATCCGTCATTACCATTGAGGGGGCTTGGGGGTTGGGCTCGGCAGT<br>GGTCTCCGGCGAGGTGACGCCGGATCGGTGGGTGATGGGGAAAATTAC<br>CGGTGAAATTTCGGTCCGCGACATTTCGGATAAACATATTCGTCAAGTG<br>CCTGCTGCGGGTGGCGGCATCGTCGATGAAGACGTGCCCGCTGAGCTG<br>CGTAAAACCCCGTGTCTGAGCGATGAAGAATTGCAAGCGTTGCGCGGC<br>GTCGGCCGTCGGATCGAAAAACACTATGGGCGGGCCCAGGATATCGAA<br>TGGGCGATCGATAAGTCGGGTGCTCTCTTGTTGTTGCAATCGCGTCCCG<br>AAACGGTGTGGTCGGCCAAAGAACAAAAACCGGTGGCCCCTGTCATTG<br>ATGATCCACTGAAGCACGTCATGACGATCTTCGGGGGGCGTCGGTAA*agt*<br>*caaaagcctccggtcggaggcttttgact*cgagctgggcatgaagcactgaacccaccgctcccatttcagccagcg<br>gggaagctccggcaagcaccccctgatccgctggtggatgctcaggataagtctgcggcaaaacgctgaccgggc<br>tggtgacaacgaaaccatcagcaaaccgcgcctctacgaaccgcgcctgccagccttgcgccggcggcgcccagtc<br>aaggctgaacgcctcactggggatgatcgtctgcgatgtgtagcgtaccccgcaggcatggcggaagtcccgatcca<br>aggggttagccgcagtccataccttcactgctacaggcatttccgaaagccgtaccagcacatgcgcagctccttcgcc<br>cggagccacttgcacgcgaggcatcggcaggctggcctggatacgctgaaccatcggcacgagggtagtgacaac<br>atgcgcacgcacgcccccatggtccgagttcggcagcaccctgaggctggtctggccaggcaggcgctgcaggtag<br>tccgtggcgtgatcaggggcgaagaaatcatccccgctggcggacaccaggtatttcggtatcttcaggcgagcaccc<br>ggctcatccaggtactgcatcgggtccatcagtgccatcaaccgagcgaaggccgggctgcccaactgctgcaatac<br>ctccgcagcctgataggaccaaagcgccaatggccatgaaccaccgtaccgcttgcgcaagccctccagcatgctgg<br>aagtgtcggcaacatcgatgacgctggg | |
| *PP_5322::P*$_{tac}$*:acvCDE*$_{SYK-6}$ | pKMM017 | caagcgaccaaggcatgcgcgtgctggcctcggccgtggaaatgggcgagctggaagtggtcgactgggccaact<br>cgcgtgaggacatgatcagcatcgacgcccatgcgcctctgaagcagatcctggccctggtgcgccgccacaagttc<br>agccgctacccggtgtacgacgccgagcgcgaagaattcaccggcctgctgcacatcaaggacctgctgctggagct<br>ggctgagctggagcacttgccagagaccatcgacctggaggaccttgcccggccactcgaacgcgtgtcgcggcac<br>atgccgctggcacagttgctggagcagttccgtaagggcggcgcgcacttcgtgctggtcgaggaagccgatggcaa<br>ggtgatcggctacctgaccatggaagacgtgctggaagtgctggtaggcgacatccaggacgagcaccgcaagacc<br>gagcgtggcattcttgcctaccagccgggcaagctgctggtgcgtggcgacaccccactgttcaaggtcgagcgcct<br>gctgggtatcgaccttgaccacatcgaggccgaaaccctggccgggttgatctacgagacgctcaagcgcgtgcctg<br>aagaagaggaagtgctggaggtcgaagggctgcgcatcatcatcaagaagatgaaaggccccaagatcgtactggc<br>caaggtgctgaagctggattgatggcccGAGCTGTTGACAATTAATCATCGGCTCGTAT<br>AATGTGTGGAATTGTGAGCGGATAACAATTTCACAC*TCAACAACTAATAT*<br>*CAAGATTAAGGGGAAAGGTTTTATT*ATGTCCCTCACCGCTAAGGACGTGGA<br>AGAGATCATGAAATTGCTGGAGGGGTCCACGTTTGATCGCTTGAGCTTG<br>GAGATGGACGGGATCCGGTTGGAACTGGAACGTGGCGGCGCGGGTGCA<br>ACACGTCCGGCACGTGCTACAGGTGCAGCTCCAGCTTCTCCTGCAGAAT<br>CTGCACCTGCACCGGCACCGGCACCTGCAGCACCTCCACCGGCAGCGG<br>CCCGCCCTGAACGTGAAGCTGGCCTGGTGGAGATTAACGCACCACTGCT<br>GGGGATTTTCTATCACGCACCCAAGCCCGGTGAAGCGCCTTTCGTCGAC<br>GTGGGGTCCCGTGTCGACACGGAAACGGTGATTGGCATTATCGAGGTG<br>ATGAAACTCATGAACTCCGTGTCGGCAGGGGTGTCGGGGGAAGTGGTC<br>GAGATCATCGGGCGGAATGGTGAATTGGTCGAACATGGGGAGGTGCTG<br>ATGCTCGTCCGCCCAGACGCGTGA*CTTTATAACCGAATATAAACCTAAGGA*<br>*CGTTTTTT*ATGACCATCCGGCGCATTCTGATTGCAAACCGGGGTGAGATC<br>GCGGTGCGGATTATTCGTACGGCACATAGCCTGGGCTTGGAAACGGTG<br>CTGGCGGTCTCCGAGGCTGATCGTGATTCCCTGGGGGCGCGCATGGCGA<br>CCCGTGCCTTGTGTATCGGTCCAGCCCGGCCAGGGGAATCCTACTTGCG<br>CGTGGAAACCATCGTGCAGGCGGCATTGGGTTCCGGCTGTGACGCAATT<br>CATCCGGGGTATGGCTTTCTGTCCGAGCGTGCCGCGTTGGCTAGCCTCT<br>GTGAAGCCGAGGGCGTGATTTTCATTGGGCCCACCGCGGCGCAGATCG<br>AAGCGGTCGGGGACAAACTCCGTGCACGTGCTGAAGCTATTGCCGCAG<br>ACGTGCCCGTCGTGCCCGGTGGTCCGGTGGATTCGCTGGCAGAAGCAC<br>AGGCATTGGCGGCCGGCATTGGGGCTCCATTGCTCATCAAAGCAGTTGG<br>TGGCGGTGGCGGTCGTATGAAACTGGTCGAAGACTTGGACGACCTCGC<br>GGCTACCATGGATATGGCCTCCGCAGAAGCAGGGGCAGCCTTCGGGGA<br>TGCTCGTGTCTACTTGGAGCGTTACGTGTCCCAAGGGCGTCATGTGGAA<br>GTCCAGGTGCTGGGGGATGGGGCTGGTAAAGTCATTCATCTGGGCGAG<br>CGGGATTGTAGCGTGCAACGTCGTTATCAGAAGCTGGTGGAAGAAACC<br>CCCGCGCCTCGGGTGCCAGATGCTGCGCGGGCTGCCATGCATGAGGCG<br>GCGATGCGCTTTGCTGCCCGCCTGGACTATCGCGGTGCGGGGACCGTCG<br>AATTCTTGTACGACATGGCCCGCGAACAATTTTACTTTTTGGAAATGAA<br>TGCCCGTATCCAAGTGGAGCACCCGGTCTCGGAAATGGTGACGGGTGT | Homology arms are shown in red. P$_{tac}$ promoter shown in BLACK UNDERLINE. ORFs for *acvC*, *acvD*, and *acvE* codon optimized for *Pseudomonas putida* shown in BLUE, PURPLE, and ORANGE in that order. Synthetic ribosome binding sites for each gene shown in *ITALICIZED GRAY*. TonB terminator shown in *italicized black*. |

| | | | |
|---|---|---|---|
| | | GGACTTGATCGCGGAACAAATCGCGATTGCTGACGGTCAAGGTCTCCG<br>GCTGGCACAGGGTGACGTCGCAATTGACGGTGCTGCTATTGAATGTCGC<br>ATCAATGCAGAAGATCCTGCCCGCGATTTTATGCCGAGCCCTGGCACGG<br>TGTCCCGCGCACACTGGCCGGCGGGCGAGGGCATCCGCGTGGATACCC<br>ATATTGTTGATGGTGCAAAAATTCCACCGTTTTATGATTCTATGATTGCC<br>AAAGTGATTGCACATGGTCCGGATCGTGCTACAGCATTGGCTCGTCTGC<br>AAGCTGCTTTGCGTGAAACACGTATTGAAGGTGTTTCTACAAATTTACG<br>TTTCCAAGCAGAAATTCTGGCAGATCCGGATTTCGCAGCAGGTGGCGTT<br>GATACAGGTTTTCTCGCTCGTCGCGCAGCACGCCAAGCAGAACGTACA<br>GAAACGTGA*AATTTCGATAATTTTTTAAAACTGTTGGATTATATTTT*ATGGCG<br>GATATTAAGTTGGTGGAGACCTCCCTGCGCGATGGTAATCAATGCCTCT<br>GGGGGGCCCTCGGTGTGGACACGGCCCGGACCTTGAGCATTGCTCCTGT<br>GCTGGAACGTGTGGGGTTCAAGGCTATCGATTTTACGACCTCGACCCAC<br>ATGGGCGTCGCGGTCCGGTATAAGCAGGAGGACCCCTGGGAACGCATC<br>CGTCTGATGAAACAAGCATGCCCAACGACGCCCTTGCAGTTTCTGTCGA<br>CGGGGTTTCGGTTCATTGCGTGGGAAACCGCTTCGCCAGATTTCATGGC<br>GTTGGCGTTTGCTACCTTGATGAAAAACGGTATTGGTCGTTTTGCTCTG<br>GCCGACCCAATGAACGACGCCGAAGCTAATGTGCAAGTCGCCCGCATG<br>ATTAAACGGGACGGCGACGCTTACGTCGTCGGGGCCCTGGTGTTTACGC<br>TGAGCCCAATCCATGATGATGCGCACTATGCCCAGGCCGCCGCCACCAT<br>GGCGGCGTCGCCTGACATCGATGCACTGTACATTAAGGATCCAGGGGG<br>CTTGTTGTCGCCCCAACGCGCGCGGACGCTGATCCCCGCAATCCTCGCA<br>GTCATTGGGGATAAGCCGTTGGAGCTCCATGCGCATTGCACGATCGGCC<br>TGGGGGAGCAATCGTATTACGAGGCTGCGGGCTTGGGGGTGGAGGCTC<br>TGCAAGGTGCCTCCGGCGGCTTGGGCGACGGTACCTCGAATCCACCACT<br>CGAACGCGTCGTCGCCAACCTGCGTGCATTGGGGCATAGCGTGGACAT<br>CGACGACGCAGCGCTGGCAGAAGCTGGGCGTTACTTTACCGATCTGGCT<br>CAAGCAGAAGGCTTGCCGACCGGGGCGCCAATGCCATTCGATGCTGCT<br>TACCTCCAGCATCAACTGCCAGGCGGCATGGTCGGGACGATGCGCCGG<br>CATCTCGCCGATCATCGCGTCCCCCACCTCGAGGGTGCAGTGGTCGAGG<br>AATTGGGGCGTGTCCGTGAAGAGCTGGGTTGGCCTATCGTGATGACCCC<br>CTTCGCCCAGATGGTGATGACCCAAGCTGTCATGAACGTCACGGGCACC<br>GAACGCTACTCGGTCATCCCCGATGAAGTCATTCGCTACGCCATTGGGC<br>GGTTCGGCCGCCCCAACCAGCCTATTGCACCAAATGTCCTGGACCGGAT<br>TATGGCGCTGCCACGTACGAAAGAATTGGCTGCAGAACCTGGGATGGC<br>TGACCTGCCTGACCTGCGGAAGCGGATCGGGACCCATTTGCCGGATGA<br>GGAATTCTTGTTGCGTGCCACCATGCCCGCGGGGCAAGTCGACGCTATG<br>CAAGCTGCAGGGCCCGCACCACGGCATTATGACCCTACCTTGCGCCCCG<br>CCATGGCATTGATCCGCAAACTGCTCGCGCGCACGGATATTGACCGCGT<br>CTCGGTCGAAAAGGCAGGCTTCCGGTTGGAACTCGAATCGCACGGGTG<br>A*agtcaaaagcctccggtcggaggcttttgact*aatcggtcaggggttgccggcagtgaaatccggcaacccgctg<br>atcggcctgtcgaagcggtaaggtatcgactccagcgccaaccccacattgcgctgcaccacaaaatgcaggtgcgg<br>cccagtgctgttgccggtattgcccgacttggccagcatctggccctgccgtacccgttgcccctccgccaccaccacc<br>gaaccacgcatcaggtgaaggtatacgcccatggtgccatccgggtgcaaaatcctgacaaagttacccgaggggtt<br>ggggccgcgcccgctctggctgttctgcgtcttcaccaccaccccttcccgcgctgcgatgatgggcgtgccttcggg<br>catggcgatatccatggcgtagcgccctttgggtccaaaatggctgacgcggccattggggccctgagtgacgcgga<br>acggcccgccaatccacgggaacgcatagcggaagccctggggccgtttgcccgggtcacccatggcctgcagca<br>gggtcgaacggtagttcagcagcagcccgggccgccccctgagcacactgagcatctgcgtggtgcgcggcggca<br>gcagccggcgcacgatacgcggggcatcgccgccttgtgcgttgaccaggttgtccagccgcaactctacctctacc<br>ggcacatgcagctcgtttcgtgccgaaaagctcacgccaccgtcgaaggttttcgcgtt | |
| Δ*crc*::P*tac*:*vanAB* | pKMM037 | gttatggccctcctcctcccttcaagtcctaccatacaagcgggtttacccgcgaaagggccagatcaggcggcacaa<br>aaaaaggcgaccctctgaagaaggtcgcctttttcattcctgggccaggatcagatcccgtactgcgcccggtaggcct<br>cgacagccggcagatgctgtttcagctgtggatcgtcagccaggaactccagcacctgggtcagcgaaacaatgctg<br>accaccgggataccgaagtcgcgttccacttcctggatcgccgacagttcgccattgccgcgctcttcacggttcagcg<br>cgatcagtacaccagcggccttggcctgctgggcgttgatgatctgcatgacctcacgaatggcggtaccggcagtaa<br>tcacgtcgtcgatgatcagcacgtcaccggccagcggcgcgccgaccaggctgccaccttcgccatggtccttggctt<br>ccttgcggttgaagcaccatggcacgtccagctgatgctggtcggcaagggccacggcggtagtcgccgccaaagg<br>gatacccttgtaggccgggccgaacagcacatcgaacgggatcttgctatcgacgatggcggcggcgtagcaacgc<br>cccagctcagccagtgcggagccggtgttgaacaagccggcattgaagaaatacgggctggtacgccccgatttcag<br>ggtgaattcaccgaaacgcagtaccccgcgatcgatggcaaaacggataaagtcgcgctgatacggctgcatgaata<br>gtcccggacaccacggatttagctaaatgggttgagctcgggtatcatacacgcacgagatttatggggccatttgaatt<br>cGAGCTGTTGACAATTAATCATCGGCTCGTATAATGTGTGGAATTGTGA<br>GCGGATAACAATTTCACACtctaga*GAGGAGGACAGCT*ATGTACCCCAAAA<br>ACACCTGGTACGTCGCCTGCACCCCCGATGAGATCGCCACCAAACCCCT<br>GGGCCGGCAGATCTGCGGGGAAAAAATCGTGTTCTACCGCGCCCGCGA<br>GAACCAAGTAGCCGCCGTCGAGGACTTCTGCCCGCACCGCGGCGCACC<br>GTTGTCGTTGGGCTATGTCGAGGACGGCAACCTGGTGTGCGGCTACCAC<br>GGCCTGGTGATGGGTTGCGACGGCAAGACCGTGTCGATGCCGGGCCAA<br>CGGGTGCGTGGCTTCCCCTGCAACAAGACCTTTGCGGCCGTCGAGCGCT | Homology arms are shown in red. P$_{tac}$ promoter shown in BLACK UNDERLINE. ORFs for *vanA* shown in BLUE and *vanB* shown in ORANGE, in that order. Synthetic ribosome binding sites for *vanA* shown in *ITALICIZED GRAY*. TonB terminator shown in *italicized black*. |

| | | | |
|---|---|---|---|
| | | ATGGCTTCATCTGGGTCTGGCCCGGTGACCAGGCGCAGGCCGACCCGG<br>CGCTGATTCCGCATCTGGAATGGGCGGTGAGTGATGAGTGGGCCTACG<br>GCGGCGGGCTGTTCCACATCGGTTGCGACTACCGCCTGATGATCGACAA<br>CCTCATGGACCTCACCCATGAAACCTATGTGCACGCCTCCAGCATCGGC<br>CAGAAGGAGATCGACGAGGCACCGCCGGTCACCACCGTCACCGGCGAC<br>GAAGTGGTCACCGCCCGGCACATGGAAAACATCATGGCGCCACCGTTC<br>TGGCGCATGGCCTTGCGTGGCAATGGCCTGGCCGACGATGTACCAGTG<br>GACCGCTGGCAGATCTGCCGTTTCACCCCACCTAGCCATGTGCTGATCG<br>AAGTGGGTGTAGCGCATGCCGGCAAGGGCGGCTACCACGCCGAGGCAC<br>AGCATAAGGCGTCGAGCATCGTGGTCGACTTCATCACCCCTGAGAGCG<br>ATACCTCTATCTGGTACTTCTGGGGCATGGCGCGCAACTTCGCTGCGCA<br>CGACCAGACCCTGACCGACAACATTCGTGAGGGCCAGGGCAAGATTTT<br>CAGCGAAGACCTGGAAATGCTCGAACGCCAGCAGCAGAACCTGCTGGC<br>CCACCCCGAGCGCAACTTGCTGAAGCTGAATATCGACGCCGGCGGCGT<br>GCAGTCACGCAAAGTGCTGGAGCGGATCATCGCCCAAGAGCGTGCGCC<br>GCAGCCGCAACTGATCGCCACCAGCGCCAACCCTGCCTGAggaacagccgac<br>ATGATCGATGCCGTAGTGGTATCCCGTAACGATGAAGCGCAGGGTATCT<br>GCAGCTTCGAGCTGGCCGCGGCAGATGGCAGCCTGCTGCCGGCGTTCA<br>GCGCCGGCGCCCATATCGACGTGCACCTGCCCGACGGGCTGGTGCGCC<br>AGTATTCGCTGTGCAACCACCCCGAAGAACGCCATCGCTATCTGATTGG<br>CGTACTCAACGACCCGGCTTCGCGGGGCGGTTCTCGTAGCCTGCACGAA<br>CAGGTGCAAGCCGGTGCCCGGCTGCGTATCAGTGCGCCGCGCAACCTG<br>TTCCCGCTGGCCGAGGGTGCGCAGCGCAGTTTGCTGTTTGCTGGCGGTA<br>TCGGCATTACCCCAATCCTGTGCATGGCCGAGCAGCTGTCCGACAGCGG<br>CCAGGCCTTCGAGCTGCACTACTGTGCCCGCTCCAGCGAGCGTGCGGCG<br>TTTGTCGAGCGGATCCGCAGCGCGCCGTTCGCTGATCGGCTGTTCGTGC<br>ATTTTGACGAGCAGCCGGAAACGGCGCTGGACATCGCCCAGGTGCTGG<br>GCAACCCGCAAGATGATGTGCACCTGTATGTATGCGGGCCCGGCGGGT<br>TCATGCAGCATGTGCTGGACAGCGCGAAGGGGCTGGGCTGGCAGGAGG<br>CCAACCTGCACCGCGAGTACTTCGCCGCAGCACCGGTGGATGCCAGCA<br>ACGATGGCAGTTTCGCGGTGCAGGTGGGCAGCACGGGACAGGTGTTCG<br>AGGTGCCAGCCGACCGGACCGTGGTGCAGGTGCTGGAAGAGAATGGTA<br>TCGAGATCGCCATGTCGTGCGAGCAGGGTATTTGCGGCACCTGCCTGAC<br>ACGCGTGCTGCAGGGCACACCGGACCATCGCGATCTGTTTCTCACCGAA<br>GAGGAACAGGCCCTGAACGATCAGTTCACGCCCTGCTGCTCGCGCTCG<br>AAGACGCCGCTGCTGGTGCTGGACATCTGAgatatcattcagactagt*agtcaaaagcct*<br>*ccgaccggaggcttttgact*aaggccattggggctgcattgcagccccaatggttatttgatcggccgccagatcatcg<br>ggtagcgataaggcttgccctcattcgcccgcaccgccgcgatgatgatcaatatcatcactgccaccgtgagcatgg<br>caaacagcaccagcccgataaacacgaacatcagcaggaagcagatgaacccggcaatggtcacgctgatctgaaa<br>gttcagcgcctctttgccctgggcgtcgataaaagggtcttgctcgcgcttgaggtgccatagcaccagcgggcccag<br>caggtgcccaagcggcactaccaggcccaacagcgcggagaggtggcagaacatcgcccattgccggatttcggc<br>attgggcggggtgatcgacaggttcgaatcgctcatgccccgcacctccttggccggtttcagtcagccagtgctgcct<br>gctgcagctcgaaaatctcgttcatgcccttctgcgccaaagcgagcatggcgttgaaatcttccggctggaacggtgc<br>gccttcggcagtaccctgcacttcgatgaaaccaccagcgctggtcatgaccacattcaggtcggtttcggctgcggag<br>tcttccaggtagtccaggtcgagcaccgcttcgccctgatacatgcctaccgacacggcagcgatcatgtgcttgagcg<br>ggttgccgcccttgaggccgccgcgcttcttgatcactgccaaagcgtcgcacaaggcaaccatggcgccggtgatg<br>gacgcggtacgggtaccgccatcggcctggatcacgtcgcaatcgacgtacagggtgatgtcaccgagcttgctcat<br>gtccagcgcggcgc | |
| *ΔampC*::P$_{tac}$:*vceAB*$_{SYK-6}$:*acvF*$_{SYK-6}$ | pKMM044 | cttgcctctgccggaaacccgtctgcagcggcgccaggcatgaaaaagcccggcctatgcaggccgggctctggta<br>agggtgcgaaccctcaattcagcggttcgatcaccttgactgcctgctgctcgcgggcagcgggccattcttgttgcag<br>ggtctgcagcacgcggccaacgaattcagtgtcggcggcggctttcttgccgacgtaaccttggccacggcggtaga<br>ccttgaagcgggcacgcatgctgggcaggtgctcttcaaattcatcttcgacggtgttcggtggcaggcggtcgaggc<br>gcacttcaccggtttggctaacccacagcaaatggtcgtcaaggctgtccttgcgcacggcgaacagctgggccaatt<br>ggtcgatagtcggattgttgttcaagttcatcataaagcccccattacccattcataggtgattttcacagttggcgctatca<br>cggtgtagcgcactaccaaggctgctacagcagcatcgcaacaggggctttctcacgctgccttttggacagtttccctc<br>tccacggacggcctgcgttaccgcgcaggccgtctggaacacccttgccaccgctcccgatcggggagacggcttca<br>tcatgcccaagcgcgatgaacggcgtcaaccattttgtagtgaatatttttgctcactacattctgtcgttttgtccgacaat<br>caacgcgagcgttaGAGCTGTTGACAATTAATCATCGGCTCGTATAATGTGTGG<br>AATTGTGAGCGGATAACAATTTCACAC*GCGATTGTAAAAAGTTAAGTAACA*<br>*CACTAAGGAGGTATTTTT*ATGGCCAAGACCTTCATCACCTGCGCGATCAC<br>CGGCGCCTCCCCGATGCCCAAGCATCCCAATTTCCCCTTCCGTCCGGAA<br>CATGTCGCGCAGGAAGCGCTCGACGCGGCGGCGGCCGGCGCCTCGATC<br>ATCCATGTCCATGTGCGCAACGGCGCGGACGGCACGCCGAGCCAGGAA<br>CTTGAGGATTATCGCAAGGTCGTGGGCCTCATCCGCGAGAAGAACACC<br>GACGTAATCCTGAACGTCACCACCGGGCCCGGCTGCATGTGGTTCCCCA<br>AGAGCGCCGAGGAGCCGGCCGTGCCGGACATGGAAAAGACGCTGATGT<br>TCACGGCCGAGCGGCGCATCGAGCATATCCTCGAGCTCAAGCCGGACA<br>TGTGCACGCTCGACATCTGCACGATGAACCTGTGGGGCGGCATCGCGAT<br>GAACCTGGAGATGATCGTCGGCAAGATGGGCACCATGTTGCAGGATGC | Homology arms are shown in red. P$_{tac}$ promoter shown in BLACK UNDERLINE. ORFs for *vceA*, *vceB*, and *acvF* codon optimized for *Pseudomonas putida* shown in BLUE, PURPLE, and ORANGE in that order. Synthetic ribosome binding sites for each gene shown in *ITALICIZED GRAY*. TonB terminator |

| | | | |
|---|---|---|---|
| | | GGGCGTGCTCACCGAGATCGAATGCTTCGAGGCCGGGGATTTCGTGTTC<br>GCGGACGATCTGATGGCCAAGGGCCTCATCCCGAAGAACTCGCCCTTC<br>ACCTTCGTGCTCGGCACGAAATACGGCCTGCCCGCCACGCCCGAGGCC<br>ATGATGTATTCCAGGAACCAGATCCCGCGCGGCGCGCACTTCACCGGTT<br>TCGGCGTCTCGCGGCACAGCTTCCCCATGGCCGCGCAGTCCGTGCTGCT<br>CGGCGGGCACATGCGCGTCGGCTTCGAGGACACGATCTACCTGCGCAA<br>GGGTGTGCTCGCGAACAGCAATGCCGAGCTGGTCGAATGGGGCGCGGA<br>CATCGTGAACAAGCTGGGCGGTGAAGTGGCCACCCCGGGCGAGACGCG<br>CGAGATGCTCGGCCTCAAGGGCTGA*TCGGTAAGCCGTCTCGCGCCTGGG*<br>*GTAAAACGAACCCAATAAGGAGGTTTTTTTT*ATGACCGACGAGCCGCGCGC<br>GTTCACGCCTTTCCTCTCCCGCCCCTATCGGATCGAATGGGGTCATTGC<br>GACCCGGCGGGCATCGTCTACGCGCCACGCTTCCTGGAGATGTTCGGCG<br>AAAGCACCATCATGTTGTTCGAGAAGGCGCTGGGCGTGCGCAAGAGGG<br>ACATGGTGAAGACGCGGGGCGTCGTCGGCTTCCCGATGGTCGATGTGTC<br>GGCCCGTTTCATGCGCCCGGCCGCTTATGGCGACGATGTGGTGCTGGAA<br>GTGGAAGCGCCGGATTTCGGCAACTCCTCCTTCACCATCCGCCACCGCC<br>TGCTGAAGGATGGGCACCCCTGCGTCGAGGGGACCGAGAAGCGTGTGT<br>GGACCGTGCGCGATGCCGAGCGTCCGGGCGGGATGCGCGCCGAGCGCG<br>TGCCCGATGACATCCGGGCCATGTTCGGCTCCGCTGCCTGA*AAAAATTAT*<br>*CAAAAAGTAACAGGAGGGTCAGG*ATGACCGCACCCTTTACGGAGACGTTG<br>CGGCCGGCACAGGGTATTATGTTCGATCTCGACGGCACCCTCATTTTGA<br>GCGACCGGAACCTCGGTGGCTACAAGTTGCTCCCAGGGGCAGTCGAGT<br>TGTTGTCCGAGCTGGAGGCTTCGGGGTTCCCCTTCCTGGCCTTGACCAA<br>TGGCTCCGCATACCCCGCAGCGCAACAGGGGCCTCGCTTGCGCGCGCTG<br>GGCTTGCCTATCCCAGACTCCCACTTGTTTACGCCCAACAGCGTGGCTG<br>GCCATGTCTTTCGGGAACGCGGCTTCGGTCGCGTCCTCGTGTTGGGGAC<br>CCAGGGCGTCGTCGACGCCCTCACGGGTGAGGGGATTGCCACGGTGCG<br>GCCGGGCGAAGATGGTGCCACCAGCGCTGATGCCGTGTACGTGGCCTG<br>GCACCCCGATTGTGCAATGCCGGATATTCACGCGGCATGTGAGGCGGTC<br>CTCGCTGGGGCCGCTCTCTTTAGCGCTAGCGATGTGCCTTTTTTCGCGTC<br>CCAATCCGGGCGGGCCTTCGGGTATTCGTGCGCGATTGGGGGGGCAAT<br>CGCCCGCGTCACCGGCAAGGAGCCTGAACTCACCGGCAAACCTTCCCTC<br>CACGCTATGAACTTTGTGGCTGCACGGCTGGGCGTCCCGATGGAGTCGA<br>TCGCCGTCATTGGGGATGATCCCAAAGTCGAAACCGAAATGGCGCGGG<br>CCGGTGGGGCTATGGGTATTGGGGTGACGACGGGGACCACCTCGGCAC<br>AAGAATGGGCAGCCCAGCCGCCGGAGCGTCGTCCACACCGGATTATTG<br>ATGGTTTGGGTGAATTGCGGGCTCTGGGGTTGCTCGGCTGA*agtcaaaagcct*<br>*ccgaccggaggcttttgact*ggtgagtccttttgtggagcgtgtccgttgagcgataccactcggggttccggcgttag<br>cgacggggagccaaagaccttatcgtctgcccacccaccatgacaagatggatacagtgatgaagtatttgcgcatgtt<br>atttgacaatttcaccctggccttgctcggtgtggtactcatcgccaccgtactgccttgctcgggcgatggcgcggtgta<br>tttcggctggctgaccaacctggccattggcctgttgttcttcctgcatggcgccaaactgtcccgtgaagccatcattgc<br>cggtgccgggcactggcgcctgcacctactggtgttctcctgtactttcgtattgttcccgctgctgggcctggcgttcaa<br>acctctgttcgtaccgctggtgggtaacgagctttatctgggcatcctgtacctgtgtgccttgcctgcaaccgtgcagtc<br>ggccattgcctttacctcgctggcccgcggtaacgtgccagcggccatctgcagcgcggcggcctccagcctgctgg<br>gtatcttcctcaccccgttgctggtgatgctgctactgggcgccggtggtgatacaggttccggcctggatgcggtgttg<br>aagatcactttgcagctgctggtgccgttcgttgccgggcaggtcgcgcggcgctggatcggcgcctgggtcaagcg<br>caatgcgcgctggctcaaggtggtggaccagggttcgatcctgctggtggtgtacaccgccttcagtgaggccgtggt<br>tac | shown in *italicized black*. |
| *$\Delta crc::P_{tac}:catA2$* | pKMM072 | gtagcgtagtgtgacttgaagggcacaccctggcgcaggcttgcttctattgtcgcttgctcgcgaggcattccttatcg<br>gcgcttgggtgttgaaaggaggaggggtgatttctcctacggtctgtactgcgcgaacagtcagaagcgagttatggc<br>cctcctcctcccttcaagtcctaccatacaagcgggtttacccgcgaaagggccagatcaggcggcacaaaaaaagg<br>cgaccctctgaagaaggtcgcctttttcattcctgggccaggatcagatcccgtactgcgcccggtaggcctcgacagc<br>cggcagatgctgtttcagctgtggatcgtcagccaggaactccagcacctgggtcagcgaaacaatgctgaccaccg<br>ggataccgaagtcgcgttccacttcctggatcgccgacagttcgccattgccgcgctcttcacggttcagcgcgatcag<br>tacaccagcggccttggcctgctgggcgttgatgatctgcatgacctcacgaatggcggtaccggcagtaatcacgtc<br>gtcgatgatcagcacgtcaccggccagcggcgcgccgaccaggctgccaccttcgccatggtccttggcttccttgcg<br>gttgaagcaccatggcacgtccagctgatgctggtcggcaagggccacggcggtagtcgccgccaaagggataccc<br>ttgtaggccgggccgaacagcacatcgaacgggatcttgctatcgacgatggcggcggcgtagcaacgccccagct<br>cagccagtgcggagccggtgttgaacaagccggcattgaagaaatacgggctggtacgccccgatttcagggtgaat<br>tcaccgaaacgcagtaccccgcgatcgatggcaaaacggataaagtcgcgctgatacggctgcatgaatagtcccgg<br>acaccacggatttagctaaatgggttgagctcgggtatcatacacgcacgagatttatggggccatttGAGCTGT<br>TGACAATTAATCATCGGCTCGTATAATGTGTGGAATTGTGAGCGGATAA<br>CAATTTCACAC*GCTCCCCAATAGCGATCGAGACTTTTAT*ATGACCGTGAAC<br>ATTTCCCATACTGCCGAGGTACAGCAGTTCTTCGAGCAGGCCGCAGGCT<br>TTTGTAATGCGGCCGGCAACCCACGCCTCAAACGCATCGTGCAGCGCCT<br>GCTGCAGGATACCGCGCGGCTGATCGAAGACCTGGACATCAGCGAAGA<br>CGAGTTCTGGCACGCCGTCGATTACCTCAACCGCCTGGGCGGTCGCGGC<br>GAAGCCGGGTTGCTGGTGGCGGGGCTGGGCATCGAACACTTCCTCGAC<br>CTGCTGCAGGATGCCAAGGACCAGGAGGCAGGGCGCGTTGGCGGCACC | Homology arms are shown in red. $P_{tac}$ promoter shown in BLACK UNDERLINE. ORFs for *catA2* shown in BLUE. Synthetic ribosome binding sites for each gene shown in *ITALICIZED GRAY*. TonB terminator shown in *italicized black*. |

| | | | |
|---|---|---|---|
| | | CCACGCACCATCGAAGGCCCGTTGTACGTGGCTGGCGCACCGATTGCCC<br>AAGGTGAAGTGCGCATGGACGACGGCAGCGAGGAGGGCGTGGCCACG<br>GTGATGTTCCTGGAAGGCCAGGTGCTGGACCCGCACGGACGCCCGCTG<br>CCGGGTGCCACGGTCGACCTGTGGCATGCCAATACCCGTGGTACCTACT<br>CGTTCTTCGACCAAAGCCAGTCGGCGTACAACCTGCGTCGGCGCATCGT<br>TACCGATGCCCAGGGGCGCTACCGCGCGCGCTCCATCGTGCCATCGGGC<br>TATGGCTGCGACCCGCAGGGGCCAACCCAGGAATGCCTGGACCTGCTG<br>GGCCGTCATGGCCAGCGCCCGGCGCACGTGCACTTCTTTATCTCGGCCC<br>CAGGGTACCGGCACCTGACCACGCAGATAAACCTGTCGGGGGACAAGT<br>ACCTGTGGGATGACTTTGCCTATGCCACACGGGATGGGCTGGTCGGGG<br>AGGTGGTGTTCGTCGAAGGGCCGGATGGTCGGCATGCCGAGCTGAAGT<br>TCGACTTCCAGTTGCAGCAGGCCCAGGGCGGTGCCGATGAGCAGCGCA<br>GCGGGCGGCCGCGAGCTTTGCAGGAGGCCTGA*agtcaaaagcctccgaccggaggc*<br>*tttttact*aaggccattggggctgcattgcagccccaatggttatttgatcggccgccagatcatcgggtagcgataagg<br>cttgccctcattcgcccgcaccgccgcgatgatgatcaatatcatcactgccaccgtgagcatggcaaacagcaccag<br>cccgataaacacgaacatcagcaggaagcagatgaacccggcaatggtcacgctgatctgaaagttcagcgcctcttt<br>gccctgggcgtcgataaaagggtcttgctcgcgcttgaggtgccatagcaccagcgggcccagcaggtgcccaagc<br>ggcactaccaggcccaacagcgcggagaggtggcagaacatcgcccattgccggatttcggcattgggcggggtga<br>tcgacaggttcgaatcgctcatgccccgcacctccttggccggtttcagtcagccagtgctgcctgctgcagctcgaaa<br>atctcgttcatgcccttctgcgccaaagcgagcatggcgttgaaatcttccggctggaacggtgcgccttcggcagtac<br>cctgcacttcgatgaaaccaccagcgctggtcatgaccacattcaggtcggtttcggctgcggagtcttccaggtagtc<br>caggtcgagcaccgcttcgccctgatacatgcctaccgacacggcagcgatcatgtgcttgagcgggttgccgccctt<br>gaggccgccgcgcttcttgatcactgccaaagcgtcgcacaaggcaaccatggcgccggtgatggacgcggtacgg<br>gtaccgccatcggcctggatcacgtcgcaatcgacgtacagggtgatgtcaccgagcttgctcatgtccagcgcggcg<br>cgcagcgaacggccgatcaggcgctggatttccagggtgcggccaccctgcttgccacggctggcttcgcgctggtt<br>acgttcgccggtggagcgcggcagcatgccatactcggcggtcagccagccttggccctggcctttgaggaagc | |
| $\Delta crc::P_{tac}$: $ligW2_{SYK\text{-}6}$ | pKMM073 | gttatggccctcctcctcccttcaagtcctaccatacaagcgggtttacccgcgaaagggccagatcaggcggcacaa<br>aaaaaggcgaccctctgaagaaggtcgcctttttcattcctgggccaggatcagatcccgtactgcgcccggtaggcct<br>cgacagccggcagatgctgtttcagctgtggatcgtcagccaggaactccagcacctgggtcagcgaaacaatgctg<br>accaccgggataccgaagtcgcgttccacttcctggatcgccgacagttcgccattgccgcgctcttcacggttcagcg<br>cgatcagtacaccagcggccttggcctgctgggcgttgatgatctgcatgacctcacgaatggcggtaccggcagtaa<br>tcacgtcgtcgatgatcagcacgtcaccggccagcggcgcgccgaccaggctgccaccttcgccatggtccttggctt<br>ccttgcggttgaagcaccatggcacgtccagctgatgctggtcggcaagggccacggcggtagtcgccgccaaagg<br>gatacccttgtaggccgggccgaacagcacatcgaacgggatcttgctatcgacgatggcggcggcgtagcaacgc<br>cccagctcagccagtgcggagccggtgttgaacaagccggcattgaagaaatacgggctggtacgccccgatttcag<br>ggtgaattcaccgaaacgcagtaccccgcgatcgatggcaaaacggataaagtcgcgctgatacggctgcatgaata<br>gtcccggacaccacggatttagctaaatgggttgagctcgggtatcatacacgcacgagatttatggggccatttgaatt<br>cGAGCTGTTGACAATTAATCATCGGCTCGTATAATGTGTGGAATTGTGA<br>GCGGATAACAATTTCACAC*TAAAGACCAAATAAATTAGGGAATCACAT*ATG<br>ACCCACGCACTCAACACCGGTGGCGACCGGGGATATCTCAGGATCGCC<br>ACGGAGGAAGCCTTCGCCACGCCCGCGCAGCTCGACGCTTATCTCAAG<br>CTCGTGCGCGAGGGACGGGCCGACAAGGCGACGACGTCGCTCTGGGGC<br>TTTTACGGCACGTCGCCTTCGGAGCGCTCGCAGTGGATCCGCGACCGGC<br>TCGTCGATCTCGACGACCTGCGCCTGCAGGCCATGGACGAGACCGGCA<br>TCGACGTCGCGATCCTCTCCATGACCTCGCCCGGCGGCCAGGTCTTTGA<br>GGCGGACGAGGCCAAGGCGCTGGTAAGCGAAGCCAATGACGTGCTCAA<br>GGCTGCGTGCGAGCGCTACCCCACGCGCTACTACGGCATGATCTCGATC<br>GTCCCGCAGGACCCGGCATGGTCCGTGGCGGAAATCCGTCGCGGCAAG<br>GAGGAGCTGGGCTTCCGGGGCGTGATGGTCAACAGCCACACCAAGGGC<br>CAGTATCTGGACGAGCCGCAGTTCGACCCCATTCTGCGCGCCTGCGCCG<br>AACAGGACCTGCCGCTCTACATCCACCCGCAGTCCCCGCCGGACGGCAT<br>GATCGCCGGCATGGTGGAAGCCGGGCTCGATGGCGCCATCTTCGGCTTC<br>GGCGTGGAAACGGGCTATCATCTGCTGCGCCTGCTCACCACCGGCGTGT<br>TCGACCGCTATCCGAACCTTCAGGTGGTCGTCGGCCATGGCGGCGAGGC<br>CATCCCGAACTGGCTGTTCCGCGTGGACTACATGCACAAGGCTGGCGTG<br>CGCTCGCAGCGCTACGAGCGCCTGAAGCCCCTGCAGCACGACATGTTCC<br>ACTACATGCGCAACAATGTGCTGGTGACCACCAGCGGCATGGCCAGCG<br>AGCCCACCATCAAGCTGTGCATGGAACAGCTCGGCGAGGACCGGGTGA<br>TGTATGCCATGGACTATCCCTATGAATATGTGGCGGACGAGGTGCGCGT<br>GCACGACAACCTCGCCATCCCCTTCGCCCAGAAGAAGAAGCTGATGCA<br>GACCAACGCCGAGCGAGTGTTCAAACTGTGA*agtcaaaagcctccggtcggaggctttt*<br>*gact*ttatttgatcggccgccagatcatcgggtagcgataaggcttgccctcattcgcccgcaccgccgcgatgatgat<br>caatatcatcactgccaccgtgagcatggcaaacagcaccagcccgataaacacgaacatcagcaggaagcagatg<br>aacccggcaatggtcacgctgatctgaaagttcagcgcctctttgccctgggcgtcgataaaagggtcttgctcgcgctt<br>gaggtgccatagcaccagcgggcccagcaggtgcccaagcggcactaccaggcccaacagcgcggagaggtgg<br>cagaacatcgcccattgccggatttcggcattgggcggggtgatcgacaggttcgaatcgctcatgccccgcacctcct<br>tggccggtttcagtcagccagtgctgcctgctgcagctcgaaaatctcgttcatgcccttctgcgccaaagcgagcatg<br>gcgttgaaatcttccggctggaacggtgcgccttcggcagtaccctgcacttcgatgaaaccaccagcgctggtcatga<br>ccacattcaggtcggtttcggctgcggagtcttccaggtagtccaggtcgagcaccgcttcgccctgatacatgcctac | Homology arms are shown in red. $P_{tac}$ promoter shown in BLACK UNDERLINE. ORFs for *ligW2* shown in BLUE. Synthetic ribosome binding sites for each gene shown in *ITALICIZED GRAY*. TonB terminator shown in *italicized black.* |

| | | | |
|---|---|---|---|
| | | cgacacggcagcgatcatgtgcttgagcgggttgccgcccttgaggccgccgcgcttcttgatcactgccaaagcgtc<br>gcacaaggcaaccatggcgccggtgatggacgcggtacgggtaccgccatcggcctggatcacgtcgcaatcgac<br>gtacagggtgatgtcaccgagcttgctcatgtccagcgcggcgc | |
| Δ*pcaHG*::$P_{tac}$:*aroY*$_{KP}$*:*ecdBD*$_{EC}$ | pKMM074 | gctggcttcggtcggcagacccgccagggtggtgtgttcctcacccggacccaccagggtctgcaggatggcgtggt<br>tgcgcatggtcacggccttcaccttgatcaccggcaggctcgggttggcctcaccgcagtagcctgggaattccggca<br>tggcgtgaccggtgttggtgtgctgatcttcgcgcacgcgcacgcctggcagcagttcgccctcgatgatgatctcggc<br>acgcgcgatcgccttctctttcacggccacgccctgcaccagttccaccggttgctgacgcagggcaccggccacac<br>ccagctcgttgtagccgaacggggtggttggcgcttcgaagcaggcaccgatgtagatggctgggtccaggcccatg<br>ttgatggtcaccggcagcggcttaccggcggcttcggccttcttgcggaacacctcgatgtggcgaccggcggccag<br>gaacatgctcagctcgtcgcgctcttgcacgcacaggcggtggatggtcacgtcggtcaggctggtatcttccgggtc<br>gctcgccagcaccaggcccaggcagaagaacggaccggcatcgatcggggtgttggttggggctggcagcagctt<br>gcgcaggtcgaagtccgggtcgtcggcgtagaacacctgttcttggcatggggcttggctggctggcaccaccactg<br>gggccaccgggttcttcacggcctggcccacgtgctgcgccagtttgcttggcacgcagcccagcagcagggcggc<br>acgttcacggctggcgtgcatgcccaccaggatgcggctgcctgggtagcccttcacgctgttgaacatcatggctgg<br>accggtgcgggtcggacgtttcacggtgccacccgcaccgatgtggcggtacacaccggccagttcggcgttcgggt<br>cgaccgggtggtcggtttcgatgtagtgacccggatggcgttgcagcagcgcgatcgcggagcgcaggtcgttgatc<br>gggttctgcat*AAAAAACCTCCTTAGGTTGAGTTTAGACTTCTTGGTCGCGGCCAAT*<br>*GGCC*gtgtgaaattgttatccgctcacaattccacacattatacgagccgatgattaattgtcaacagctcagatctgg<br>aattgtgagaacgcctgggcgtggcaggcggacttgcagtttccggcaggtcgcgcgtgatgaatattgaaatcgacc<br>aactcatacataaccaaatggtgatgtcttaacacctttgcctgacaccagccgctttcctgaactaatctttgtcttcttgttt<br>cacgctcggagagcattcatggaatggcgaaaaggccgacgcagcgacaacgtggtcgatgcccggggcgaaggt<br>ggcggtggcggcatgcgcttcggcggcggcaaggggctggggctcggggcgattctgctgatcgtcggcatcggct<br>ggctcaccgggcaggacccgctgcagattcttggccagctcaccggccagctggagcaacagcaaacggccccca<br>gtggtacaagcgccaaggcccccccggccaacgaccagcaggccgagttcgtcgcctcgattctgggcgacaccg<br>aggacacctggaaagccctgttcgcccaggccggcaaacaataccgtgaccccaaactggtcctgttcagcggccag<br>gtcaattcggcctgtggttttgcctcggcagcggtcgggccgttctattgcccggcggaccagcgggtgtacctggaca<br>tgtcgttcttccgcgaaatggaaacccgcttcgctgcagcgggcgacttcgcccaggcctacgtgatcgcccacgaaa<br>tcggtcaccatgtgcagacactgctgggcgtttcggccaaggtggatgccgcacgacgcgccggccagcgcatgga<br>gggcgacaacggcctgttggtgcgccaggagttgcaggctgactgcctggccggggtatgggcctaccaggcgca<br>gaaacgcctgaactggctggagccaggcgatgtcgaggaagcgctgaatgcggccaatgccattggcgacgaccg<br>cctgcagcagcaaggcc | Homology arms are shown in red. Synthetic ribosome binding site for *aroY* shown in *ITALICIZED GRAY*. |
| Δ*crc*::$P_{tac}$:<br>*ligW2*$_{SYK-6}$:*catA2* | pKMM075 | gttatggccctcctcctcccttcaagtcctaccatacaagcgggtttacccgcgaaagggccagatcaggcggcacaa<br>aaaaaggcgaccctctgaagaaggtcgcctttttcattcctgggccaggatcagatcccgtactgcgcccggtaggcct<br>cgacagccggcagatgctgtttcagctgtggatcgtcagccaggaactccagcacctgggtcagcgaaacaatgctg<br>accaccgggataccgaagtcgcgttccacttcctggatcgccgacagttcgccattgccgcgctcttcacggttcagcg<br>cgatcagtacaccagcggccttggcctgctgggcgttgatgatctgcatgacctcacgaatggcggtaccggcagtaa<br>tcacgtcgtcgatgatcagcacgtcaccggccagcggcgcgccgaccaggctgccaccttcgccatggtccttggctt<br>ccttgcggttgaagcaccatggcacgtccagctgatgctggtcggcaagggccacggcggtagtcgccgccaaagg<br>gatacccttgtaggccgggccgaacagcacatcgaacgggatcttgctatcgacgatggcggcggcgtagcaacgc<br>cccagctcagccagtgcggagccggtgttgaacaagccggcattgaagaaatacgggctggtacgccccgatttcag<br>ggtgaattcaccgaaacgcagtaccccgcgatcgatggcaaaacggataaagtcgcgctgatacggctgcatgaata<br>gtcccggacaccacggatttagctaaatgggttgagctcgggtatcatacacgcacgagatttatggggccatttgaatt<br>cGAGCTGTTGACAATTAATCATCGGCTCGTATAATGTGTGGAATTGTGA<br>GCGGATAACAATTTCACAC*TAAAGACCAAATAAATTAGGGAATCACAT*ATG<br>ACCCACGCACTCAACACCGGTGGCGACCGGGGATATCTCAGGATCGCC<br>ACGGAGGAAGCCTTCGCCACGCCCGCGCAGCTCGACGCTTATCTCAAG<br>CTCGTGCGCGAGGGACGGGCCGACAAGGCGACGACGTCGCTCTGGGGC<br>TTTTACGGCACGTCGCCTTCGGAGCGCTCGCAGTGGATCCGCGACCGGC<br>TCGTCGATCTCGACGACCTGCGCCTGCAGGCCATGGACGAGACCGGCA<br>TCGACGTCGCGATCCTCTCCATGACCTCGCCCGGCGGCCAGGTCTTTGA<br>GGCGGACGAGGCCAAGGCGCTGGTAAGCGAAGCCAATGACGTGCTCAA<br>GGCTGCGTGCGAGCGCTACCCCACGCGCTACTACGGCATGATCTCGATC<br>GTCCCGCAGGACCCGGCATGGTCCGTGGCGGAAATCCGTCGCGGCAAG<br>GAGGAGCTGGGCTTCCGGGGCGTGATGGTCAACAGCCACACCAAGGGC<br>CAGTATCTGGACGAGCCGCAGTTCGACCCCATTCTGCGCGCCTGCGCCG<br>AACAGGACCTGCCGCTCTACATCCACCCGCAGTCCCCGCCGGACGGCAT<br>GATCGCCGGCATGGTGGAAGCCGGGCTCGATGGCGCCATCTTCGGCTTC<br>GGCGTGGAAACGGGCTATCATCTGCTGCGCCTGCTCACCACCGGCGTGT<br>TCGACCGCTATCCGAACCTTCAGGTGGTCGTCGGCCATGGCGGCGAGGC<br>CATCCCGAACTGGCTGTTCCGCGTGGACTACATGCACAAGGCTGGCGTG<br>CGCTCGCAGCGCTACGAGCGCCTGAAGCCCCTGCAGCACGACATGTTCC<br>ACTACATGCGCAACAATGTGCTGGTGACCACCAGCGGCATGGCCAGCG<br>AGCCCACCATCAAGCTGTGCATGGAACAGCTCGGCGAGGACCGGGTGA<br>TGTATGCCATGGACTATCCCTATGAATATGTGGCGGACGAGGTGCGCGT<br>GCACGACAACCTCGCCATCCCCTTCGCCCAGAAGAAGAAGCTGATGCA<br>GACCAACGCCGAGCGAGTGTTCAAACTGTGA*GCTCCCCAATAGCGATCG*<br>*AGACTTTTAT*ATGACCGTGAACATTTCCCATACTGCCGAGGTACAGCAGT<br>TCTTCGAGCAGGCCGCAGGCTTTTGTAATGCGGCCGGCAACCCACGCCT<br>CAAACGCATCGTGCAGCGCCTGCTGCAGGATACCGCGCGGCTGATCGA<br>AGACCTGGACATCAGCGAAGACGAGTTCTGGCACGCCGTCGATTACCT | Homology arms are shown in red. $P_{tac}$ promoter shown in BLACK UNDERLINE. ORFs for *ligW2*$_{SYK-6}$ *catA2* are shown in BLUE and ORANGE, in that order. Synthetic ribosome binding sites for each gene shown in *ITALICIZED GRAY*. TonB terminator shown in *italicized black*. |

| | | | |
|---|---|---|---|
| | | CAACCGCCTGGGCGGTCGCGGCGAAGCCGGGTTGCTGGTGGCGGGGCT<br>GGGCATCGAACACTTCCTCGACCTGCTGCAGGATGCCAAGGACCAGGA<br>GGCAGGGCGCGTTGGCGGCACCCCACGCACCATCGAAGGCCCGTTGTA<br>CGTGGCTGGCGCACCGATTGCCCAAGGTGAAGTGCGCATGGACGACGG<br>CAGCGAGGAGGGCGTGGCCACGGTGATGTTCCTGGAAGGCCAGGTGCT<br>GGACCCGCACGGACGCCCGCTGCCGGGTGCCACGGTCGACCTGTGGCA<br>TGCCAATACCCGTGGTACCTACTCGTTCTTCGACCAAAGCCAGTCGGCG<br>TACAACCTGCGTCGGCGCATCGTTACCGATGCCCAGGGGCGCTACCGCG<br>CGCGCTCCATCGTGCCATCGGGCTATGGCTGCGACCCGCAGGGGCCAA<br>CCCAGGAATGCCTGGACCTGCTGGGCCGTCATGGCCAGCGCCCGGCGC<br>ACGTGCACTTCTTTATCTCGGCCCCAGGGTACCGGCACCTGACCACGCA<br>GATAAACCTGTCGGGGGACAAGTACCTGTGGGATGACTTTGCCTATGCC<br>ACACGGGATGGGCTGGTCGGGGAGGTGGTGTTCGTCGAAGGGCCGGAT<br>GGTCGGCATGCCGAGCTGAAGTTCGACTTCCAGTTGCAGCAGGCCCAG<br>GGCGGTGCCGATGAGCAGCGCAGCGGGCGGCCGCGAGCTTTGCAGGAG<br>GCCTGA*agtcaaaagcctccgaccggaggcttttttac*taaggccattggggctgcattgcagccccaatggttat<br>ttgatcggccgccagatcatcgggtagcgataaggcttgccctcattcgcccgcaccgccgcgatgatgatcaatatca<br>tcactgccaccgtgagcatggcaaacagcaccagcccgataaacacgaacatcagcaggaagcagatgaacccgg<br>caatggtcacgctgatctgaaagttcagcgcctctttgccctgggcgtcgataaaagggtcttgctcgcgcttgaggtgc<br>catagcaccagcgggcccagcaggtgcccaagcggcactaccaggcccaacagcgcggagaggtggcagaacat<br>cgcccattgccggatttcggcattgggcggggtgatcgacaggttcgaatcgctcatgccccgcacctccttggccggt<br>ttcagtcagccagtgctgcctgctgcagctcgaaaatctcgttcatgcccttctgcgccaaagcgagcatggcgttgaaa<br>tcttccggctggaacggtgcgccttcggcagtaccctgcacttcgatgaaaccaccagcgctggtcatgaccacattca<br>ggtcggtttcggctgcggagtcttccaggtagtccaggtcgagcaccgcttcgccctgatacatgcctaccgacacgg<br>cagcgatcatgtgcttgagcgggttgccgcccttgaggccgccgcgcttcttgatcactgccaaagcgtcgcacaagg<br>caaccatggcgccggtgatggacgcggtacgggtaccgccatcggcctggatcacgtcgcaatcgacgtacagggt<br>gatgtcaccgagcttgctcatgtccagcgcggc | |
| *PP_5042::P$_{tac}$:marK*$^{E16K}$ *Novosphingobium aromaticivorans* DSM12444 | pKMM111 | ggcttcgaaccagtcatggtcgaaggcgtgctcggggtccttgccgactgcataggtcagcttgttcagtacagcagc<br>gcgaaaatccgcgacttcggcgtcacgtgctttaggttcctgggacatgcggcatcctcgggcaagttgacgaaagcg<br>gagggagagtgttgagactagacccttcgacagagagcgacagttacggttcgcagttttcatgcccactctgcgattc<br>cggcaaaaggttgttcataaattgaacaattccggtatcatcgcgcgccccaagaccgtgatccaccctataaagatga<br>aaacgaccctgatcgccgcggccgaagtcgaccgcctggagacctggcagcgctacaccagcaacatgtgccatgg<br>ctgccattcgacctgctgcaccctgccggtggaggtgaagatcaaggatctgatccgtatcggcgtggtcgacgagttc<br>gaaaaagacgaaccacccaagaacgtggccaagcgcctgcaaaaagaaggcatcatcgagcgcttcaaccagaag<br>tccgggatcttcaccctgacccggatgagcaacgacgactgcatgtacctggatcgtaaaagccggctgtgcaccattt<br>atgacaagcgcccggatacctgccgcaaccaccccaaggtcgggccacggccggggtattgcgcctacaagccca<br>aggtggttgggcgttgagtcttgcgccggggcataaatcgagcgccgcgcgggcggcgctcgattctgtggcgatgc<br>GAGCTGTTGACAATTAATCATCGGCTCGTATAATGTGTGGAATTGTGAG<br>CGGATAACAATTTCACAC*TAAAACTACATTAGGAGCCATTA*ATGACCGACA<br>AGCCGAGCGTCCTGTTCATCTGCACCCAGGACACCAAAGAGGAAGAGG<br>CTCGCTTCACCCGTGCCGCCCTGGAAGCGGCCGGCGTCGAAGTCGTCCA<br>CTTGGACCCAAGCGTCCGCCGCAGCCTCGGCGGGGCCGAAATCTCGCCT<br>GAGATGGTCGCCCAGGCAGGGGGGATGACCATTGAGGAGGTGCGCGCC<br>CTGGGCCACGAGGGCAAGTGCCAGGACGCCATGATCCGCGGCGCCATT<br>GCGGCTGCCCACGAGTGGGATGCTCGTCACCCGGTCAGCGGCATCTTGG<br>CCGTCGGCGGCTCGATGGGGAGCGCGCTCGCGGGCGCGCTGATGCAGA<br>GCTTCCCGTACGGCCTCCCGAAGCTGATCGTCTCGACCATGGCCTCGGG<br>TTTCACCAAGCCGTATATGGGCGTCAAGGACATCGCGATGATGAACGC<br>AGTGACGGACATCTCGGGCATCAACACCATCTCGCGCGACGTCTTCCGT<br>AACGCTGCCAACGCCGTGGCGGGCATGGCCAAGGGGTACGACCGCGAC<br>AAGGGGCCGGAGAAGCCCCTGGTCCTGATTACCACGCTGGGCACCACC<br>GAGACGTCCGTGAAGCGCATCCGCCAGGCACTCGAATCCGACGGTTGC<br>GAGGTCATGGTCTTCCACAGCTCCGGCGCGGGGGGTCCCACCCTGGAC<br>GGCCTGGCGGCCGATAAAGATGTCGCGCTGGTGTTGGACTTGAGCCCC<br>ACCGAGATCTTGGACCACTTGTTCGGCGGCCTGGCGGACGCGGGCCCC<br>GACCGCGGCCGCGCCGCGCTCCGTAAGGGCATTCCTACCATTCTCGCCC<br>CGGGCAACGCCGATTTCATTATCGGTGGCCCGATCGACGCGGCCGAGG<br>CACAGTTTCCCGGCCGTCGCTACCATCAGCATAACCCCCAGTTGACCGC<br>CGTGCGCACCAACGTGGCCGACCTGCGTAAGCTGGCTGACCACCTGGC<br>GGCGAACGTCCGCGAAGCGAAGGGCCCGGTGCGCGTCTTCACGCCACT<br>CAAGGGCTTCAGCTCGCACGACTCGGAGACGGGCCACCTGCTGGACCT<br>CTCCGTCCCGGGCCCCTTCGCCGAGTATTTGGCCTCGGTGATGCCGGGC<br>CATGTGCCCGTGACGGCCGTGGATGCTCACTTCAACGATGAGGCCTTCA<br>GCAGCGCTGTGATTGCCGCAGCCCGGGAAATGCTCGCCGCCAAGAACT<br>AAcgactcgattatcaacgggtgtcttgcagttctcatcgctactggtactagt*agtcaaaagcctccgaccggaggc<br>ttttgact*cgagctgggcatgaagcactgaacccaccgctcccatttcagccagcggggaagctccggcaagcaccc<br>ccctgatccgctggtggatgctcaggataagtctgcggcaaaacgctgaccgggctggtgacaacgaaaccatcagc<br>aaaccgcgcctctacgaaccgcgcctgccagccttgcgccggcggcgcccagtcaaggctgaacgcctcactggg<br>gatgatcgtctgcgatgtgtagcgtaccccgcaggcatggcggaagtcccgatccaaggggttagccgcagtccata<br>ccttcactgctacaggcatttccgaaagccgtaccagcacatgcgcagctccttcgcccggagccacttgcacgcgag | Homology arms are shown in red. P$_{tac}$ promoter shown in BLACK UNDERLINE. ORFs for *marK*$^{E16K}$ codon optimized for *Pseudomonas putida* shown in BLUE. Synthetic ribosome binding sites for each gene shown in *ITALICIZED GRAY*. TonB terminator shown in *italicized black*. |

| | | | |
|---|---|---|---|
| | | gcatcggcaggctggcctggatacgctgaaccatcggcacgagggtagtgacaacatgcgcacgcacgcccccatg<br>gtccgagttcggcagcaccctgaggctggtctggccaggcaggcgctgcaggtagtccgtggcgtgatcaggggcg<br>aagaaatcatccccgctggcggacaccaggtatttcggtatcttcaggcgagcacccggctcatccaggtactgcatc<br>gggtccatcagtgccatcaaccgagcgaaggccgggctgcccaactgctgcaatacctccgcagcctgataggacc<br>aaagcgccaatggccatgaaccaccgtaccgcttgcgcaagccctccagcatgctggaagtgtcggcaacatcgatg<br>acgctggg | |
| $P_{tac}$:*mdlC*$_{PP}$ | pKMM118 | CTATGGAGGTCAGGTATGATTACTATTGACAATTAATCATCGGCTCGTA<br>TAATGTGATCAGACCTGGAATTGTGAGCGGATAACAATT*CTTAAGATTAA*<br>*CTCACACAGGAGGGTATCAT*ATGGCCTCGGTCCATGGCACCACCTACGAG<br>CTCCTGCGGCGCCAGGGCATCGACACCGTGTTCGGCAATCCGGGCAGC<br>AACGAGTTGCCCTTTCTCAAAGACTTCCCCGAAGATTTTCGCTACATCC<br>TGGCCTTGCAGGAGGCGTGCGTGGTGGGTATCGCCGACGGTTACGCCC<br>AGGCGTCGCGCAAGCCGGCCTTTATCAACTTGCACTCCGCCGCCGGCAC<br>CGGCAATGCCATGGGCGCCCTGTCGAATGCCTGGAACAGCCACAGCCC<br>CCTGATCGTCACGGCGGGCCAGCAGACCCGCGCCATGATTGGCGTGGA<br>GGCGCTCCTGACGAACGTGGATGCCGCCAACCTCCCGCGCCCGCTGGTG<br>AAGTGGTCCTACGAACCGGCGAGCGCAGCAGAAGTCCCTCACGCCATG<br>TCCCGTGCGATTCATATGGCCTCCATGGCCCCGCAGGGCCCGGTGTACT<br>TGAGCGTCCCCTATGACGATTGGGATAAGGACGCGGACCCGCAGAGCC<br>ACCATTTGTTCGACCGCCACGTGTCCTCGTCGGTGCGGCTGAACGATCA<br>GGACCTGGACATCCTGGTCAAAGCACTGAACTCGGCGAGCAACCCAGC<br>GATTGTCTTGGGTCCCGATGTGGATGCAGCGAACGCTAACGCGGATTGC<br>GTGATGCTGGCTGAACGCCTGAAAGCGCCGGTGTGGGTGGCCCCCAGC<br>GCGCCGCGTTGTCCGTTCCCCACCCGGCATCCATGCTTTCGCGGTTTGAT<br>GCCGGCGGGCATTGCGGCCATCTCCCAGCTGTTGGAAGGCCACGACGT<br>CGTGCTGGTGATTGGCGCACCGGTCTTCCGTTATCATCAGTATGATCCT<br>GGTCAGTACCTCAAGCCGGGCACCCGCCTGATCAGCGTGACCTGCGAC<br>CCGCTGGAGGCCGCCCGGGCCCCGATGGGTGATGCCATCGTCGCGGAT<br>ATTGGCGCCATGGCGAGCGCCCTGGCAAATCTGGTGGAAGAGTCCTCG<br>CGCCAGCTGCCTACCGCCGCCCCAGAGCCGGCAAAGGTCGACCAAGAC<br>GCCGGCCGCCTGCACCCGGAAACCGTCTTCGACACGCTGAACGACATG<br>GCCCCCGAGAACGCGATTTACCTGAATGAGTCGACCTCGACGACCGCC<br>CAGATGTGGCAGCGCCTGAACATGCGCAACCCGGGCTCGTACTACTTCT<br>GCGCAGCAGGTGGCCTCGGCTTCGCCCTGCCGGCGGCCATTGGCGTGCA<br>GCTCGCCGAACCGGAGCGCCAGGTGATCGCTGTCATCGGCGACGGCAG<br>CGCCAATTACTCGATCAGCGCGCTGTGGACCGCCGCGCAGTACAACATC<br>CCGACCATCTTCGTGATCATGAACAATGGCACCTACGGCGCACTCCGCT<br>GGTTCGCGGGCGTGCTCGAGGCCGAAAACGTGCCGGGCCTGGACGTGC<br>CCGGTATCGATTTCCGCGCGCTGGCCAAGGGTTATGGCGTGCAGGCCCT<br>CAAGGCCGACAACCTCGAACAGTTGAAAGGTTCGCTGCAAGAAGCCCT<br>CAGCGCCAAAGGCCCCGTCCTGATCGAAGTGAGCACGGTCAGCCCGGT<br>CAAGTAA | $P_{JE111111}$ promoter shown in BLACK UNDERLINE. ORF for *mdlC*$_{PP}$ is shown in BLUE. Synthetic ribosome binding site JER01 shown in *ITALICIZED GRAY*. |
| $P_{tac}$:*dpgB*$_{ST\text{-}201}$ | pKMM119 | CTATGGAGGTCAGGTATGATTACTATTGACAATTAATCATCGGCTCGTA<br>TAATGTGATCAGACCTGGAATTGTGAGCGGATAACAATT*CTTAAGATTAA*<br>*CTCACACAGGAGGGTATCAT*ATGGCCTCGGTCCACTCCATCACGTACGAA<br>CTGCTGCGTCGCCAGGGGATCGACACCGTGTTCGGCAATCCTGGCAGCA<br>ACGAATTGCCGTTCCTGAAGGATTTCCCCGAAGACTTTCGTTACATTCT<br>GGCCCTGCAGGAAGCATGCGTGGTGGGCATCGCAGACGGCTACGCCCA<br>GGCCTCGCGTAAGCCGGCGTTCATCAATCTCCACTCCGCCGCCGGTACG<br>GGTAACGCTATGGGCGCGATGAGCAACGCCTGGAACTGCCATTCGCCG<br>TTGATCGTCACCGCGGGGCAACAGAACCGCGCGATGATCGGCGTCGAG<br>GCGTTGCTCACCAACGTGGACGCAGCAAGCCTCCCGCGTCCGCTGGTCA<br>AATGGAGCTACGAGCCAGCGTCGGCAGCCGAAGTCCCGCACGCCATGA<br>GCCGCGCGATCCATATGGCCTCGATGGCTCCCCGCGGCCCGGTGTACCT<br>CTCGGTGCCCTATGACGATTGGGACAAGGAGGCGGATCCGCAAAGCCA<br>CCACCTGTATGATCGGAGCGTCAACTCCGCCGTGCGGTTGAACGATCAG<br>GACTTGGAAGTGCTGGTCGAAGCCTTGAACAGCGCGTCGAACCCCGCG<br>ATCGTGCTGGGCCCTGACGTGGATTCCGCGAATGCCAACGCCGACTGCG<br>TCACGCTGGCTGAGCGCCTGAAGGCCCCTGTGTGGGTGGCTCCGAGCGC<br>CCCCCGGTGCCCGTTCCCTACCCGTCACCCGTGTTTTCGCGGCCTGATGC<br>CCGCCGGCATCGCCGCCATTTCGCAGCTCCTCGAGGGTCACGACGTGGT<br>CCTGGTGATTGGTGCGCCAGTGTTCCGCTATCATCAGTACGACCCTGGC<br>CAGTATTTGAAACCGGGCACCCGGCTGATCTCGATTACGTGCGACCCAT<br>TGGAAGCGGCACGGGCTCCGATGGGGGATGCCATCGTGGCCGACATCG<br>GCACCATGACGGCCGCTCTGGCGAGCCGCATTGGCGAAAGCGAGCGCC<br>AGCTGCCCGCCGTGTTGCCCAGCCCGGAACGTGTCAACCAGGACGCCG<br>GTCGCCTGCGGCCAGAGACCGTGTTTGACACCCTCAATGAAATGGCCCC<br>GGAAGACGCCATTTACCTGAACGAGTCCACCAGCACCACCGCACAGAT<br>GTGGCAGCGCCTGAACATGCGGAATCCGGGCAGCTACTACTTCTGCGC<br>AGCCGGCGGTCTGGGCTTTGCCCTGCCCGCCGCCATCGGCGTCCAGCTG | $P_{JE111111}$ promoter shown in BLACK UNDERLINE. ORF for *dpgB*$_{ST\text{-}201}$ codon optimized for *Pseudomonas putida* is shown in BLUE. Synthetic ribosome binding site JER01 shown in *ITALICIZED GRAY*. |

| | | | |
|---|---|---|---|
| | | GCGGAGCCAGATCGCCAGGTGATCGCGGTGATCGGCGACGGTTCGGCC<br>AATTATAGCATTAGCGCCCTGTGGACCGCCGCACATTACAACATTCCTG<br>CGATCTTCCTGATCATGAACAACGGCACCTACGGGGCCTTGCGGTGGTT<br>CGCCGGTGTGCTCGAGGCGGAGAATGTGCCGGGTCTCGATGTCCCAGG<br>TATTGACTTCTGTGCAATTGCCAAGGGCTACGGCATCCCTGCTCTGAAA<br>GCCGACAACCTGGAACAGCTGAAAGGCTCCATCCACGAAGCCCTGAGC<br>GCCAAGGGCCCAGTGCTGATCGAGGTCTCCACCGTGAGCTTGTAA | |
| $P_{tac}$:*EC844_1132*$_{AC}$ | pKMM120 | CTATGGAGGTCAGGTATGATTACTATTGACAATTAATCATCGGCTCGTA<br>TAATGTGATCAGACCTGGAATTGTGAGCGGATAACAATT*CTTAAGATTAA*<br>*CTCACACAGGAGGGTATCAT*ATGAAAACCGTGCATCAGTATAGCTACGAT<br>ATCCTGCGCAAGAACAACATCAAGACCATCTTCGGTAACCCTGGCAGC<br>AACGAACTGCCCTTCCTGAAAAACTTCCCGGACGACTTTCAATATATTC<br>TGGCGCTGCATGAGGGCGCCGCCGTGAGCATGGCAGACGGTTACGCTG<br>CCGCCAGCGGCACCGTCGCGTTTGTGAATCTGCACAGCGCCGCCGGCAC<br>CGGTAATGCCATGGGGGCGCTGTCCAACGCCTGCACGAGCCACACGGC<br>CATGGTCATCACCAGCGGGCAGCAGACCCGCCTGATGCTGGGCGTGGA<br>GGCCCTGCTGACCAACGTGAACGCCGTCCAGCTGCCGCAGCCCTTGGTC<br>AAGTGGTCGCATGAGCCGGCCTGCGCCGCGGAAGTGCCGCATGCTATG<br>AGCCGTGCTATCCATATTGCGAAGAGCGAACCCGCGGGCCCTGTGTACC<br>TGAGCGTCCCGTATAACGACTGGGATGTGGAAGTCAGCACCGAGAACG<br>ATCACCTCCTGCAGCGCGCCGTGGACAGCGGCAACTGTCTGTCCATCGA<br>ACATCTGTCCGTCATCCTCGGCCACATCCACGCGGCCAACAATATCGCG<br>CTGGTGCTGGGCCCGGATGTGGAGCGCCTGCACGCGGTGCACGACGCC<br>GTGTACCTCGCAGAGAAGCTGAACGTGCCTGTCTGGGGTGCCCCCTCGC<br>CCTCCCGCTGCCCCTTCCCGAACCGCCACCCGTACTTCCAGGGTTTCTTG<br>CCGGCTTCCATCGCGGGCATCTCGCAGATCCTCGCAGACTTCGATCTGA<br>TTTTGGTGTTCGGCGCCCCTGTGTTCCGTTACCATCAGTACGAACCAGG<br>CTGCTACATTGCGGAGAAGTGTAAGCTGATCAGCCTGAGCTGTGACATT<br>CAGGAGGCCGCGCGTGCCCCGATGGGTCTCAGCTATGTGTGCGACATC<br>AAGGACGCCCTGCGGCGGATCAACAACGCCCTGTCCTTGCGCCTGAAC<br>AAGGTGAACACCCTGAAGCCACTGCAGGCAGCCGAAGCAAGCGTGGAC<br>GGTTACATTAAGCCGGAACGCCTGTTTGACCTGATCGATGCGCTCGCCC<br>CTGAGAACGTCATCTACACCAATGAAGCGACCGCCACCAACAGCATTTT<br>CTGGCAGCGCATCGGCCTGAAAGGCCAGGGCTCGTATTATTTCGCCGCA<br>GCTGGTGGGCTCGGCTACGCAATGCCGGCGGCAGTGGGTATCCAACTG<br>GCCAAGCCGAAATGTCGGGTGGTGGCAATCATTGGTGACGGCTCGGCG<br>AATTACAACATCACCGCCCTGTGGACCGCCGCACAGTACAAACTCGCTG<br>TCGTGTTCATCATTCTGAAAAACGGCTCGTATGCTGCGCTGAAGTGGTT<br>CGCCGAGGTGCTCAAGGCGAGCCACGTGCCAGGGATGGATATCCCGGG<br>GATCGACTTTGTGCAGCTGGCGCGTGGCTACGGCGTCCAGGGCATCCGC<br>GCGAACAACGATGAGGACTTTATCAACGCCTACAAACATGCGCTCAGC<br>TCGCATACCCCGACCCTGATCGAAGTGACGACCGTGCCGCCCCACATTC<br>CCAACCCTAAGCCGTAA | $P_{JE111111}$ promoter shown in BLACK UNDERLINE. ORF for *EC844_1132*$_{AC}$ codon optimized for *Pseudomonas putida* is shown in BLUE. Synthetic ribosome binding site JER01 shown in *ITALICIZED GRAY*. |
| Δ*hsdRM*::$P_{tac}$:*mdlC*$_{PP}$ | pKMM123 | gcatttacgagctcatttcttgccttgggtctgtaagcgtcgctaagccgtatgacgcagtcctgggtcttcaaaataatttc<br>agcacgctcctttgagcccagctctttggaatccatcatcgttgcaagttgggtactcaacgagtccttcatcgatgcact<br>gaaatctacccctttagacttggtcgcatccgcatactgaaaagcgcctttgaatgcggcggttacatctgccgaatacc<br>caaggccacatgccttaatcgcctgatcaactggtgcaacagatcgatagggagaaaggcctgaacaagctgacaag<br>aacacagcggaaagcaaccagatatgactacgcattaacaatttctccacccccaaataatggcactaagccactatac<br>cacttattcgacgtataggtttggccagagtggcaggcgccacttgcgagcatccacccaacgcgaaaatgtcattaat<br>agtagggctgagctaactgagcgctgacagctgggaacaccttgggcatgcggcttggcttcatgaacaccgatgag<br>cttggtattgcaacgtactggccatgcggctgctgaatcatcatttagaaaagccgtgaggtttccactgccatcagcgc<br>ataggaaggcccgacgaccgggcctgagagccccccgacgatagtggcagcatgcctctagctattttgttaaattcg<br>gcaacacgccatagggacatagccagcaCTATGGAGGTCAGGTATGATTACTATTGAC<br>AATTAATCATCGGCTCGTATAATGTGATCAGACCTGGAATTGTGAGCGG<br>ATAACAATT*CTTAAGATTAACTCACACAGGAGGGTATCAT*ATGGCCTCGGT<br>CCATGGCACCACCTACGAGCTCCTGCGGCGCCAGGGCATCGACACCGT<br>GTTCGGCAATCCGGGCAGCAACGAGTTGCCCTTTCTCAAAGACTTCCCC<br>GAAGATTTTCGCTACATCCTGGCCTTGCAGGAGGCGTGCGTGGTGGGTA<br>TCGCCGACGGTTACGCCCAGGCGTCGCGCAAGCCGGCCTTTATCAACTT<br>GCACTCCGCCGCCGGCACCGGCAATGCCATGGGCGCCCTGTCGAATGC<br>CTGGAACAGCCACAGCCCCCTGATCGTCACGGCGGGCCAGCAGACCCG<br>CGCCATGATTGGCGTGGAGGCGCTCCTGACGAACGTGGATGCCGCCAA<br>CCTCCCGCGCCCGCTGGTGAAGTGGTCCTACGAACCGGCGAGCGCAGC<br>AGAAGTCCCTCACGCCATGTCCCGTGCGATTCATATGGCCTCCATGGCC<br>CCGCAGGGCCCGGTGTACTTGAGCGTCCCCTATGACGATTGGGATAAG<br>GACGCGGACCCGCAGAGCCACCATTTGTTCGACCGCCACGTGTCCTCGT<br>CGGTGCGGCTGAACGATCAGGACCTGGACATCCTGGTCAAAGCACTGA<br>ACTCGGCGAGCAACCCAGCGATTGTCTTGGGTCCCGATGTGGATGCAGC<br>GAACGCTAACGCGGATTGCGTGATGCTGGCTGAACGCCTGAAAGCGCC<br>GGTGTGGGTGGCCCCCAGCGCGCCGCGTTGTCCGTTCCCCACCCGGCAT | Homology arms are shown in red. $P_{JE111111}$ promoter shown in BLACK UNDERLINE. ORF for *mdlC*$_{PP}$ is shown in BLUE. Synthetic ribosome binding site JER01 shown in *ITALICIZED GRAY*. The L3S1P13 and ECK120010850 terminators shown in *italicized black* in that order. |

| | | |
|---|---|---|
| | | CCATGCTTTCGCGGTTTGATGCCGGCGGGCATTGCGGCCATCTCCCAGC<br>TGTTGGAAGGCCACGACGTCGTGCTGGTGATTGGCGCACCGGTCTTCCG<br>TTATCATCAGTATGATCCTGGTCAGTACCTCAAGCCGGGCACCCGCCTG<br>ATCAGCGTGACCTGCGACCCGCTGGAGGCCGCCCGGGCCCCGATGGGT<br>GATGCCATCGTCGCGGATATTGGCGCCATGGCGAGCGCCCTGGCAAAT<br>CTGGTGGAAGAGTCCTCGCGCCAGCTGCCTACCGCCGCCCCAGAGCCG<br>GCAAAGGTCGACCAAGACGCCGGCCGCCTGCACCCGGAAACCGTCTTC<br>GACACGCTGAACGACATGGCCCCCGAGAACGCGATTTACCTGAATGAG<br>TCGACCTCGACGACCGCCCAGATGTGGCAGCGCCTGAACATGCGCAAC<br>CCGGGCTCGTACTACTTCTGCGCAGCAGGTGGCCTCGGCTTCGCCCTGC<br>CGGCGGCCATTGGCGTGCAGCTCGCCGAACCGGAGCGCCAGGTGATCG<br>CTGTCATCGGCGACGGCAGCGCCAATTACTCGATCAGCGCGCTGTGGAC<br>CGCCGCGCAGTACAACATCCCGACCATCTTCGTGATCATGAACAATGGC<br>ACCTACGGCGCACTCCGCTGGTTCGCGGGCGTGCTCGAGGCCGAAAAC<br>GTGCCGGGCCTGGACGTGCCCGGTATCGATTTCCGCGCGCTGGCCAAGG<br>GTTATGGCGTGCAGGCCCTCAAGGCCGACAACCTCGAACAGTTGAAAG<br>GTTCGCTGCAAGAAGCCCTCAGCGCCAAAGGCCCCGTCCTGATCGAAG<br>TGAGCACGGTCAGCCCGGTCAAGTAAaggttctgttaagtaactgaacccaatgtcgttagtga<br>cgcttacctcttaagaggtcactgacctaacattcaggtagaagacaactggtctagactcgag*gacgaacaataagg*<br>*cctccctaacggggggcctttttattgataacaaaa*atccaca*aggaaaaattaaaggggagataaaatcccccc*<br>*tttttggttaact*gcggccgcgcacaggtgtagtggaacctgcacgaatacagcaacacaccctcggcctttgtggtgc<br>gcagcgtgtttgaagacgcctacaactacatgaaatccggccagctgctgcgccaggtgatcaacaagattcaggaag<br>gcgtggacttcaacagggcccaggaacgccacgagttcggcaacctctatgaacaattgctgcgcgacctgcagaac<br>gccggcaacgccggtgagttctacacacctcgaccagtcaccgaatttatggtgcgcatggttgatcccaagctggctg<br>aaaaggtcatggacccggcctgcggcaccggcggctttctcacctgcgccatcgagcacaagcgcagacgctatgta<br>aaaaccgccgaagacgaacgcaccttgcaggccagcatttttggcgtggagaaaaaaccgctgccgcacctgctgg<br>ccaccaccaacatgatcctgcatggcatcgaagtgcccagccagatccgtcacgacaacaccctgagcaaaccgctg<br>atcagctggggcccaagcgagcgcgtgcattgtatcgtcgccaacccgccgttcggcggcatggaagaagacggtat<br>cgaaacaaattttcctgccgctttccgcacccgggaaaccgccgatttgttcttggtattgattatgcagctgctcaaagat<br>ggtggccgcgccgcagtggtactgcccgatggcttcctttttggcgaaggcatcaaaagcc |

**Table S6. Retention time, absorbance values, and UV responses from calibration curves of each compound as described in section 4.4.**

| Compound ID | Compound Name | Retention time (min) | $\lambda_{max}$ (nm) | UV Response/ mM |
|---|---|---|---|---|
| VGA | vanillyl glyoxylic acid | 0.423 | 230, 283, 310 | 684000 |
| 4HBAld | 4-hydroxybenzaldehyde | 1.123 | 255 | 263000 |
| 4HBA | 4-hydroxybenzoic acid | 1.701 | 284 | 1150000 |
| VA | vanillic acid | 1.792 | 260, 292 | 370000 |
| 5CVA | 5-carboxyvanillic acid | 2.067 | 227, 268, 308 | 484000 |
| 5-CVn | 5-carboxyvanillin | 2.464 | 231, 285, 317 | 748000 |
| Vn | vanillin | 2.620 | 279, 309 | 856000 |
| AV | acetovanillone | 3.688 | 275, 304 | 765000 |
| 5FVn | 5-formylvanillin | 3.925 | 247, 336 | 630000 |
| 1,4-DMB | 1,4-dimethoxybenzene | 7.224 | 288 | 165000 |

**Table S7. UPLC gradient profile for aromatics quantitation (section 4.4).**

| Time (mins) | Flow (mL/min) | %A | %B |
|---|---|---|---|
| Initial | 0.750 | 97.0 | 3.0 |
| 0.30 | 0.750 | 97.0 | 3.0 |
| 6.00 | 0.750 | 87.5 | 12.5 |
| 7.00 | 0.750 | 70.0 | 30.0 |
| 7.80 | 0.750 | 65.0 | 35.0 |
| 8.70 | 0.750 | 10.0 | 90.0 |
| 9.00 | 0.750 | 10.0 | 90.0 |
| 10.00 | 0.750 | 97.0 | 3.0 |
| 10.70 | 0.750 | 97.0 | 3.0 |
